# Universal orthologs infer deep phylogenies and improve genome quality assessments

**DOI:** 10.1101/2025.02.17.638702

**Authors:** Md Nafis Ul Alam, Cristian Román-Palacios, Dario Copetti, Rod A. Wing

## Abstract

Universal single-copy orthologs are the most conserved components of genomes. Although they are routinely used for studying evolutionary histories and assessing new assemblies, current methods do not incorporate information from available genomic data. Here, we first determine the influence of evolutionary history on universal gene content in plants, fungi and animals. We find that across 11,098 genomes comprising 2,606 taxonomic groups, 215 groups significantly vary from their respective lineages in terms of their BUSCO (Benchmarking Universal Single Copy Orthologs) completeness. Additionally, 169 groups display an elevated complement of duplicated orthologs, likely as an artifact of whole genome duplication events. Secondly, we investigate the extent of taxonomic congruence in BUSCO-derived whole-genome phylogenies. For 275 suitable families out of 543 tested, sites evolving at higher rates produce at most 23.84% more taxonomically concordant, and at least 46.15% less terminally variable phylogenies compared to lower-rate sites. We find topological differences between BUSCO concatenated and coalescent trees to be marginal and conclude that higher rate sites from concatenated alignments produce the most congruent and least variable phylogenies. Finally, we show that BUSCO misannotations can lead to misrepresentations of assembly quality. To overcome this issue, we filter a Curated set of BUSCOs (CUSCOs) that provide up to 6.99% fewer false positives compared to the standard BUSCO search and introduce novel methods for comparing assemblies using BUSCO synteny. Overall, we highlight the importance of considering evolutionary histories during assembly evaluations and release the phyca software toolkit that reconstructs consistent phylogenies and reports phylogenetically informed assembly assessments.

## Introduction

High-quality reference genomes are becoming available for earth’s flora and fauna at an accelerating rate. For example, between 12 August 2022 and 21 August 2023, 7,845 new organism genomes were released by NCBI alone (Sayers et al., 2024; Sayers et al., 2023). With advancements in long-read sequencing, nuclear conformation capture and optical mapping, the reconstruction of high-quality telomere-to-telomere assemblies (Li & Durbin, 2023; Rautiainen et al., 2023) is now becoming routine across all extant clades in the tree of life (Garg et al., 2024).

Conserved single-copy orthologs are used to create phylogenies (Van Damme et al., 2022) and evaluate the completeness of new assemblies (Manni et al., 2021), yet current tools and databases remain mostly oblivious to their varying evolutionary histories and taxonomic biases. For instance, OrthoDB (Kriventseva et al., 2019) is an established database of universal orthologs, but does not specifically explore the genome-wide variations in gene presence within major taxonomic groups. Similarly, OrthoFinder (Emms & Kelly, 2019) is used to reconstruct gene trees and species phylogenies, but does not analyze phylogenetic conflicts within and between gene features in alignment sites. Moreover, the detrimental effects of disregarding information about evolutionary history when using universal orthologs for assembly completeness tests (Cunha et al., 2023) has been overlooked in popular methods (Manni et al., 2021). Hence, a systematic exploration of public genomic data has the potential to improve existing methods for the utilization of universal orthologs in phylogenomics and assembly quality assessments.

Universal single-copy orthologs are the most stable components of genomes as they remain identifiably conserved in higher eukaryotes that diverged over millions of years ago (Gundappa et al., 2022). A query set of universal single-copy orthologs (BUSCOs) (Manni et al., 2021) serves as a standard method for benchmarking gene content in newly assembled genomes. Fluctuations in BUSCO gene incidence is seen in some taxonomic groups (Cunha et al., 2023) but the full extent of BUSCO gene absence across genomically well-represented lineages has not been the subject of a focused or recent study. Although these genes remain under an evolutionary constraint of being maintained as single copies to balance dosage, polyploids (Fornasiero et al., 2024) and descendants of recently genome duplicated ancestors (Liu et al., 2020; Mansfeld et al., 2021; Wighard et al., 2022) carry fractionally elevated copy numbers. As such, BUSCO copy number variations have not been cataloged in detail across taxonomies or by gene identity.

BUSCO gene sets have been the basis for some deep molecular phylogenies (Timilsena et al., 2022; Van Damme et al., 2022). BUSCOphylo (Sahbou et al., 2022) allows users to create BUSCO phylogenies, but it is not computationally feasible for gigabase-scale genomes or a large number of taxa. It also does not explore the accuracies or inconsistencies of BUSCO-derived phylogenies. Moreover, from the perspective of molecular phylogenetics, while substitution models have been trained on empirical sequences (Jarvis et al., 2015; Misof et al., 2014; Ran et al., 2018) to improve likelihood estimates, there have been limited efforts in incorporating divergent reference genome data (Armstrong et al., 2020) to derive improved inferences. Among many unknowns, there are known sources of model inadequacies that violate basic phylogenetic assumptions. For instance, gene histories are often obscured by incomplete lineage sorting (Yan et al., 2021), horizontal gene transfer (Schrempf & Szöllősi, 2020) or hybridization (Komarova & Lavrenchenko, 2022) and sites in gene alignments may support conflicting histories due to alignment errors (Edgar, 2021), recombination, long-branch attractions (Susko & Roger, 2021) or node-density artifacts (Venditti et al., 2006). Furthermore, alignment concatenation has been shown to be statistically inconsistent for tree reconstructions (Kubatko & Degnan, 2007). This has led to many researchers assaying both concatenated and coalescent trees (Jarvis et al., 2015; Luo et al., 2022). Therefore, an explorative study that decouples sites in large, concatenated alignments from gene structures based on the column’s rate of evolution has the potential to improve current methods of phylogenomic reconstructions.

In this study, we compiled BUSCO statistics for all plant, fungal and animal genomes cataloged in NCBI Genome (Sayers et al., 2022) up to January of 2024. Our objective was to improve methods for the utilization of BUSCO genes in phylogenomics and genome completeness evaluations. Under a wide range of rate and site configurations, we assessed the capacity of BUSCO genes in reconstructing taxonomically congruent phylogenies. We tested individual trees for taxonomic concordance, and tree distributions under the same conditions for variations in terminal leaf bifurcations. Through the constructed BUSCO database, we provided evidence for 2.25% to 13.33% mean lineage-wise gene misidentifications using the most widely used default BUSCO search parameters. Categorically, we procured a Curated set of BUSCO orthologs (CUSCOs) that attains a higher specificity for 10 major BUSCO eukaryotic lineages, namely Viridiplantae, Liliopsida, Eudicots, Chlorophyta, Fungi, Ascomycota, Basidiomycota, Metazoa, Arthropoda and Vertebrata. For robust comparisons and evaluations of closely related assemblies, a syntenic BUSCO metric was derived that offers higher contrast and better resolution than standard BUSCO gene searches. Our results, data and source code have been made available through a public database and software module named phyca.

## Results

### BUSCO gene content is influenced by evolutionary history

We compiled 11,098 eukaryotic genome assemblies from NCBI and observed that genomes for new animal genera were being released at a greater rate than plants and fungi (Figure 1A). The majority of NCBI genome assemblies contained a complete or near-complete complement of single and duplicated BUSCO genes (Figure 1B). Plant lineages had a much higher mean BUSCO duplication rate at 16.57% compared to fungi and animals at 2.79% and 2.21% respectively (Figure 1B and 1C). It is known that genomes of higher ploidy are often assembled into variable sets of pseudomolecules (Fornasiero et al., 2024; Healey et al., 2024) and this is reflected in our database (Supplementary Figure 1). The mean number of observed copies for the complete BUSCO gene set had 99.05% linear correlation with the number of copies of pseudomolecules in phased and partly phased assemblies (Supplementary Figure 1). There were 169, 165 and 258 taxonomic groups out of 2,606 total that had significantly elevated means for duplicated BUSCO genes, mean BUSCO copy numbers and log assembly size respectively (Supplementary Table 1). For example, among the well-represented fungal classes, all 13 assemblies of the family Backusellaceae had duplicated BUSCOs significantly greater than other fungal groups with a minimum of 11.42% and mean of 12.18%. For the 25 assemblies in the Mucoraceae family, the minimum and mean for duplicated BUSCOs were 5.1% and 6.54% respectively. The assembly counts, mean, minimum and maximum number of BUSCO metrics for every taxonomic group including Mann-Whitney U test p-values for deviation from group means are provided in Supplementary Table 1.

**Figure 1.**
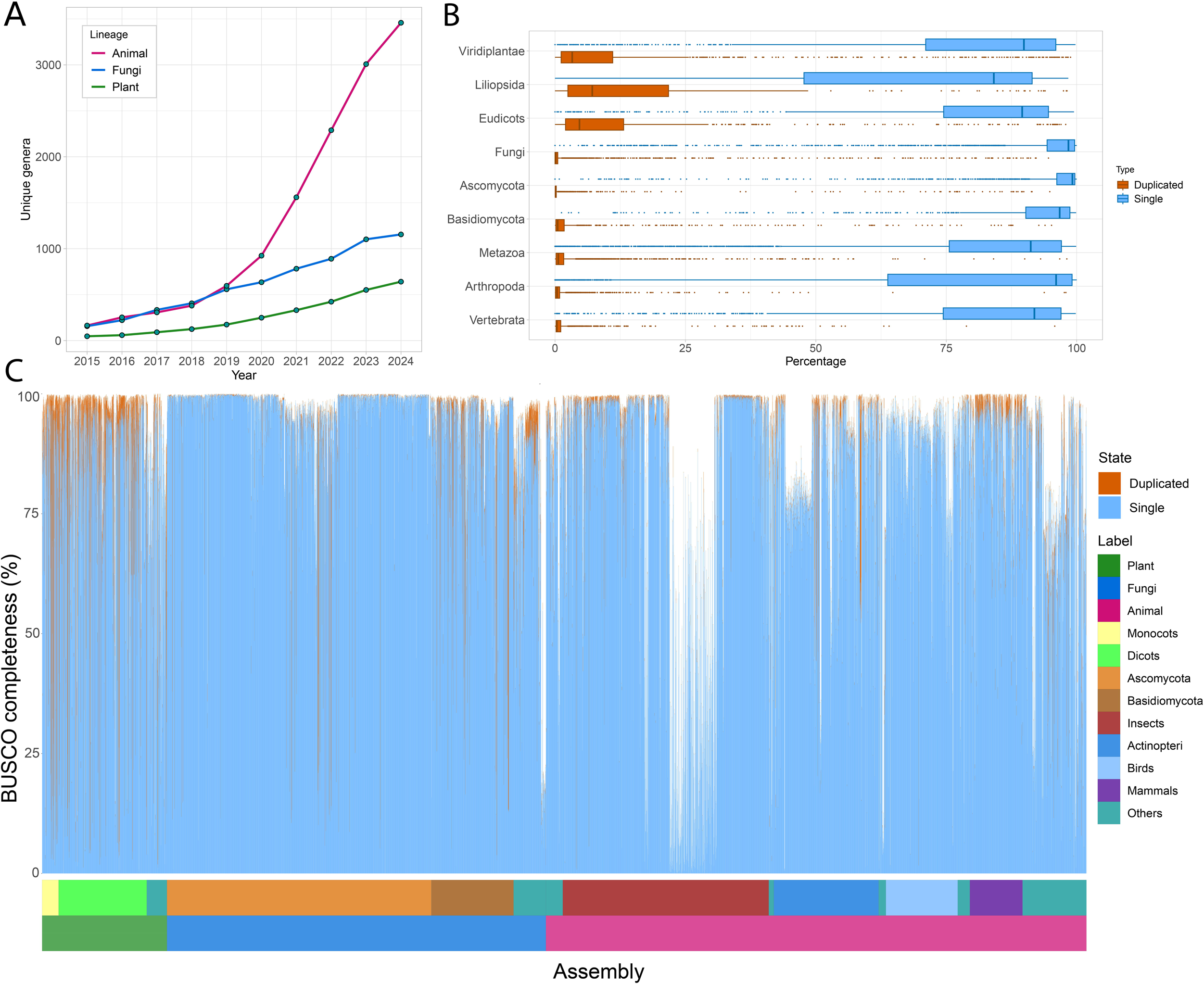
BUSCO database statistics. **A.** Genome assemblies for new genera and species are growing linearly for plants and fungi and rapidly for animals, especially in recent years. **B.** BUSCO statistics vary for plants, fungi and animals. The fraction of single-copy and duplicated genes are complementary. More duplications are observed in plants and less variation is notable for the fungi. **C.** Some taxonomic groups, such as ascomycetes and insects are better represented in NCBI genome. Assemblies from bulk genome sequencing projects with relatively low cost per genome appear as a stretch with lower BUSCO completeness. Duplicated fractions are more prominent in plants owing primarily to higher duplication rates and greater incidence of polyploidy.

Extended drops in BUSCO completeness in Figure 1C are a result of bulk genome sequencing projects that resulted in large numbers of draft genome assemblies, e.g., Ellis et al., 2021 (Ellis et al., 2021) who submitted 822 *de novo* butterfly genomes, Ronco et al., 2021 (Ronco et al., 2021) who submitted 539 cichlid fish genomes. Some taxonomic groups do show a predisposition to comparatively lower BUSCO completeness, as outlined in Supplementary Table 1. For instance, a number of *Incertae sedis* fungi-like organisms (mostly microsporidia) were found to contain <25% BUSCO genes and are seen as a dip at the trail of the fungal bars in Figure 1C (Supplementary table 1). In terms of taxonomy, it was found that across all BUSCO lineages and taxonomic levels, 215 groups had significantly different mean BUSCO completeness. The complete database, along with taxonomic classifications, assembly and BUSCO statistics are available to download and view at www.phyca.org.

### Sites evolving at higher rates produce more taxonomically congruent phylogenies

From our compiled data, we sought to determine the best way to utilize BUSCO genes to create broad whole-genome phylogenies spanning large evolutionary distances. Individual phylogenies were tested for agreement with NCBI taxonomic classifications. To assess taxonomic congruence, we created 3,566 phylogenetic trees for the 5 largest BUSCO lineages in terms of assembly and gene count. Our tests were focused on the Eudicots, Ascomycota, Basidiomycota, Arthropoda and Vertebrata lineages. Gene alignments for divergent taxa varied significantly based on parameters passed to the alignment algorithm (Supplementary figure 2). Different lineages had different rate profiles for aligned sites (Supplementary figure 3). Algae, fungi and early diverging metazoans displayed greater site heterogeneity in their alignments (Supplementary figures 2 and 3).

Phylogenetic trees under different evolutionary rates and alignment lengths were compared for taxonomic congruity. Variations of the LG and JTT substitution models (Le & Gascuel, 2008) with different rate categories were consistently found to have the highest likelihood under all conditions (Supplementary Table 2). The top 5 best substitution models based on Bayesian Information Criterion (BIC) for each condition with model comparison metrics are included in Supplementary Table 3. The number of unique amino acid residues in an alignment column was used as a proxy for site evolutionary rate. Sites evolving at higher rates together with longer alignments generally produced more taxonomically concordant trees (Figure 2A and Supplementary figure 4). Taxonomic concordance was predominant in eudicots with either 68 or 69 out of 69 total families (98.55-100%) being reconstructed as monophyletic above 4,000 alignment length and 5 or more unique amino acids. In arthropods and vertebrates, up to 113 out of 125 (90.40%) and 187 out of 225 (83.11%) respectively were reconstructed as monophyletic. In ascomycetes and basidiomycetes, only up to 60 out of 97 (61.86%) and 63 out of 88 (71.59%) respectively were found monophyletic in any single condition. The lineage and condition-wise monophyly counts are presented in Supplementary Figure 4. For each lineage, a consistent number of families were resolved as monophyletic in most of the trees, whilst some families precariously only appeared monophyletic at certain conditions (Figure 2B and Supplementary table 4). Alignments with greater numbers of sites and unique residues almost always resolved greater numbers of families (Figure 2C). Rate effects were more potent than alignment length (Figure 2D). For instance, 32 families out of 543 total were monophyletic under all tested conditions. Of the remaining 511, 59.47%, 84.61% and 86.53% were monophyletic when reconstructed with 2, 8 and 14 (low, moderate and high) unique amino acids per column respectively and 67.18%, 80.12% and 83.32% were reconstructed as monophyletic with 1,000, 5,000 and 10,000 alignment lengths respectively. Under conditions where the alignments did not provide sufficient information to accurately resolve tree topology (Supplementary figure 5), likelihoods were higher at greater rates and alignment lengths (Figure 2E). Variations in taxonomic concordance receded with increasing site counts and evolutionary rates in eudicots, arthropods and vertebrates, but the pattern was less prominent in ascomycetes and basidiomycetes (Figure 2F-G and Supplementary Figure 6).

**Figure 2.**
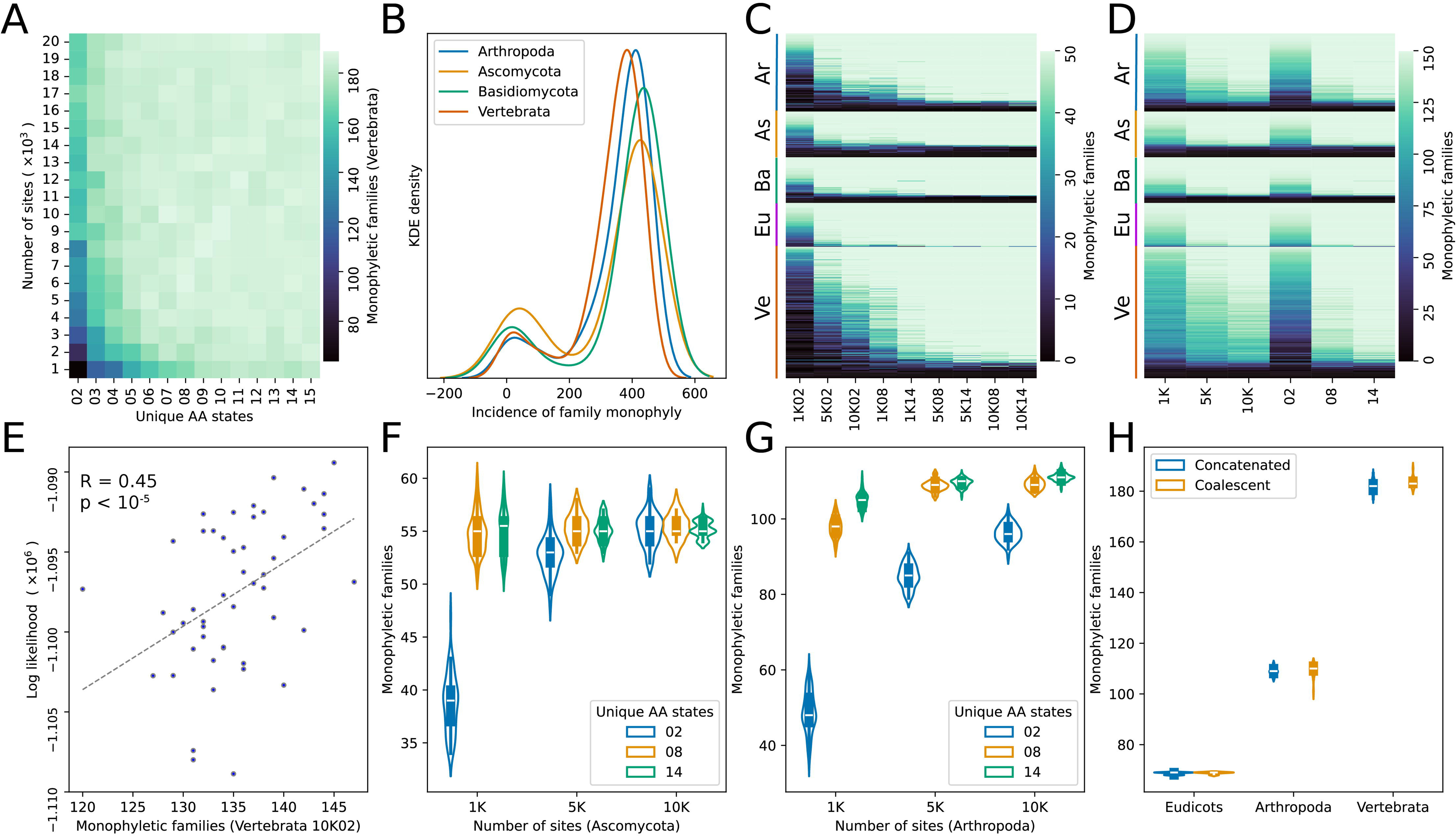
Higher rates are more informative and produce better phylogenies overall. **A.** Taxonomic concordance across 13 rate profiles and 20 alignment lengths. Sites evolving at higher rates and longer alignments share more agreement with taxonomic groupings. **B.** Most families in 4 groups are resolved as monophyletic in most trees whereas a smaller number of families are more sporadic and appear to be monophyletic either randomly or under specific rates. **C.** Ve, Eu, Ba, As and Ar represent in Vertebrate, Eudicots, Basidiomycota, Ascomycota and Arthropoda lineages respectively. Each vertical bar is a unique family. With few exceptions, families are more likely to be found monophyletic at greater rates and sites. **D.** Increasing rates have a greater effect on tree concordance relative to increasing sites. **E.** Under optimum tree search conditions, tree likelihoods correlate with taxonomic agreement. **F.** Ascomycota represents the fungal clade which shows increased variance in tree sets and is less responsive to rate and site adjustments. **G.** For the Arthropoda lineage, increasing rates and sites increases concordance and reduces tree set variance. **H.** Differences between concatenated and coalescent species trees are marginal.

We observed that all five tested lineages showed a similar trend where 462 out of 543 families were found monophyletic at the most informative condition with 14-character columns and an alignment length of 10,000 (Figure 2C and Supplementary table 4). Of the remaining 81, 42 families could not be resolved as monophyletic (0 out of 50 trees) and the monophyly status of the remaining 39 families remained inconsistent. Rate preferences for monophyly in the queried families were not observed. The Petroicidae family of birds was the only family that yielded monophyletic trees across all 50 trees at rate condition 8, but was not consistently monophyletic in the higher rate condition of 14 with monophyly in 49 out of 50 trees in alignments of length 10,000 (Supplementary table 4).

To interpret the relationship between tree likelihoods and taxonomic concordance, we recomputed likelihoods for all trees under a fixed set of alignments. Correlations between mean tree likelihood and taxonomic concordance diminished with longer alignments and faster evolving sites (Supplementary figure 5). At the same time, tree topologies were more stable at the terminal taxa for all lineages at higher evolutionary rates and greater site counts (Supplementary figure 7). The sets of trees created from sites with 8 unique characters and an alignment length of 10,000 for eudicots, arthropods and vertebrates were compared to BUSCO coalescent trees to contrast tree concordance between the two popular methods. No significant variations in taxonomic agreement between concatenated trees and trees created under the multispecies coalescent model were observed (Figure 2H).

### A filtered BUSCO set provides improved assembly assessments

Across all 10 lineages, on average 2.25% to 13.33% of BUSCO genes were misidentified in genomes where all BUSCO genes had been removed (Figure 3A and Table 1). Misidentification implies that a default BUSCO search would not identify divergent copies of these genes and the absence of the identified BUSCO gene in a query assembly would result in the inadvertent identification of the divergent copy. The magnitude of misidentification rates varies by lineage and was observed to be lowest across the fungal assemblies and highest across vertebrate and plant assemblies (Figure 3A and Table 1). Roughly 10% of BUSCO genes in all 10 lineages were misidentified at a far greater number of assemblies than others (Supplementary figure 8). Assessment of BUSCO completeness with these genes removed resulted in reduced numbers (Table 1) of BUSCO gene misidentifications in all lineages (Figure 3B). The reduction in false hits was more pronounced in the Vertebrata, Liliopsida, Eudicots and Chlorophyta lineages (Figure 3B and Table 1). For clarity, the Curated set of BUSCO genes has been named CUSCOs and the remaining Misannotation-prone BUSCO genes are hereon abbreviated as MUSCOs.

**Figure 3.**
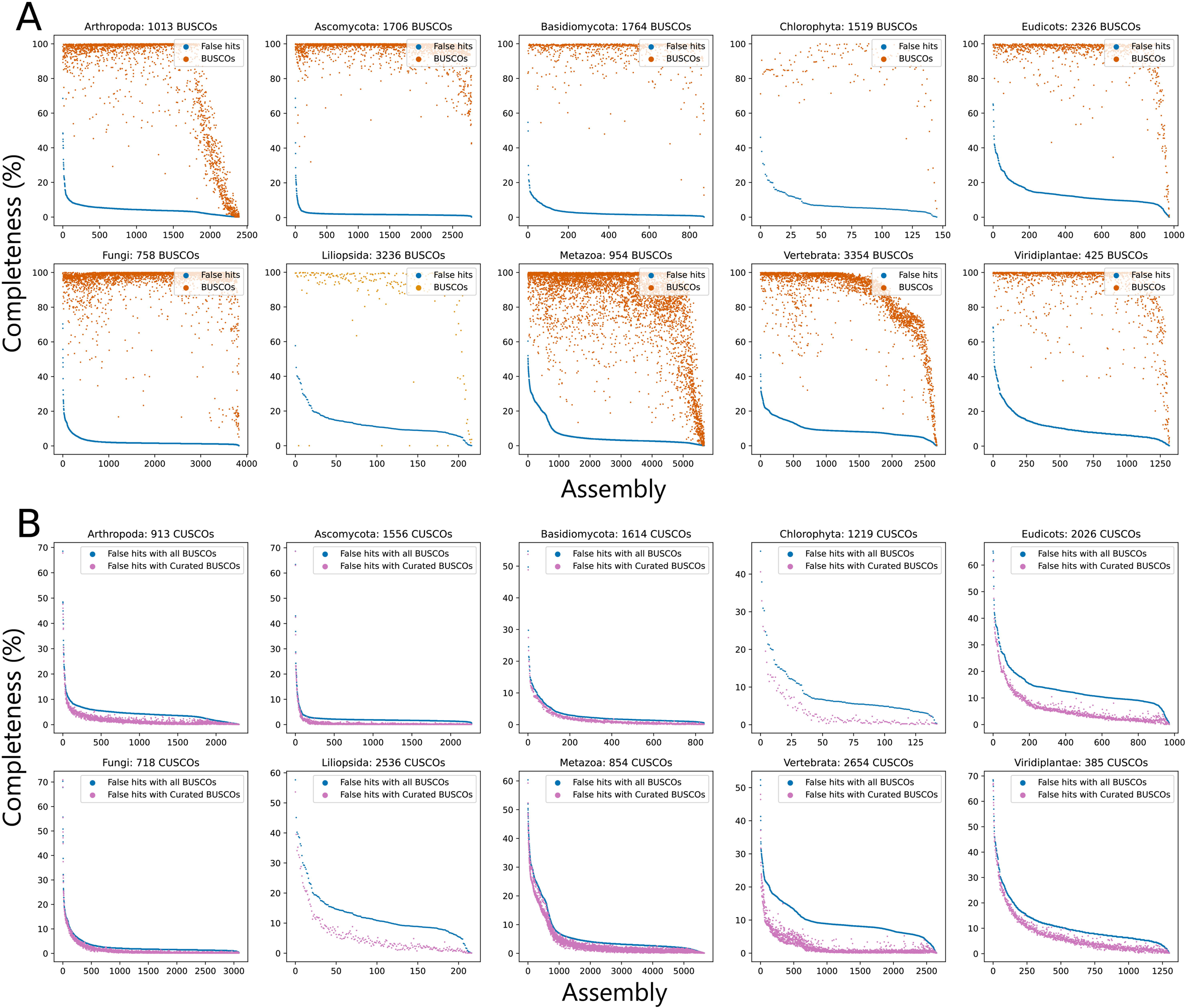
Removal of erratic BUSCO genes reduces BUSCO misidentification rates. **A.** BUSCO genes are misidentified at different rates in different lineages. Median fraction of false identification is around 15% for most plants and vertebrates, but much lower in fungi. **B.** Only considering our Curated set of BUSCO genes (CUSCOs) markedly reduces false hits in some lineages.

**Table 1.** BUSCO and CUSCO misidentification rates.

| Lineage | BUSCO<br>completeness<br>(mean) | BUSCO<br>completeness<br>(SD) | CUSCO<br>completeness<br>(mean) | CUSCO<br>completeness<br>(SD) | BUSCO<br>false hits<br>(mean) | BUSCO<br>false hits<br>(SD) | CUSCO<br>false hits<br>(mean) | CUSCO<br>false hits<br>(SD) |
| --- | --- | --- | --- | --- | --- | --- | --- | --- |
| Viridiplantae | 91.88 | 15.90 | 90.67 | 18.37 | 11.35 | 8.69 | 7.52 | 8.88 |
| Liliopsida | 87.30 | 24.47 | 90.06 | 19.54 | 12.60 | 8.01 | 5.90 | 7.70 |
| Eudicots | 92.37 | 15.00 | 91.81 | 16.52 | 13.34 | 7.30 | 6.35 | 7.40 |
| Fungi | 94.66 | 12.99 | 94.93 | 12.91 | 2.86 | 4.01 | 1.64 | 4.02 |
| Ascomycota | 96.74 | 6.92 | 96.84 | 7.13 | 2.25 | 2.97 | 0.69 | 3.01 |
| Basidiomycota | 95.59 | 8.08 | 95.29 | 9.27 | 3.00 | 3.98 | 2.02 | 3.86 |
| Metazoa | 83.41 | 23.86 | 82.32 | 25.47 | 6.02 | 7.44 | 4.00 | 6.69 |
| Arthropoda | 81.86 | 29.35 | 78.86 | 32.58 | 4.55 | 3.92 | 1.92 | 3.75 |
| Vertebrata | 85.09 | 19.63 | 84.72 | 20.07 | 9.57 | 5.12 | 2.17 | 3.74 |
| Chlorophyta | 86.09 | 14.29 | 85.19 | 16.37 | 8.12 | 6.80 | 3.57 | 6.09 |

\*all numbers are in percentiles

We analyzed the incidence of BUSCO misannotations by assembly and gene identity to extrapolate the source of this phenomenon. Gene misannotations were found to be more weighted towards the query gene rather than the query genome assembly (Figure 4A). Removal of MUSCOs resulted in better assembly assessment metrics and shifted the assembly quantiles of BUSCO misidentification towards the gene quantities (Figure 4B). Correlation analysis of lineage-wise misannotation rates with assembly metrics revealed that BUSCO gene misidentifications correlated most with the mean number of BUSCO copies in the assembly, a metric we termed inflation (Figure 4C). Other variables showing the highest correlations were the number of miniProt hits (MPH) and the log of assembly size, being more pronounced in chlorophytes and vertebrates, respectively (Figure 4C).

**Figure 4.**
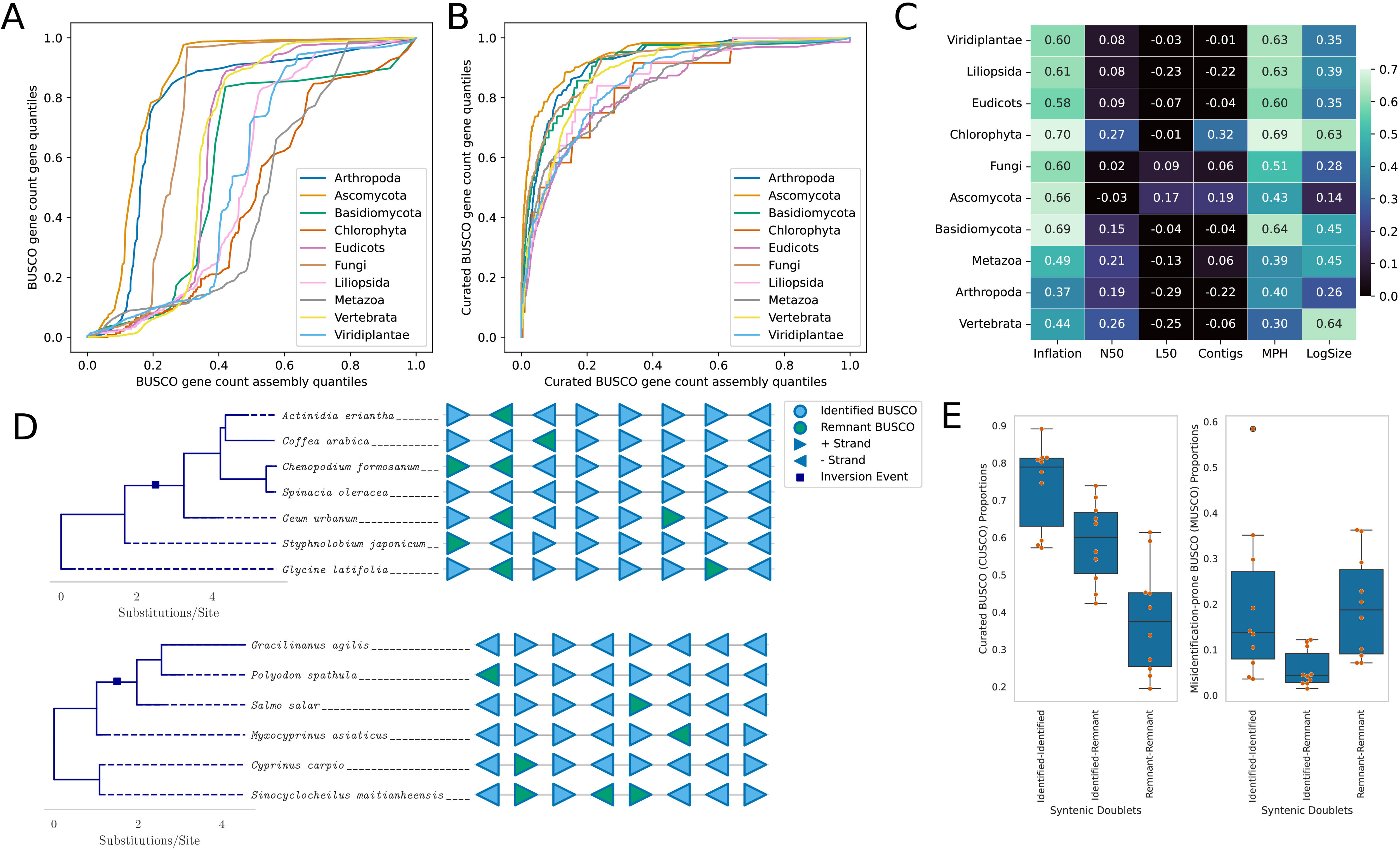
Misidentification events are weighted more towards the identity of the gene rather than assembly and false hits correlate most with assembly complexity and gene content. **A.** A graph of gene quantiles against assembly quantiles for false hit counts shows that the majority of assemblies show some false gene hits but the gene quantiles rise more shapely. **B.** Considering only the curated BUSCO set shifts the assembly quantiles at the lower range towards the genes. CUSCO genes are misidentified in far fewer assemblies and do not show assembly preference. **C.** False identification rates correlate most with the number of miniProt hits (MPH) and mean BUSCO copy counts (Inflation). Moderate correlation to the log of assembly size is also observed. **D.** Two example blocks of 8 genes conserved beyond the species level for eudicots (top) and vertebrates (bottom) showing misidentified/remnant BUSCO genes in syntenic order. **E.** CUSCO and MUSCO proportions for syntenic doublets with 0, 1 and 2 remnant genes. Remnant proportions gradually recede for CUSCOs, but rise back up in remnant doublets for MUSCOs.

Given the observed preponderance of misannotation rates in complex genomes in terms of assembly size, gene hits and BUSCO inflation (Figure 4C), we analyzed the syntenic patterns of identified and misidentified BUSCO genes to query potential evolutionary origins. For computational feasibility, all possible permutations of identified and misidentified BUSCO genes in 10 sets of gene blocks harboring up to 10 genes were tested. Gene block analysis revealed that beyond the species level, misidentified BUSCO genes are preserved in syntenic order at the highest rates in the Liliopsida, Viridiplantae and Eudicots lineages at 4.07%, 3.97% and 3.78% respectively. The fourth highest rate of syntenic misidentifications was in the Basidiomycota at just 0.88% and the lowest was in Arthropoda at 0.14%. Two such representative gene blocks from the Eudicots and Vertebrata lineages are shown in Figure 4D top and bottom respectively. This suggests that some misidentified BUSCO genes are remnants of gene duplication events where the syntenic copy became more divergent. Details for all computed gene blocks are available to download at www.phyca.org/data.html. The syntenic analysis was extended to our complete data set with syntenic gene pairs to determine whether CUSCO and MUSCO genes contained pairs with one and two remnant genes in similar proportions. CUSCO syntenic doublets were progressively found in lower proportions with one and two remnant genes (Figure 4E). However, MUSCO syntenic doublets appeared in similar proportions with pairs of identified and pairs of remnant genes (Figure 4E). MUSCO genes are therefore more syntenic in the remnant-remnant configuration compared to CUSCO genes.

### BUSCO collinearity is an indicator of pseudomolecule quality

To demonstrate the utility of BUSCO synteny in assembly comparisons, we compiled and compared 1035 pairs of genomes of the same species with contrasting quality metrics. We employed an adjusted Intersection Over Union (IoU) metric with BUSCO gene doublets found in the same order and orientation to compare two assemblies. The denominator is adjusted by the difference in the number of contigs such that highly fragmented assemblies with the same gene order and orientation would be syntenically equivalent to highly contiguous assemblies. Hence, the syntenic doublet metric is designed to only capture differences in gene synteny and to not be influenced by varying numbers of contigs in query assemblies (Supplementary Figure 9). BUSCO syntenic connections were able to capture far greater contrast in the assembly pairs compared to simply the difference in BUSCO completeness (Figure 5A). Syntenic BUSCO connections decayed exponentially with phylogenetic distance in our six non-overlapping BUSCO lineages (Figure 5B and 5C). We further compiled the 40 least contiguous NCBI assemblies of *Oryza sativa*, *Mus musculus*, *Drosophila melanogaster*, *Ovis aries* and *Arabidopsis thaliana* to represent the BUSCO syntenic distance between the assemblies as a dendrogram. Metrics for the full set of assemblies are provided in Supplementary Figures 10, 11, 12, 13 and 14 respectively. An example of a dendogram with 8 fragmented *Mus musculus* assemblies and a highly contiguous reference assembly is shown in Figure 5D. Less contiguous assemblies were found to be at greater syntenic distances to the higher quality assembly, implying greater numbers of BUSCO misidentification events or more extensive misassemblies.

**Figure 5.**
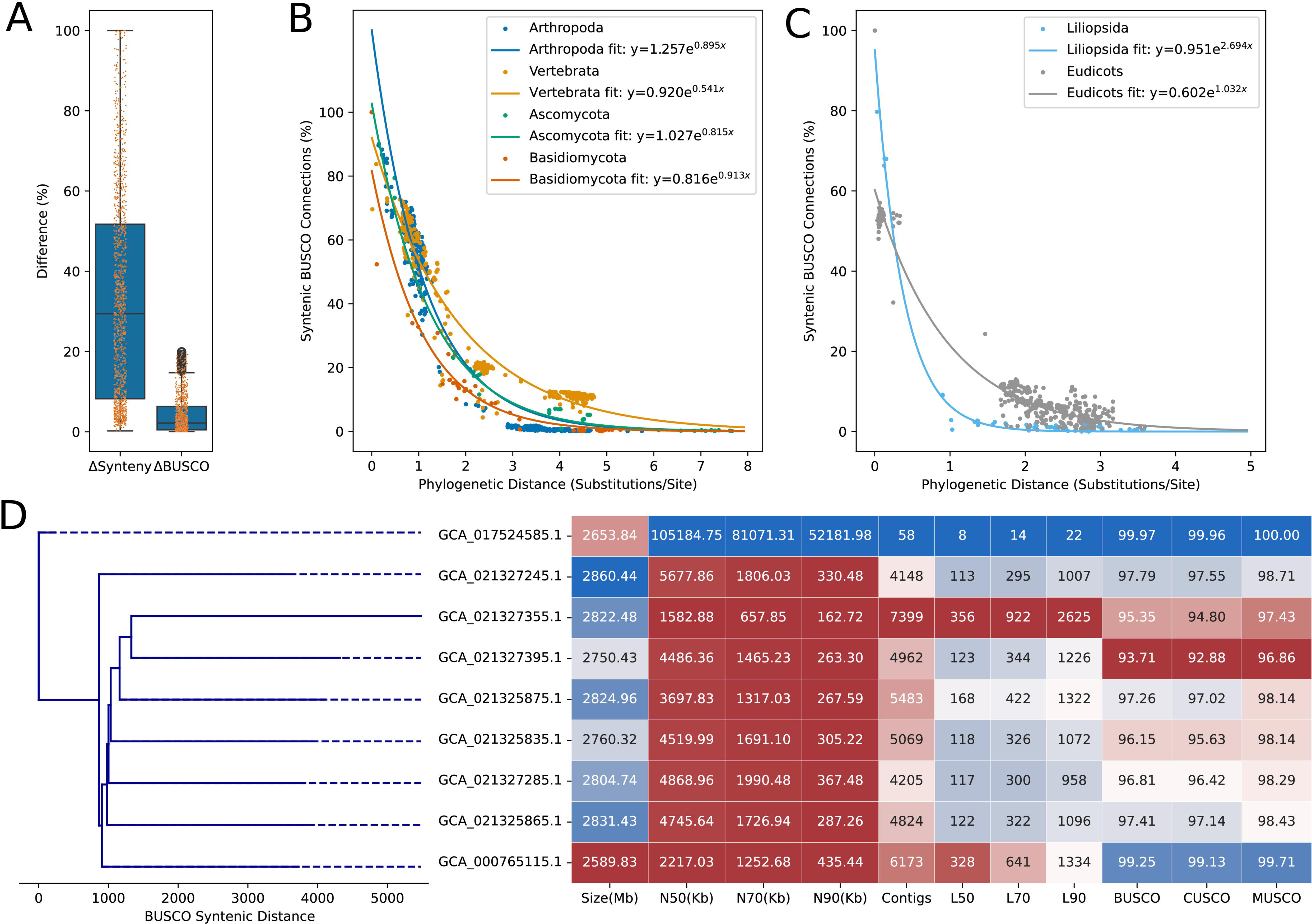
BUSCO syntenic distance offers greater contrast than BUSCO content, decays exponentially with phylogenetic distance and serves as a robust metric to compare closely related assemblies. **A.** Boxplot showing differences in BUSCO completeness and BUSCO syntenic distance between 1035 sets of assemblies that vary in quality. **B.** General trend and histogram of BUSCO syntenic distance and BUSCO completeness differences. BUSCO syntenic differences can offer far greater contrast. **C.** Exponential decay and curve function of BUSCO syntenic similarity for Arthropoda, Vertebrata, Ascomycota, and Basidiomycota lineages **D.** Exponential decay and curve function for Liliopsida and Eudicots lineages **E.** Eight highly fragmented *Mus musculus* assemblies compared against a highly contiguous assembly through BUSCO syntenic distance and quality assembly metrics.

To further assess how BUSCO synteny can indicate assembly quality, we visualized chromosome-wise BUSCO collinearity in a set of *Oryza* assemblies as a case study. The *Oryza* genus is genomically well characterized with several state-of the-art chromosome level assemblies (Fornasiero et al., 2024). We demonstrate with a draft assembly (GenBank ID: GCA_009805545.1) and a high-quality assembly of *Oryza longistaminata* (Reuscher et al., 2018) that BUSCO synteny can provide greater contrast between assemblies of varying quality compared to BUSCO metrics alone (Figure 6). Between the two *O. longistaminata* assemblies, although the number of curated BUSCO genes identified was comparable (98.82% and 93.17%), BUSCO collinearity was not preserved across the closely related sister taxa within the genus (Figure 6). These observed syntenic deviations are quantified by our adjusted IoU metric based on BUSCO gene connections (Supplementary Figure 9) and the syntenic distance between the two *O. longistaminata* assemblies was 82.25%. The full set of chromosomes for this test case is available on the phyca website at www.phyca.org/data.html. The phyca software package allows users to similarly compare and visualize syntenic distances between assemblies and query genomes.

**Figure 6.**
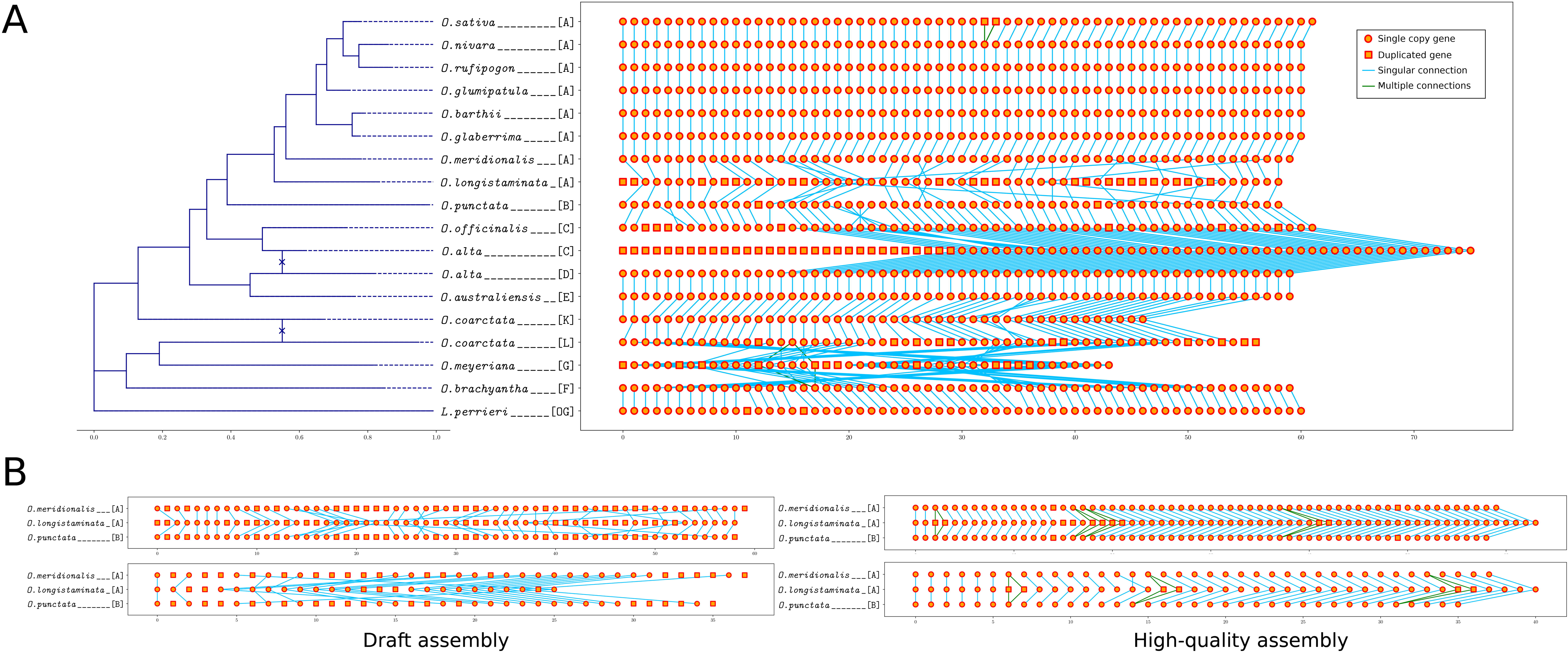
Phylogenetic and syntenic information improves assembly assessment. **A.** BUSCOs in chromosome 1 of *Oryza longistaminata* and *O. meyeriana* assemblies are less syntenic to sister taxa. A chromosomal translocation event from chromosome 3 to 1 in *O. alta* subgenome C is also visualized. **B.** Assessment of an improved *O. longistaminata* assembly reveals that BUSCO genes were either misidentified or contigs were scaffolded poorly in the inferior assembly. Chromosomes 1 and 7 are visualized at the top and bottom respectively.

## Discussion

Here, we presented our studies across three facets. First, we determined the prevalence of BUSCO gene variations by taxonomy through the compilation of available plant, fungi and animal genomes in the public domain. Second, we optimized site conditions for consistent phylogenomic reconstructions by maximizing taxonomic congruity and minimizing tree set variability. We then created large whole-genome phylogenies under the best determined conditions for 10 major BUSCO lineages. Third, we provided evidence for BUSCO misannotations with the current software defaults and filtered a curated set of BUSCO genes for better genome quality assessments. To mitigate the effects of BUSCO misannotations during assembly evaluations, we described a novel method of comparing assemblies with BUSCO synteny that provides much better contrast for closely related assemblies of varying quality.

### BUSCO completeness and copy number variations

Universal genes have been instrumental for querying gene space completeness and assembly quality (Manni et al., 2021). Our results show that the evolutionary history of a genome influences its BUSCO score and that this influence is prevalent in many taxonomic groups rather than just a few (Cunha et al., 2023). It was also observed that some groups vary more dramatically than others in BUSCO metrics (Supplementary table 1). Therefore, for assemblies from early diverging groups with few extant taxa or available genomes, BUSCO genes may provide an inadequate representation of gene space completeness. Given these observations, we propose that it is necessary to consider the evolutionary history of related taxa when evaluating the gene content of new genome assemblies.

Assembly gene content is influenced drastically by evolutionary history. Polyploid organisms are known for being able to maintain multiple sets of single-copy orthologs (Fornasiero et al., 2024) and genomes fractionate at varying rates post-duplication (Garsmeur et al., 2014). It is likely that groups that were found to harbor large sets of duplicated BUSCO genes in haploid assemblies have either experienced recent whole-genome duplication events or have adjusted their gene regulation to accommodate an inflated complement of some single-copy orthologs. The set of genes that are more likely to be misidentified (Supplementary figure 8) are likely tolerated more in genomes at greater copy numbers. This is supported by the high correlation of gene misannotations to the BUSCO inflation metric shown in Figure 4C and the preservation of some syntenic remnant genes across large phylogenetic distances (Figures 4D and 4E). It is probable that misannotation-prone genes duplicated and subsequently functionalized in ancient ancestral genomes multiple times. Some of the duplicated copies may have taken up important functions that prevented the sequences from diverging drastically and the shared homology is now responsible for the observed false hits. The availability of a consolidated database of BUSCO results from public genomes allows researchers to derive meaningful copy number expectations for BUSCO genes in new assemblies based on evolutionary history.

### Decoupling aligned sites from gene features and a case for fast evolving columns

Likelihood estimation in phylogenetics assumes that all sites evolve independently (Yang, 2006). Since this is not biologically meaningful (Nasrallah et al., 2010), advanced tree search algorithms split columns into invariant sites (Yang, 1996) and several rate categories (Yang, 1994) to address rate heterogeneity. We assumed that unique amino acid counts in aligned columns could serve as a proxy for evolutionary rate at that site and filtering sites by evolutionary rate would decouple sites from intragenic evolutionary influences. In practice, researchers often select fast evolving sites for dense phylogenies (Matschiner et al., 2020) and conversely, for deep phylogenies, they tend to use slowly evolving sites to optimize information content in the alignment (Misof et al., 2014). Our study broadly highlights the practical effects of rate variation and alignment information content on tree reconstruction. Rosenberg and Kumar, 2001 (Rosenberg & Kumar, 2001) showed that the number of sites have greater effect on tree accuracy compared to substitution rates. On the contrary, our results show that when an adequate number of sites are sampled (Figure 2D), site evolutionary rate has a greater effect on tree accuracy in terms of taxonomic congruity. In our studies, higher rate sites were generally found to produce better trees and there was minimal hindrance caused by long-branch attraction biases and heterotachy (Figure 2C and 2D).

Slow evolving sites have been favored throughout the history of molecular phylogenetics (Pisani, 2004). Slow-fast analysis was popularized for phylogenetic reconstructions in the context of substitution saturation and long-branch biases (Cummins & McInerney, 2011; Kostka et al., 2008). Similarly, chi-squared tests are employed to detect compositional heterogeneity in alignments (Boudinot et al., 2023; Foster, 2004). The primary goal of these analyses has been to identify and prune fast evolving sites to improve phylogenies (Pisani, 2004). Such practices have recently been perceived with scrutiny (Superson & Battistuzzi, 2022) and Rangel and Fournier, 2023 (Rangel & Fournier, 2023) has shown that fast evolving alignment sites can be highly informative. We show in Figure 2 (and Supplementary table 4) that higher rate sites improve taxonomic concordance across almost all 543 families tested, and always increase tree set consistencies (Supplementary figure 7) compared to lower rate sites. Therefore, contrary to popular practices, our results suggest that with adequate taxon sampling, faster rates for protein characters may produce more accurate phylogenies regardless of node depth.

### Phylogenies within the kingdom Fungi and recalcitrant evolutionary histories

Some taxonomic classifications in the fungal domain are based on molecular ITS data (Carbone et al., 2017). Although ITS-based primers are commonly used for phylogenetic placement, the drawbacks of ITS sequences are apparent. RNA code has fewer letters than protein code and the ITS sequences are much shorter than most protein coding genes. Further, rRNA genes appear in large copy numbers (Lavrinienko et al., 2021; Lofgren et al., 2019) making them amenable to multiple evolutionary histories at greater divergence times. In contrast, single-copy orthologs exist under dosage restraints and this generally prevents copy number variations from persisting throughout evolutionary timescales (Garsmeur et al., 2014). Additionally, sampling greater numbers of taxa generally has a strong positive effect on phylogenetic accuracy (Heath et al., 2008) and BUSCO genes offer the means to include highly divergent clades. For these reasons, it is reasonable that BUSCO genes would be able to resolve deeper phylogenies with greater precision than ITS sequences.

We found taxonomic classifications to be more obscure for the kingdom fungi. Although tree entropy at the termini reduced by about 50% (Supplementary figure 7), we did not observe the same level of gradual reductions in the variance of monophyletic counts as seen from plants and higher animals (Supplementary figure 6). One likely explanation for these complications is their significantly higher rate of evolution and shorter generation times compared to other clades (Naranjo-Ortiz & Gabaldón, 2020). This can be seen in the greater fraction of high-rate sites shown in the state frequency spectra in Supplementary figure 3. This effect in conjunction with their compact genome sizes, relatively higher rates of gene flow (Gonçalves & Gonçalves, 2022) and very short generation times compared to higher eukaryotes makes the accurate reconstruction of fungal evolutionary histories challenging. Despite these challenges, the fungal families did follow the same trend as the higher eukaryotes in response to increasing evolutionary rates in Figures 2C and 2D, albeit a greater fraction of families seemed to have members descended from more than one most recent common ancestor. The greater fraction of non-monophyletic groups could be an artifact of the limitations of the standard ITS-based classification scheme. These views are supported by a 9.72% observed higher fraction of monophyly in the higher fungi, basidiomycetes compared to the lower fungal phyla, ascomycetes (Supplementary table 4). Furthermore, compared to the other three lineages tested, ascomycetes and basidiomycetes show noticeably greater numbers of monophyletic groups with alignments of slowly evolving sites (Supplementary tables 4 and 5). One cause behind this could be higher rates of alignment errors in more distantly related taxa. In this regard, we did not consider the consistently reproduced alignments in Supplementary Figure 2 to be infallible since they are biased by the heuristics of multiple-sequence alignment algorithms (Edgar, 2021). Additionally, the higher range of monophyly in shorter alignments (Supplementary figure 4) could be explained by ITS-derived taxonomic classifications since those alignments resemble ITS alignments better in terms of length, slower rates of evolution and overall information content. Because of these ambiguities, and a higher fraction of families found not to be monophyletic, the ascomycetes and basidiomycetes weren’t included in the coalescent study in Figure 2H.

### BUSCO provides a standard for whole-genome phylogenies

At present, both concatenated and coalescent phylogenies are used in practice (Jarvis et al., 2015; Luo et al., 2022). The multispecies coalescent model corrects for incomplete lineage sorting to resolve ancestral relationships in higher taxa speciating from large populations. Jian et al., 2019 (Jian et al., 2019) showed that the multispecies coalescent outperforms concatenation across a range of metazoan groups. Our results suggest that such differences are usually marginal in terms of taxonomic congruity (Figure 2H). In the arthropods for instance, the coalescent tree set demonstrates a greater range of variation. It is also important to note that the total number of sites in the coalescent trees were far greater than the concatenated trees since up to 75 whole genes were included. For comparison, the vertebrate tree likelihoods were still improving at the 10,000 site count mark (Supplementary figure 5). We propose that when there is adequate information content in the alignments, the high dimensional likelihood surface flattens out harboring several vicinal and localized peaks and valleys. This results in the distribution of alternate topologies with varying model likelihoods spread out within a range of monophyly counts in the correlation plots shown in Supplementary figure 5. We thus conclude that the multispecies coalescent offers a powerful framework, but results should still be interpreted with caution, and our BUSCO concatenation method offers a robust alternative when suitable.

The search space for phylogenetic trees grows faster than exponentials with increasing numbers of terminal nodes (Yang, 2006). Our smallest tested tree had 592 terminal nodes which equates to a search space of 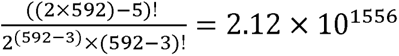. This high number of taxa makes the tree space numerically intractable even with the best available heuristics. The exact same tree topology was never reproduced in our results under any condition. From our evaluations of the tree distributions, we suggest that: 1) consistent reconstruction of a greater number of groups as monophyletic offers support for internal nodes, and 2) reduced terminal variability in tree distributions provides confidence for accuracy of overall tree topology. Combined, ancestral histories reconstructed from our method of sampling high-rate sites from whole-genome BUSCO data should be deemed more reliable than ITS or gene trees, and on par with coalescent-based trees. In the phyca website and software package, the 39 clades with undetermined monophyly status have been shared in Supplementary Table 4 to alert users to be cautious about drawing interpretations. It is important to be aware that with large datasets, model inadequacies (Delsuc et al., 2005) could result in erroneous topologies having high support values. It is therefore possible that for any individual taxa or clade, the reduced terminal variability in our tree sets may have reinforced erroneous placements. We recommend that researchers with more nuanced evolutionary questions should consider rebuilding subtrees within their clade of interest. For this purpose, phyca provides a user-friendly implementation of our proposed methods to construct phylogenies from user defined sets of query taxa.

### Shortcomings of homology-based and probabilistic gene predictions

BUSCO has been the unrivaled standard for gene space completeness tests since 2019 (Seppey et al., 2019). BUSCO relies on sequence homology searches through sequence alignments and subsequent refinement of search results by trained hidden Markov models (Manni et al., 2021). In general, alignment-based methods for gene identification are employed using arbitrary cutoffs (Levy Karin et al., 2020) and probabilistic models are used with empirically trained probabilities (Edgar, 2021; Wheeler & Eddy, 2013). BUSCO gene prediction by Compleasm (Huang & Li, 2023), a better implementation of BUSCO, starts with a miniProt (Li, 2023) search that is restricted to report duplicate genes only if the alignment score is at least 95% of the best alignment. Compleasm has four additional threshold parameters for secondary hits, gene identity, fraction and completeness respectively. These thresholds have been empirically optimized by the developers to maximize precision and recall (Huang & Li, 2023). Almost all user-reported BUSCO results are reported based on default parameters (Ellis et al., 2021; Fornasiero et al., 2024; Healey et al., 2024; Liu et al., 2020; Mansfeld et al., 2021). Readjustment of these parameters would adversely alter the preoptimized tunings, and for experimental explorations, there would be an inordinate number of permutations to consider. Our method of removing genes and rerunning under default settings mimics the effect of natural gene loss events. Our analysis of false positive hits revealed a set of less reliable BUSCO genes with a significantly higher propensity of being misannotated (Supplementary figure 8). We surmise that for gene predictions there may be no “one glove fits all” method that will work for all genes across all possible lineages. With this view in mind, integrative approaches have been suggested in the past to improve gene prediction accuracies (Alam & Chowdhury, 2020). We conclude that putative gene prediction is a tricky endeavor and demonstrate in Figure 3B and Table 1 that removing the less reliable genes from the BUSCO gene set improves precision without compromising recall.

## Conclusion

Universal orthologs are critical inferential tools for evolutionary genomic research. To improve the utilization of BUSCO genes in this field, we first compiled and comprehensively analyzed their presence and copy number variations within the expansive higher eukaryotic domain. Based on our findings, we suggest that evolutionary histories must be considered for proper interpretation of BUSCO completeness metrics. Second, we determined the extent to which the ancestral histories of major eukaryotic lineages could be resolved through universal single-copy orthologs. Our results imply that columns evolving at higher rates in alignments of protein characters are more robust for deep phylogenomic reconstructions. We described a novel way to consider phylogenetic accuracy using taxonomy and a simplified way to express tree set variability by enumerating terminal leaf bifurcations. In light of our findings, we produced the largest unified nuclear genome-based phylogenies for 10 major taxonomic groups in the plant, fungi and animal kingdoms to date. Within these phylogenies, we highlighted clades that were consistently reconstructed as monophyletic with respect to their taxonomic labels and distinguished clades that demonstrated more recalcitrant ancestral histories. Finally, our database yielded a filtered set of BUSCO orthologs that provide a better representation of assembly gene content compared to the standard BUSCO search. We showed that more robust evaluation of genome quality can be attained through the incorporation of BUSCO syntenic information from related assemblies. Our processed data and tools have been made easily accessible for robust phylogenomic reconstructions, rapid placement of query assemblies by appending BUSCOs to large, precomputed alignments and for deriving phylogenetically informed assembly quality evaluations.

## Materials and methods

### Database compilation and classification

Metadata for plant, fungi and animal genome assemblies were sourced from the NCBI genome database (Sayers et al., 2022) accessed on January 14, 2024. Assemblies flagged by NCBI as partial and contaminated were not used. Special characters (\’()-/#:=+[]) were removed from organism names to avoid software errors during automation. The assembly metadata were sorted by level of assembly set by NCBI (complete, chromosome, scaffold, contig), date of release (newest to oldest) and assembly size (largest to smallest) respectively. Only the top entry for identical organism names was kept. Batch downloads were executed using the cURL application (www.curl.se). The NCBItax2lin software (https://github.com/zyxue/ncbitax2lin) was used to assign taxonomic classifications at the phylum, class, order, family and genus levels to the assemblies. The Mann-Whitney test was used to test the hypotheses of whether assemblies within a taxonomic group had a significantly different mean for a metric compared to all assemblies in the BUSCO lineage. A Bonferroni correction of 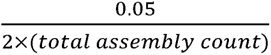 was carried out to determine the p-value cutoff thresholds.

### Finding and aligning universal orthologs

Searches for universal orthologs was executed using Compleasm version 0.2.5 (Huang & Li, 2023) with OrthoDB version 10 reference sequences (Kriventseva et al., 2019) for the Viridiplantae, Chlorophyta, Liliopsida, Eudicots, Fungi, Ascomycota, Basidiomycota, Metazoa, Arthropoda and Vertebrata lineages using the default settings. For duplicated universal single-copy orthologs, the ortholog that was more syntenic with the database was selected. Gene copies sharing adjacent BUSCO orthologs at greater frequency within the database were defined as more syntenic. For equally syntenic duplicates, the gene with greater sequence identity was retained. Assemblies that did not contain 90% of the BUSCO orthologs in each lineage were included in the database but dropped from the subsequent phylogenetic analysis for suboptimal quality. All identified orthologs for each gene in each lineage were aligned using MUSCLE version 5.1 (Edgar, 2021). Alignments for the Viridiplantae, Fungi, Metazoa and Arthropoda lineages were done with 16 total combinations of four parameter perturbations and four guide tree permutations to create a stratified ensemble of multiple sequence alignments (Edgar, 2021). Confidence for each column in the alignment was computed using the addconfseq flag in MUSCLE v5.

### Phylogenetic assessment

For Eudicots, Ascomycota, Basidiomycota, Arthropoda and Vertebrata lineages, aligned sites were filtered by the number of unique amino acids in the column as a proxy for rate of evolution at that site. For rate categories 2 to 15, we selected between 1,000 and 20,000 sites at 1,000 site increments. This resulted in a total of 14×20 = 280 alignments per lineage. For the Arthropoda lineage, we could only select up to 14,000 sites per category because of the relatively lower number of aligned sites. We had 14×14 = 196 total alignments for arthropods. Assemblies that had fewer than 90% BUSCO genes and aligned sites that comprised more than 10% gaps were removed. IQ-TREE version 2.1.2 (Minh et al., 2020) was used with default settings including built-in ModelFinder2 (Kalyaanamoorthy et al., 2017) to create maximum likelihood trees for every alignment. There were 198 total nuclear substitution models to test including alternate site specifications and rate category variations. A total of 280×4+196 = 1316 individual trees were created and tested in this step. Trees were assessed for taxonomic congruity by counting the number of families that descended monophyletically from a common ancestor. For the terminal and central rates 2, 8 and 14, five sets of alignments were sampled for site counts 1,000, 5,000 and 10,000. We carried out 10 independent searches on the tree space for each alignment with a different random seed, resulting in a total of 10×5 = 50 trees for 3×3 = 9 conditions in 5 lineages. The total number of trees at this stage was 50×9×5 = 2250. Each individual tree was assessed for congruity by counting the number of monophyletic families. The set of trees in each condition was assessed for entropy or degree of variation at the terminal leaves by counting the total number of unique terminal bifurcations in the set. The 5 alignment sets at site rate 8 and site count 5,000 were used to compute likelihoods for all 2,250 trees using IQ-TREE. The mean likelihood score of 5 alignments was used as the likelihood for each individual tree. Gene trees were created for the 200 longest genes in the Eudicots, Arthropoda and Vertebrata lineages.

From 5 to 75 genes were selected with increments of 5 genes at random to create 15 coalescent trees under the multi-species coalescent model in Astral-pro3 version 1.19.3.5 (Zhang & Mirarab, 2022).

### Assessing misidentified BUSCOs

For BUSCO misidentification studies, all single and duplicate BUSCO genes identified by Compleasm were first removed using scripts available on the phyca GitHub page and Compleasm was rerun on the genome set. Genes found in fragments were not considered. The curated BUSCO gene set was selected manually by looking at the frequency at which each BUSCO gene was misidentified. For each assembly, genome inflation was defined as the average number of times the BUSCO gene set was found in the assembly. Polyploid genomes shown in Supplementary Figure 1 were labeled manually according to literature through searches done by the species names. Assembly level for chromosome scale assemblies was determined by the labels assigned to the pseudomolecules.

Gene blocks were traced with all possible permutations of identified and remnant BUSCO genes up to 11 genes in length using phyca scripts. To compute CUSCO and MUSCO proportions, Remnant-Identified gene doublets were considered syntenic when they were matched in gene identity and orientation by a Identified-Identified doublet within the same lineage. Remnant-Remnant gene doublets were considered syntenic when they were matched by either a Remnant-Identified doublet or an Identified-Identified doublet. For each set of BUSCO doublets, fraction of doublets where both genes were CUSCO genes was defined as the CUSCO proportion and the fraction of doublets where both genes were MUSCO genes were defined as MUSCO proportions.

For comparisons of BUSCO gene content and syntenic distance, two assemblies of the highest and lowest N50 were selected for organisms with more than one available genome assembly from NCBI Genome. Only pairs where the difference in N50 was greater than 200Kb were considered. Assemblies with an N50 of less than 1Mb or less than 80% BUSCO content were filtered out. Systenic distance and distance matrices were computed by phyca. Exponential curves were fit using the curve_fit function from SciPy (Virtanen et al., 2020) version 1.14.1. Distance matrices were converted to newick trees using scikit-bio version 0.6.2 (https://scikit.bio).

The *Oryza alta* assembly was from Yu et al., 2021 (Yu et al., 2021) and *Oryza coarctata* was from Fornasiero et al., 2024 (Fornasiero et al., 2024). Pseudomolecules of two subgenomes of the polyploid *Oryza* species were separated through their sequence headers. All dendograms and cladograms were created using BioNick version 0.0.3 (https://pypi.org/project/BioNick/0.0.3/). The phyca website uses phylotree.js (https://phylotree.hyphy.org/) for dynamic tree visualizations.

## Supporting information

All figures and tables

## Acknowledgements

We would like to acknowledge Robert C. Edgar and Derrick Zwickl for their valuable insights and suggestions on alignment and phylogenetic methods. We acknowledge all members of the Arizona Genomics Institute and Data Diversity Lab teams for their continued support and encouragement. We acknowledge Abid Mahmood and his team for their help with the project website development. We also thank Chandler Sobel-Sorenson and the University of Arizona High Performance Computing team for maintaining and assisting us with necessary computational resources and software.

## Author information

### Contributions

RAW and MNUA conceived and planned the project. RAW, CRP and DC supervised the work. CRP reviewed the phylogenetic methods and helped design further experiments to validate the results. MNUA implemented the methods, compiled the data set, developed the algorithms and wrote the scripts and manuscript. All authors reviewed and edited the manuscript.

## Ethics declarations

### Ethics approval and consent to participate

Not applicable.

### Consent for publication

Not applicable.

## References

Alam, M. N. U., & Chowdhury, U. F. (2020). Short k-mer abundance profiles yield robust machine learning features and accurate classifiers for RNA viruses. PLoS One, 15(9), e0239381. 10.1371/journal.pone.0239381

Armstrong, J., Hickey, G., Diekhans, M., Fiddes, I. T., Novak, A. M., Deran, A., Fang, Q., Xie, D., Feng, S., Stiller, J., Genereux, D., Johnson, J., Marinescu, V. D., Alföldi, J., Harris, R. S., Lindblad-Toh, K., Haussler, D., Karlsson, E., Jarvis, E. D., … Paten, B. (2020). Progressive Cactus is a multiple-genome aligner for the thousand-genome era. Nature, 587(7833), 246–251. 10.1038/s41586-020-2871-y

Boudinot, B. E., Fikáček, M., Lieberman, Z. E., Kusy, D., Bocak, L., Mckenna, D. D., & Beutel, R. G. (2023). Systematic bias and the phylogeny of Coleoptera—A response to Cai et al. (2022) following the responses to Cai et al. (2020). Systematic Entomology, 48(2), 223–232. 10.1111/syen.12570

Carbone, I., White, J. B., Miadlikowska, J., Arnold, A. E., Miller, M. A., Kauff, F., U’Ren, J. M., May, G., & Lutzoni, F. (2017). T-BAS: Tree-Based Alignment Selector toolkit for phylogenetic-based placement, alignment downloads and metadata visualization: an example with the Pezizomycotina tree of life. Bioinformatics, 33(8), 1160–1168. 10.1093/bioinformatics/btw808

Cummins, C. A., & McInerney, J. O. (2011). A method for inferring the rate of evolution of homologous characters that can potentially improve phylogenetic inference, resolve deep divergence and correct systematic biases. Systematic Biology, 60(6), 833–844.

Cunha, T. J., de Medeiros, B. A. S., Lord, A., Sørensen, M. V., & Giribet, G. (2023). Rampant loss of universal metazoan genes revealed by a chromosome-level genome assembly of the parasitic Nematomorpha. Curr. Biol., 33(16), 3514–3521.e3514. 10.1016/j.cub.2023.07.003

Delsuc, F., Brinkmann, H., & Philippe, H. (2005). Phylogenomics and the reconstruction of the tree of life. Nature Reviews Genetics, 6(5), 361–375. 10.1038/nrg1603

Edgar, R. C. (2021). Muscle5: High-accuracy alignment ensembles enable unbiased assessments of sequence homology and phylogeny. Nature Communications, 13(1), 6968. 10.1038/s41467-022-34630-w

Ellis, E. A., Storer, C. G., & Kawahara, A. Y. (2021). De novo genome assemblies of butterflies. Gigascience, 10(6). 10.1093/gigascience/giab041

Emms, D. M., & Kelly, S. (2019). OrthoFinder: phylogenetic orthology inference for comparative genomics. Genome Biol., 20(1), 238. 10.1186/s13059-019-1832-y

Fornasiero, A., Feng, T., Al-Bader, N., Alsantely, A., Mussurova, S., Hoang, N. V., Misra, G., Zhou, Y., Fabbian, L., Mohammed, N., Rivera Serna, L., Thimma, M., Llaca, V., Parakkal, P., Kudrna, D., Copetti, D., Rajasekar, S., Lee, S., Talag, J., … Wing, R. A. (2024). Oryza genome evolution through a tetraploid lens. bioRxiv. 10.1101/2024.05.29.596369

Foster, P. G. (2004). Modeling Compositional Heterogeneity. Systematic Biology, 53(3), 485–495. 10.1080/10635150490445779

Garg, V., Bohra, A., Mascher, M., Spannagl, M., Xu, X., Bevan, M. W., Bennetzen, J. L., & Varshney, R. K. (2024). Unlocking plant genetics with telomere-to-telomere genome assemblies. Nat. Genet., 1-12. 10.1038/s41588-024-01830-7

Garsmeur, O., Schnable, J. C., Almeida, A., Jourda, C., D’Hont, A., & Freeling, M. (2014). Two evolutionarily distinct classes of paleopolyploidy. Mol. Biol. Evol., 31(2), 448–454. 10.1093/molbev/mst230

Gonçalves, P., & Gonçalves, C. (2022). Horizontal gene transfer in yeasts. Current Opinion in Genetics & Development, 76, 101950. 10.1016/j.gde.2022.101950

Gundappa, M. K., To, T.-H., Grønvold, L., Martin, S. A. M., Lien, S., Geist, J., Hazlerigg, D., Sandve, S. R., & Macqueen, D. J. (2022). Genome-wide reconstruction of rediploidization following autopolyploidization across one hundred million years of Salmonid evolution. Mol. Biol. Evol., 39(1). 10.1093/molbev/msab310

Healey, A. L., Garsmeur, O., Lovell, J. T., Shengquiang, S., Sreedasyam, A., Jenkins, J., Plott, C. B., Piperidis, N., Pompidor, N., Llaca, V., Metcalfe, C. J., Doležel, J., Cápal, P., Carlson, J. W., Hoarau, J. Y., Hervouet, C., Zini, C., Dievart, A., Lipzen, A., … D’Hont, A. (2024). The complex polyploid genome architecture of sugarcane. Nature, 628(8009), 804–810. 10.1038/s41586-024-07231-4

Heath, T. A., Hedtke, S. M., & Hillis, D. M. (2008). Taxon sampling and the accuracy of phylogenetic analyses. Journal of systematics and evolution, 46(3), 239.

Huang, N., & Li, H. (2023). compleasm: a faster and more accurate reimplementation of BUSCO. Bioinformatics, 39(10). 10.1093/bioinformatics/btad595

Jarvis, E. D., Mirarab, S., Aberer, A. J., Li, B., Houde, P., Li, C., Ho, S. Y. W., Faircloth, B. C., Nabholz, B., Howard, J. T., Suh, A., Weber, C. C., da Fonseca, R. R., Alfaro-Núñez, A., Narula, N., Liu, L., Burt, D., Ellegren, H., Edwards, S. V., … Avian Phylogenomics, C. (2015). Phylogenomic analyses data of the avian phylogenomics project. Gigascience, 4, 4. 10.1186/s13742-014-0038-1

Jian, X., Edwards, S., & Liu, L. (2019). The multispecies coalescent model outperforms concatenation across diverse phylogenomic data sets. Systematic Biology, 69, 795–812. 10.1093/sysbio/syaa008

Kalyaanamoorthy, S., Minh, B. Q., Wong, T. K. F., von Haeseler, A., & Jermiin, L. S. (2017). ModelFinder: fast model selection for accurate phylogenetic estimates. Nat. Methods, 14(6), 587–589. 10.1038/nmeth.4285

Komarova, V. A., & Lavrenchenko, L. A. (2022). Approaches to the detection of hybridization events and genetic introgression upon phylogenetic incongruence. Biol. Bull. Rev., 12(3), 240–253. 10.1134/s2079086422030045

Kostka, M., Uzlikova, M., Cepicka, I., & Flegr, J. (2008). SlowFaster, a user-friendly program for slow-fast analysis and its application on phylogeny of Blastocystis. BMC Bioinformatics, 9(1), 341. 10.1186/1471-2105-9-341

Kriventseva, E. V., Kuznetsov, D., Tegenfeldt, F., Manni, M., Dias, R., Simão, F. A., & Zdobnov, E. M. (2019). OrthoDB v10: sampling the diversity of animal, plant, fungal, protist, bacterial and viral genomes for evolutionary and functional annotations of orthologs. Nucleic Acids Res., 47(D1), D807–D811. 10.1093/nar/gky1053

Kubatko, L. S., & Degnan, J. H. (2007). Inconsistency of phylogenetic estimates from concatenated data under coalescence. Syst. Biol., 56(1), 17–24. 10.1080/10635150601146041

Lavrinienko, A., Jernfors, T., Koskimäki, J. J., Pirttilä, A. M., & Watts, P. C. (2021). Does Intraspecific Variation in rDNA Copy Number Affect Analysis of Microbial Communities? Trends in Microbiology, 29(1), 19–27. 10.1016/j.tim.2020.05.019

Le, S. Q., & Gascuel, O. (2008). An Improved General Amino Acid Replacement Matrix. Molecular Biology and Evolution, 25(7), 1307–1320. 10.1093/molbev/msn067

Levy Karin, E., Mirdita, M., & Söding, J. (2020). MetaEuk-sensitive, high-throughput gene discovery, and annotation for large-scale eukaryotic metagenomics. Microbiome, 8(1), 48. 10.1186/s40168-020-00808-x

Li, H. (2023). Protein-to-genome alignment with miniprot. Bioinformatics, 39(1), btad014.

Li, H., & Durbin, R. (2023). Genome assembly in the telomere-to-telomere era. ArXiv. http://arxiv.org/abs/2308.07877

Liu, J., Shi, C., Shi, C.-C., Li, W., Zhang, Q.-J., Zhang, Y., Li, K., Lu, H.-F., Shi, C., Zhu, S.-T., Xiao, Z.-Y., Nan, H., Yue, Y., Zhu, X.-G., Wu, Y., Hong, X.-N., Fan, G.-Y., Tong, Y., Zhang, D., … Gao, L.-Z. (2020). The chromosome-based rubber tree genome provides new insights into spurge genome evolution and rubber biosynthesis. Mol. Plant, 13(2), 336–350. 10.1016/j.molp.2019.10.017

Lofgren, L. A., Uehling, J. K., Branco, S., Bruns, T. D., Martin, F., & Kennedy, P. G. (2019). Genome-based estimates of fungal rDNA copy number variation across phylogenetic scales and ecological lifestyles. Molecular Ecology, 28(4), 721–730. 10.1111/mec.14995

Luo, J., Chen, J., Guo, W., Yang, Z., Lim, K.-J., & Wang, Z. (2022). Correction: Luo et al. Reassessment of Annamocarya sinesis (Carya sinensis) Taxonomy through Concatenation and Coalescence Phylogenetic Analysis. Plants 2022, 11, 52. Plants, 11(23), 3282. 10.3390/plants11233282

Manni, M., Berkeley, M. R., Seppey, M., & Zdobnov, E. M. (2021). BUSCO: Assessing Genomic Data Quality and Beyond. Current Protocols, 1(12). 10.1002/cpz1.323

Mansfeld, B. N., Boyher, A., Berry, J. C., Wilson, M., Ou, S., Polydore, S., Michael, T. P., Fahlgren, N., & Bart, R. S. (2021). Large structural variations in the haplotype-resolved African cassava genome. Plant J., 108(6), 1830–1848. 10.1111/tpj.15543

Matschiner, M., Böhne, A., Ronco, F., & Salzburger, W. (2020). The genomic timeline of cichlid fish diversification across continents. Nature Communications, 11(1), 5895. 10.1038/s41467-020-17827-9

Minh, B. Q., Schmidt, H. A., Chernomor, O., Schrempf, D., Woodhams, M. D., von Haeseler, A., & Lanfear, R. (2020). IQ-TREE 2: New Models and Efficient Methods for Phylogenetic Inference in the Genomic Era. Mol. Biol. Evol., 37(5), 1530–1534. 10.1093/molbev/msaa015

Misof, B., Liu, S., Meusemann, K., Peters, R. S., Donath, A., Mayer, C., Frandsen, P. B., Ware, J., Flouri, T., Beutel, R. G., Niehuis, O., Petersen, M., Izquierdo-Carrasco, F., Wappler, T., Rust, J., Aberer, A. J., Aspöck, U., Aspöck, H., Bartel, D., … Zhou, X. (2014). Phylogenomics resolves the timing and pattern of insect evolution. Science, 346(6210), 763–767. 10.1126/science.1257570

Naranjo-Ortiz, M. A., & Gabaldón, T. (2020). Fungal evolution: cellular, genomic and metabolic complexity. Biological Reviews, 95(5), 1198–1232. 10.1111/brv.12605

Nasrallah, C. A., Mathews, D. H., & Huelsenbeck, J. P. (2010). Quantifying the Impact of Dependent Evolution among Sites in Phylogenetic Inference. Systematic Biology, 60(1), 60–73. 10.1093/sysbio/syq074

Pisani, D. (2004). Identifying and removing fast-evolving sites using compatibility analysis: an example from the Arthropoda. Systematic Biology, 53(6), 978–989.

Ran, J.-H., Shen, T.-T., Wang, M.-M., & Wang, X.-Q. (2018). Phylogenomics resolves the deep phylogeny of seed plants and indicates partial convergent or homoplastic evolution between Gnetales and angiosperms. Proc. Biol. Sci., 285(1881). 10.1098/rspb.2018.1012

Rangel, L. T., & Fournier, G. P. (2023). Fast-Evolving Alignment Sites Are Highly Informative for Reconstructions of Deep Tree of Life Phylogenies. Microorganisms, 11(10), 2499. https://www.mdpi.com/2076-2607/11/10/2499

Rautiainen, M., Nurk, S., Walenz, B. P., Logsdon, G. A., Porubsky, D., Rhie, A., Eichler, E. E., Phillippy, A. M., & Koren, S. (2023). Telomere-to-telomere assembly of diploid chromosomes with Verkko. Nat. Biotechnol., 41(10), 1474–1482. 10.1038/s41587-023-01662-6

Reuscher, S., Furuta, T., Bessho-Uehara, K., Cosi, M., Jena, K. K., Toyoda, A., Fujiyama, A., Kurata, N., & Ashikari, M. (2018). Assembling the genome of the African wild rice Oryza longistaminata by exploiting synteny in closely related Oryza species. Commun Biol, 1, 162. 10.1038/s42003-018-0171-y

Ronco, F., Matschiner, M., Böhne, A., Boila, A., Büscher, H. H., El Taher, A., Indermaur, A., Malinsky, M., Ricci, V., Kahmen, A., Jentoft, S., & Salzburger, W. (2021). Drivers and dynamics of a massive adaptive radiation in cichlid fishes. Nature, 589(7840), 76–81. 10.1038/s41586-020-2930-4

Rosenberg, M. S., & Kumar, S. (2001). Incomplete taxon sampling is not a problem for phylogenetic inference. Proceedings of the National Academy of Sciences, 98(19), 10751–10756. doi:10.1073/pnas.191248498

Sahbou, A.-E., Iraqi, D., Mentag, R., & Khayi, S. (2022). BuscoPhylo: a webserver for Busco-based phylogenomic analysis for non-specialists. Sci. Rep., 12(1), 17352. 10.1038/s41598-022-22461-0

Sayers, E. W., Beck, J., Bolton, E. E., Brister, J. R., Chan, J., Comeau, D. C., Connor, R., DiCuccio, M., Farrell, C. M., Feldgarden, M., Fine, A. M., Funk, K., Hatcher, E., Hoeppner, M., Kane, M., Kannan, S., Katz, K. S., Kelly, C., Klimke, W., … Sherry, S. T. (2024). Database resources of the National Center for Biotechnology Information. Nucleic Acids Res, 52(D1), D33–d43. 10.1093/nar/gkad1044

Sayers, E. W., Bolton, E. E., Brister, J. R., Canese, K., Chan, J., Comeau, D. C., Connor, R., Funk, K., Kelly, C., Kim, S., Madej, T., Marchler-Bauer, A., Lanczycki, C., Lathrop, S., Lu, Z., Thibaud-Nissen, F., Murphy, T., Phan, L., Skripchenko, Y., … Sherry, S. T. (2022). Database resources of the national center for biotechnology information. Nucleic Acids Res., 50(D1), D20–D26. 10.1093/nar/gkab1112

Sayers, E. W., Bolton, E. E., Brister, J. R., Canese, K., Chan, J., Comeau, D. C., Farrell, C. M., Feldgarden, M., Fine, A. M., Funk, K., Hatcher, E., Kannan, S., Kelly, C., Kim, S., Klimke, W., Landrum, M. J., Lathrop, S., Lu, Z., Madden, T. L., … Sherry, S. T. (2023). Database resources of the National Center for Biotechnology Information in 2023. Nucleic Acids Res, 51(D1), D29–d38. 10.1093/nar/gkac1032

Schrempf, D., & Szöllősi, G. (2020). The sources of phylogenetic conflicts. Phylogenetics in the genomic era, 3–1. https://hal.science/hal-02535482/file/chapter_3.1_Schrempf_Szollosi.pdf

Seppey, M., Manni, M., & Zdobnov, E. M. (2019). BUSCO: assessing genome assembly and annotation completeness. Gene prediction: methods and protocols, 227–245.

Superson, A., & Battistuzzi, F. (2022). Exclusion of fast evolving genes or fast evolving sites produces different archaean phylogenies. Molecular Phylogenetics and Evolution, 170, 107438.

Susko, E., & Roger, A. (2021). Long Branch Attraction Biases in Phylogenetics. Syst. Biol. 10.1093/sysbio/syab001

Timilsena, P. R., Wafula, E. K., Barrett, C. F., Ayyampalayam, S., McNeal, J. R., Rentsch, J. D., McKain, M. R., Heyduk, K., Harkess, A., Villegente, M., Conran, J. G., Illing, N., Fogliani, B., Ané, C., Pires, J. C., Davis, J. I., Zomlefer, W. B., Stevenson, D. W., Graham, S. W., … dePamphilis, C. W. (2022). Phylogenomic resolution of order- and family-level monocot relationships using 602 single-copy nuclear genes and 1375 BUSCO genes [Original Research]. Frontiers in Plant Science, 13. 10.3389/fpls.2022.876779

Van Damme, K., Cornetti, L., Fields, P. D., & Ebert, D. (2022). Whole-genome phylogenetic reconstruction as a powerful tool to reveal homoplasy and ancient rapid radiation in waterflea evolution. Syst. Biol., 71(4), 777–787. 10.1093/sysbio/syab094

Venditti, C., Meade, A., & Pagel, M. (2006). Detecting the node-density artifact in phylogeny reconstruction. Syst. Biol., 55(4), 637–643. 10.1080/10635150600865567

Virtanen, P., Gommers, R., Oliphant, T. E., Haberland, M., Reddy, T., Cournapeau, D., Burovski, E., Peterson, P., Weckesser, W., Bright, J., van der Walt, S. J., Brett, M., Wilson, J., Millman, K. J., Mayorov, N., Nelson, A. R. J., Jones, E., Kern, R., Larson, E., … SciPy, C. (2020). SciPy 1.0: fundamental algorithms for scientific computing in Python. Nature methods, 17(3), 261–272. 10.1038/s41592-019-0686-2

Wheeler, T. J., & Eddy, S. R. (2013). nhmmer: DNA homology search with profile HMMs. Bioinformatics, 29(19), 2487–2489. 10.1093/bioinformatics/btt403

Wighard, S. S., Athanasouli, M., Witte, H., Rödelsperger, C., & Sommer, R. J. (2022). A New Hope: A hermaphroditic nematode enables analysis of a recent whole genome duplication event. Genome Biol. Evol., 14(12). 10.1093/gbe/evac169

Yan, Z., Smith, M. L., Du, P., Hahn, M. W., & Nakhleh, L. (2021). Species tree inference methods intended to deal with incomplete lineage sorting are robust to the presence of paralogs. Syst. Biol., 71, 367–381. 10.1093/sysbio/syab056

Yang, Z. (1994). Maximum likelihood phylogenetic estimation from DNA sequences with variable rates over sites: approximate methods. Journal of Molecular evolution, 39, 306–314.

Yang, Z. (1996). Among-site rate variation and its impact on phylogenetic analyses. Trends in ecology & evolution, 11(9), 367–372.

Yang, Z. (2006). Computational molecular evolution. OUP Oxford.

Yu, H., Lin, T., Meng, X., Du, H., Zhang, J., Liu, G., Chen, M., Jing, Y., Kou, L., Li, X., Gao, Q., Liang, Y., Liu, X., Fan, Z., Liang, Y., Cheng, Z., Chen, M., Tian, Z., Wang, Y., … Li, J. (2021). A route to de novo domestication of wild allotetraploid rice. Cell, 184(5), 1156–1170.e1114. 10.1016/j.cell.2021.01.013

Zhang, C., & Mirarab, S. (2022). ASTRAL-Pro 2: ultrafast species tree reconstruction from multi-copy gene family trees. Bioinformatics, 38(21), 4949–4950. 10.1093/bioinformatics/btac620

