## Supplemental methods and notes for "Universal orthologs infer deep phylogenies and improve genome quality assessments"

**This PDF file includes:**

Supplementary Text

Figs. S1 to S14

References

**Other Supplementary Materials for this manuscript include the following:**

Tables S1 to S5

Supplementary Text

**1. Methods for miscellaneous numerical figures**

In January 2024, the data set had 1327 viridiplantae assemblies, 4027 fungi assemblies, and 5744 animal assemblies representing the bulk of the species in NCBI genome. They added up to 11,098.

There were 34 phyla, 146 classes, 548 orders and 1,878 families. They added up to 2,606.

Of the 543 families tested, 275 were claimed ‘suitable’ in the abstract since these families were not resolved as monophyletic in all 150 trees at 10,000 alignment length. For the 275 families in rate 2 and alignment length 10,000, the mean incidence of monophyly was 26.65 and at rate 14 and alignment length 10,000, it was 38.57. This was described as a max 23.84% increase in taxonomic concordance in the abstract.

For terminal leaf bifurcations in alignments of length 10,000, the largest difference caused by rates was observed in the Vertebrata lineage going from 3,000 unique bifurcations in rate 2, to 648 in rate 14. This was described as a max 78.40% reduction in terminal variability. The smallest difference was seen in the Basidiomycota lineage, going from 611 to 329. This was described as a minimum 46.15% reduction in terminal variability.

The 6 non-overlapping lineages of plants, fungi and animals, i.e., Liliopsida, Eudicots, Ascomycota, Basidiomycota, Arthropoda and Vertebrata were used to compute the duplication percentiles. For example, the mean percentile of duplicated BUSCOs in the Liliopsida lineage was 19.78% and in Eudicots, it was 13.34%. The average, 16.57%, was used in the first section of the results.

CUSCO (Curated Universal Single Copy Orthologs) gene percentiles were used in the reported Mann-Whitney U tests for differences in CUSCO completeness and duplication levels because of their greater sensitivity over BUSCOs.

**2. Computation and commands list:**

Computation was split between the Arizona Genomics Institute’s Eagi and Pac clusters and the University of Arizona’s central HPC clusters Puma and Ocelote. Eagi consists of 5 supermicro nodes each running dual 64 core (256 threads total) AMD Epyc processors with 1TB RAM per node. Computation on Pac was limited to its 2 high performance nodes having dual 32 core Intel processors and 1TB of RAM each. Job scheduling and parallelization was primarily managed by Slurm (<https://slurm.schedmd.com>) and GNU parallel (<https://www.gnu.org/software/parallel/>). Puma and Ocelote have variable node configurations and were accessed by batch and array job submissions only through Slurm. Jobs were split manually across these clusters based on expected computation time and CPU+memory requirements. The longest CPU-time demands were seen during the ensemble multiple sequence alignments in MUSCLE v5 (Edgar, 2021) and tree searches in IQ-TREE v2.1.2 (Minh et al., 2020), each totaling over a million CPU hours. The largest memory requirements were observed during whole-genome alignment post-processing.

Downloads were initiated by curl using the following command:

curl -OJX GET \"[URL]\" -H \"Accept: application/zip\"

Command for assembly statistics:

assembly-stats -t assembly.fna > assembly.stats

Command for stratified alignments with MUSCLE v5:

muscle -threads 32 -align unaligned.fa -output aln_stratified.efa -stratified
muscle -addconfseq aln_stratified.efa -output aln_sc.efa

muscle -maxcc aln_stratified.efa -output aln_sbest.efa

Column confidence of alignment ensemble:

muscle -addconfseq stratified.efa -output sc.efa

Command for extracting the first alignment from an ensemble:

sed '/^<abc.*$/,$d' aln_stratified.efa > sfirst.efa

Example command for Compleasm run:

compleasm run -t 4 -a assembly.fa -l lineage -o output

Converting Newick to Tabular formats:

cat tree.nw | sed 's/(//g' | sed 's/)//g' | sed 's/,/\n/g' | sed 's/:/\t/g' | tac > output

Count of miniport hits:

awk '$3 == "mRNA"' folder/fungi_odb10/miniprot_output.gff | wc -l

Example IQ-TREE command for tree search:

iqtree2 -s alignment.afa -pre tree.nw -msub nuclear -nt AUTO --threads-max 4 -b 100 -alrt 1000

IQ-TREE command for recomputing likelihood on fixed tree:

iqtree2 -s alignment.afa -te tree.nw -pre output -m LG+F+R6 -nt 20

Astral-Pro3 version 1.19.3.5 (Zhang & Mirarab, 2022) command for computing coalescent trees from gene trees:

ASTER-Linux/bin/astral-pro -t 4 -i treeset.nw -o output.nw

Scripts and functions for more sophisticated tasks are documented with the phyca software toolkit on github at <https://github.com/DeadlineWasYesterday/phyca> and also the legacy software <https://github.com/DeadlineWasYesterday/phyloBUSCO>.

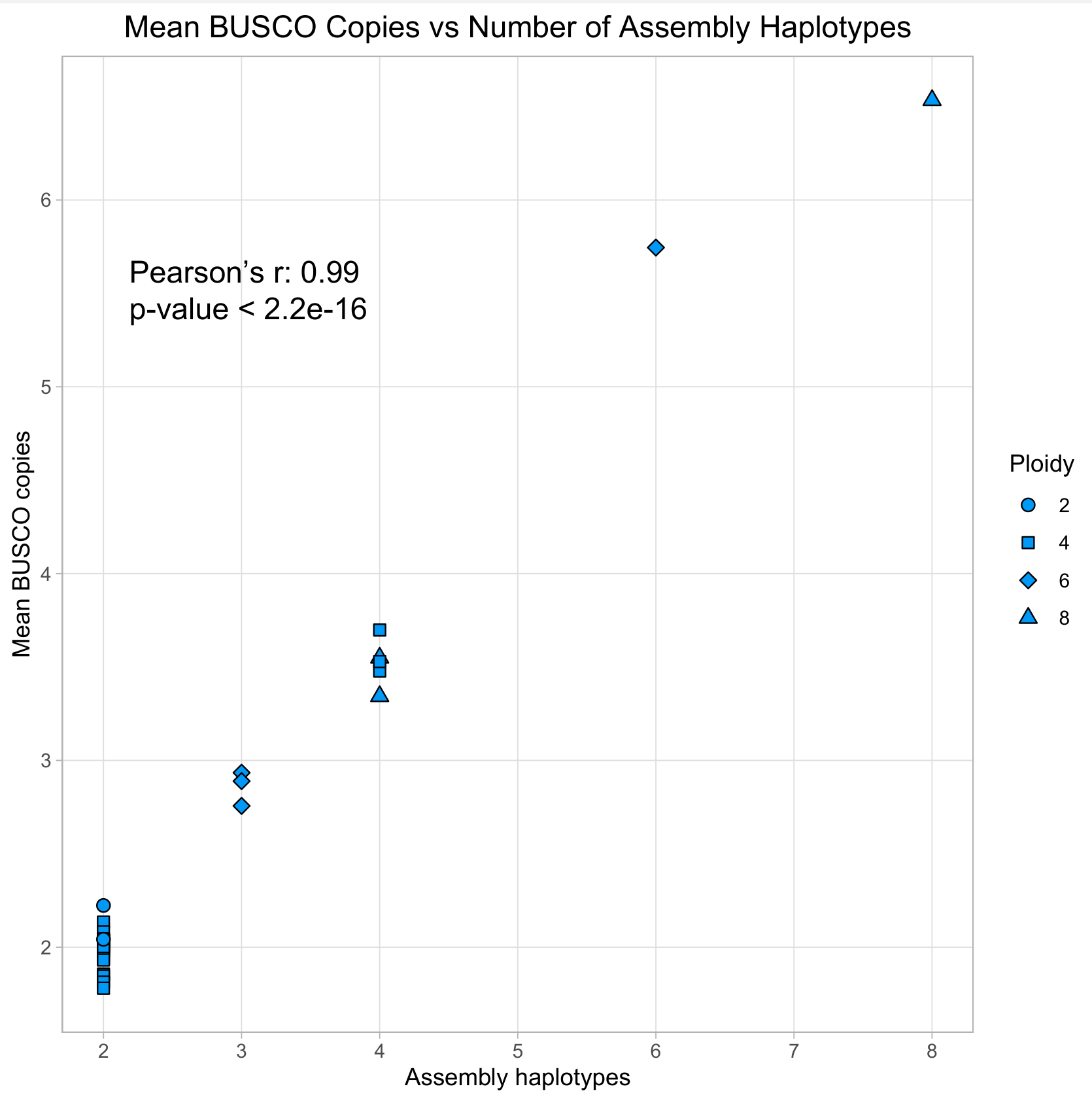

Fig. S1.

Mean BUSCO copy numbers correlate with heterozygous and polyploid genomes assembled at higher levels. Complex genomes with higher ploidy are often assembled in sets of pseudomolecules, and each set could represent either a subgenome or a haploid chromosome set. For example, wheat is hexaploid with three highly homozygous subgenomes. Hence, a high quality wheat assembly generally has three sets of pseudomolecules to represent its three subgenomes.

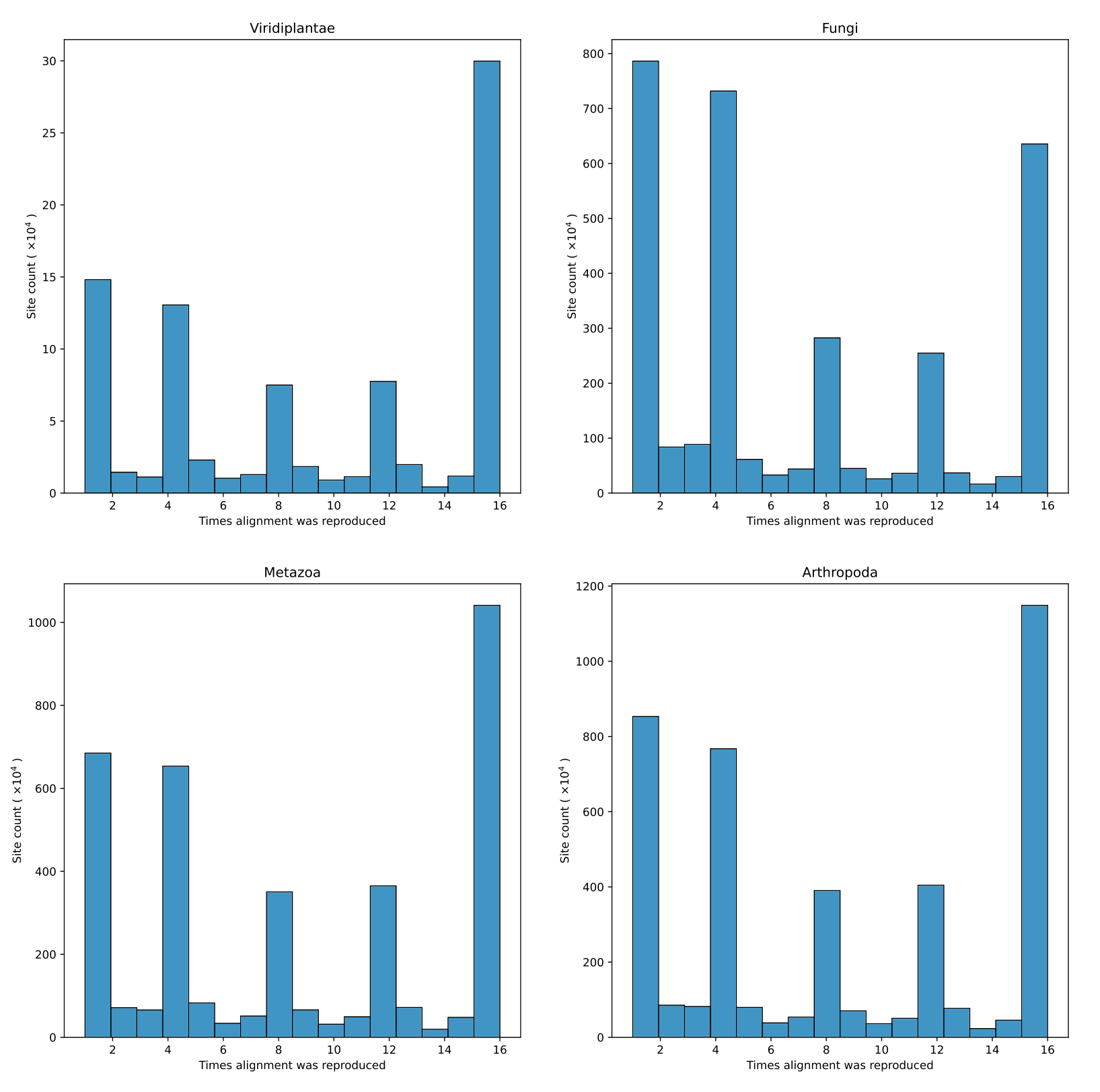

Fig. S2.

Multiple sequence alignments vary based on alignment algorithm parameters. Stratified alignments are generated by MUSCLE v5 by tuning the HMM and guide tree parameters. We created stratified alignments with 4 HMM perturbations and 4 guide tree permutations, resulting in a set of 16 alignments. From the 16 alignments, the figure shows the number of times alignment columns were reproduced.

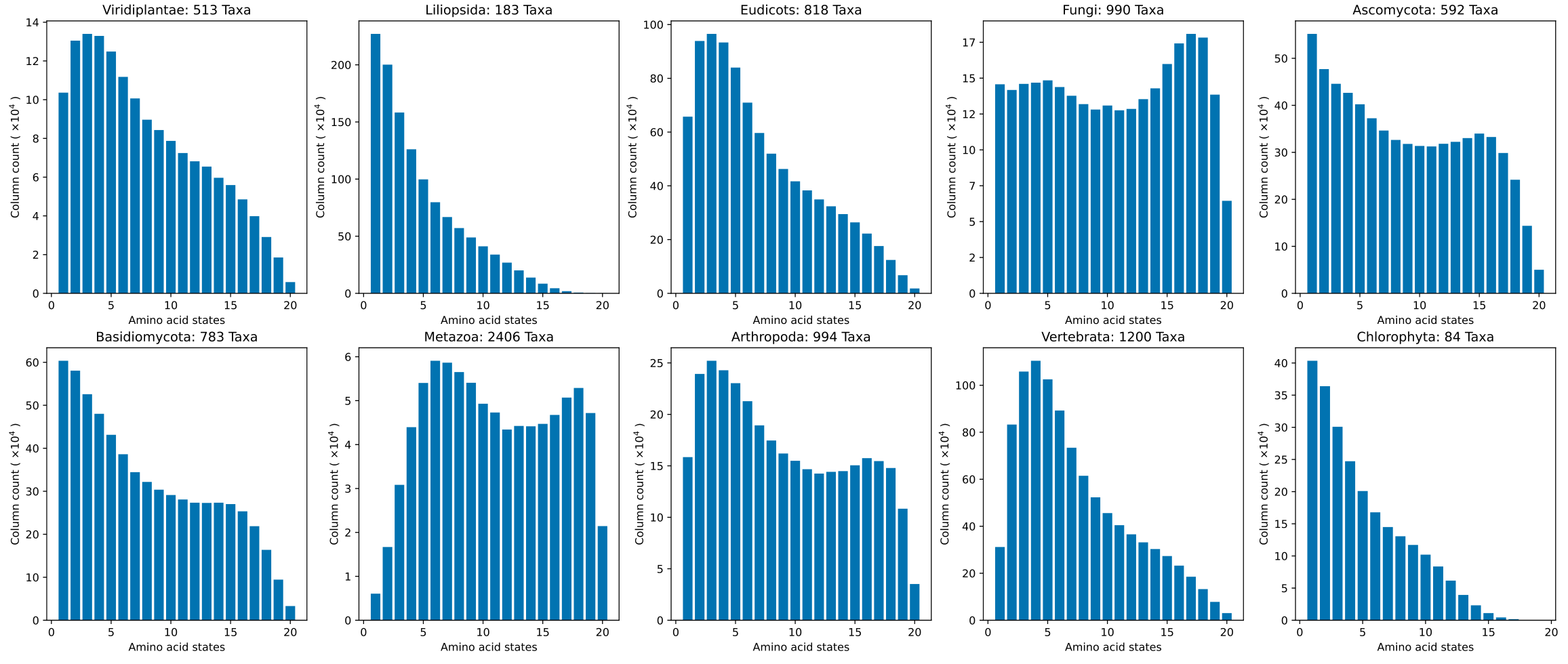

Fig. S3.

Amino acid state frequency spectra for 10 lineages. Some lineages are more divergent than others. The number of amino acid residues in alignment columns were enumerated and plotted. Gap characters were not considered. The amino acid counts serve as a proxy for evolutionary rate of the site of the alignment column.

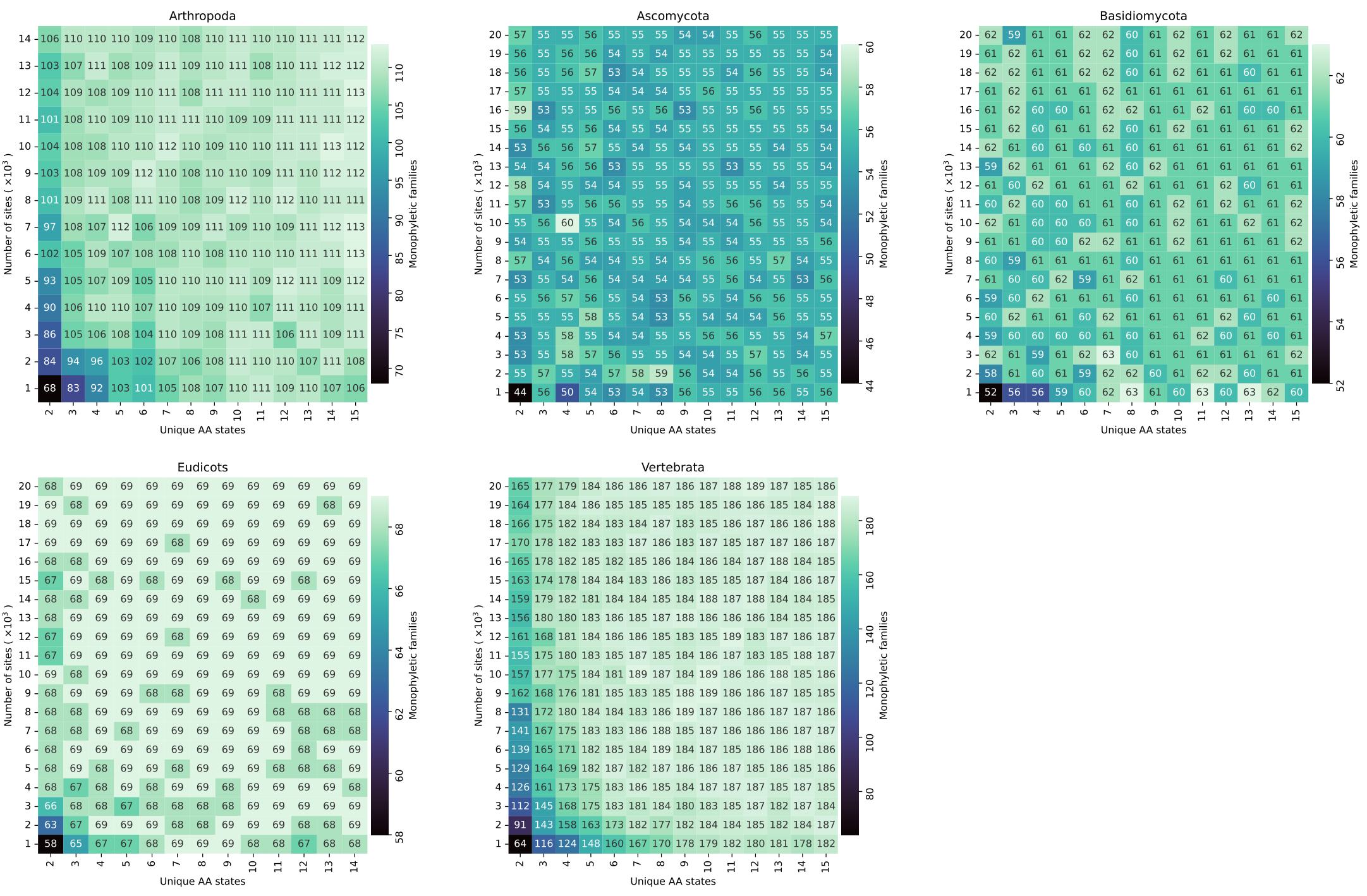

Fig. S4.

Taxonomic concordance of trees as a function of site evolution rates and alignment lengths. The number of monophyletic families were enumerated at varying rate and alignment length conditions. Trees with higher numbers of monophyletic families are considered to be more taxonomically congruent.

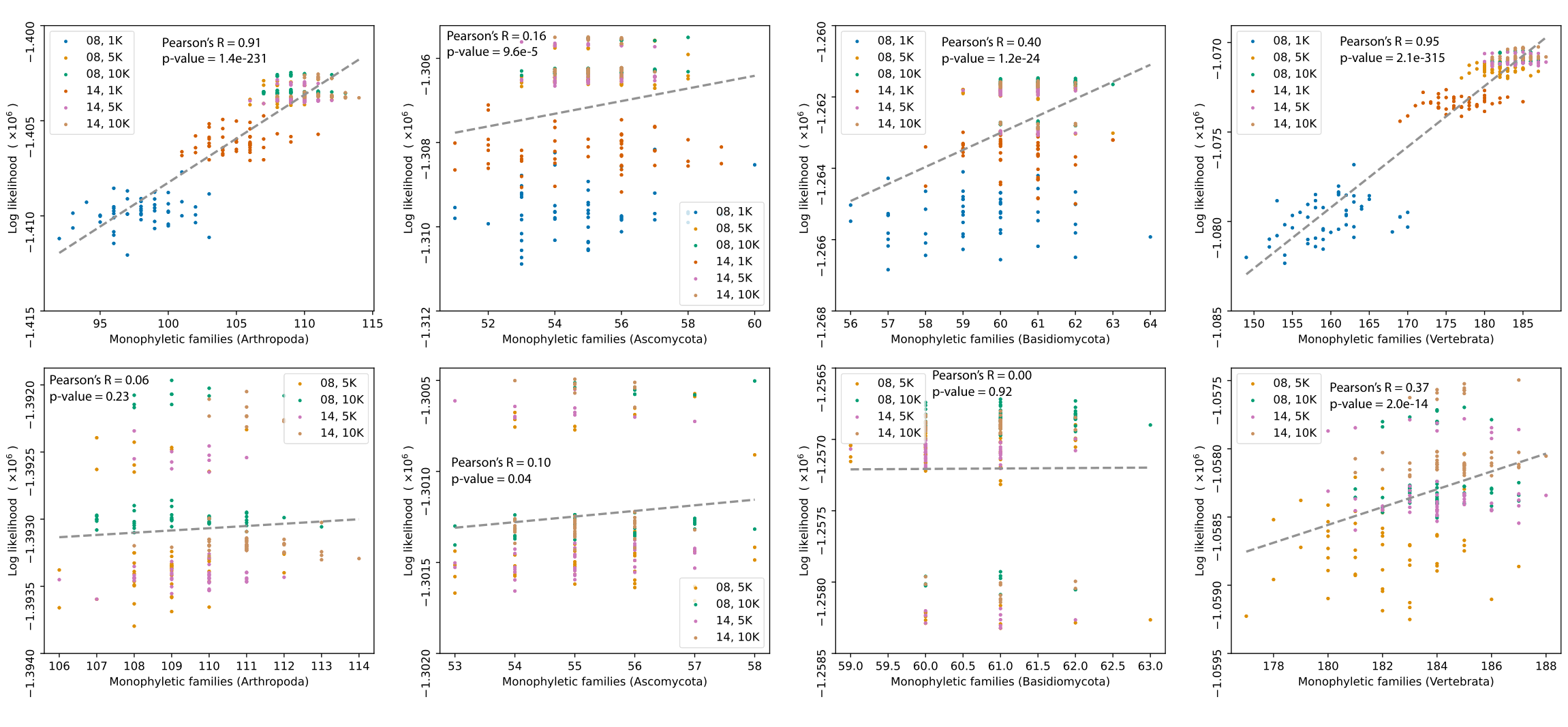

Fig. S5.
Correlations between tree likelihood and taxonomic concordance under different rate and site configurations. The top row of graphs display tree likelihoods from 6 conditions, whereas the bottom row of graphs only show the 4 conditions with the highest rates and alignment lengths.

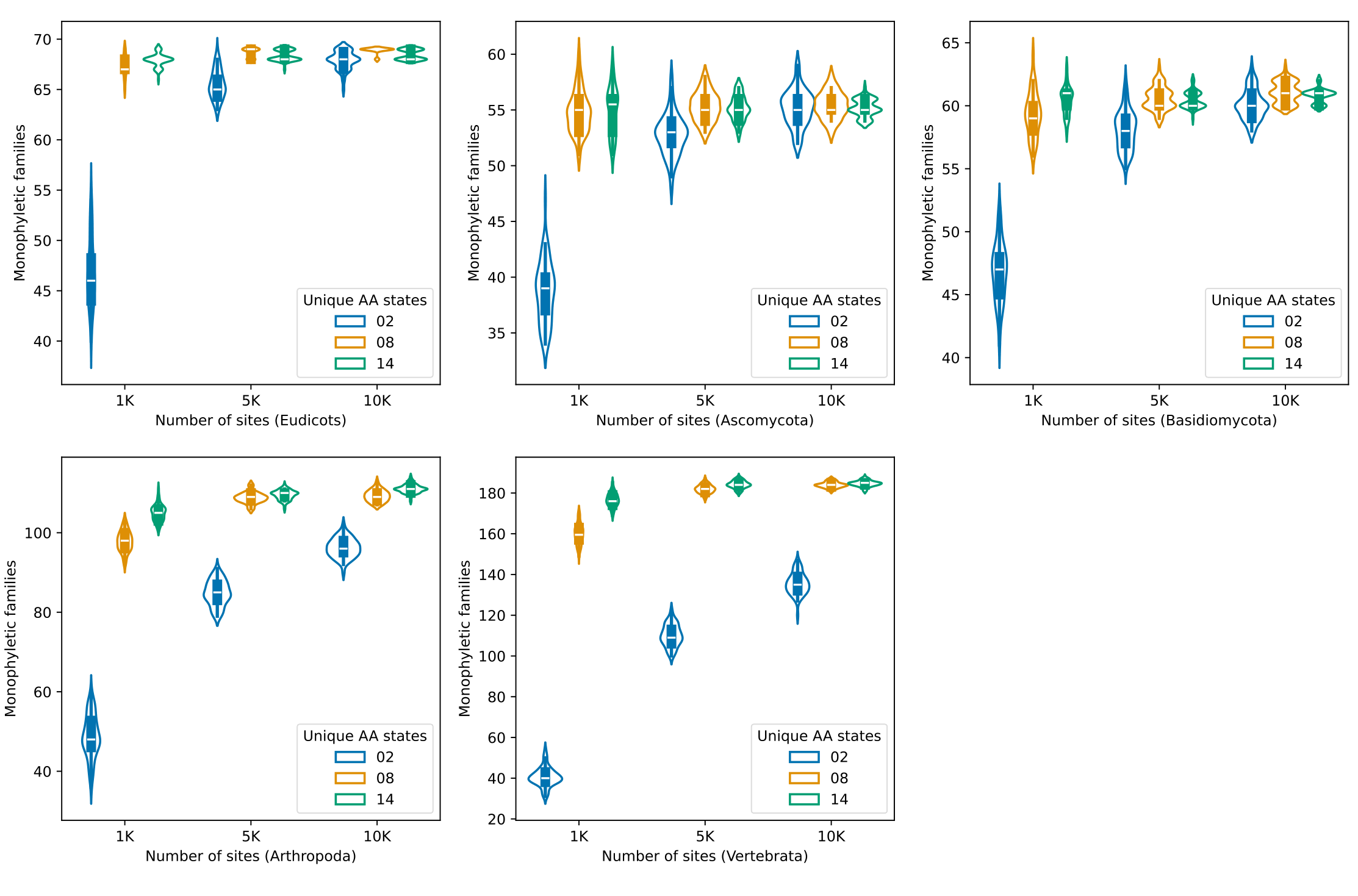

Fig. S6.

Variations in taxonomic concordance within trees created under the same rate and site conditions. For eudicots, arthropods and vertebrates, tree sets become more taxonomically congruent with less variation at higher rates and alignment lengths. For ascomycetes and basidiomycetes, only variation in taxonomic congruence is reduced at higher rates and alignment lengths.

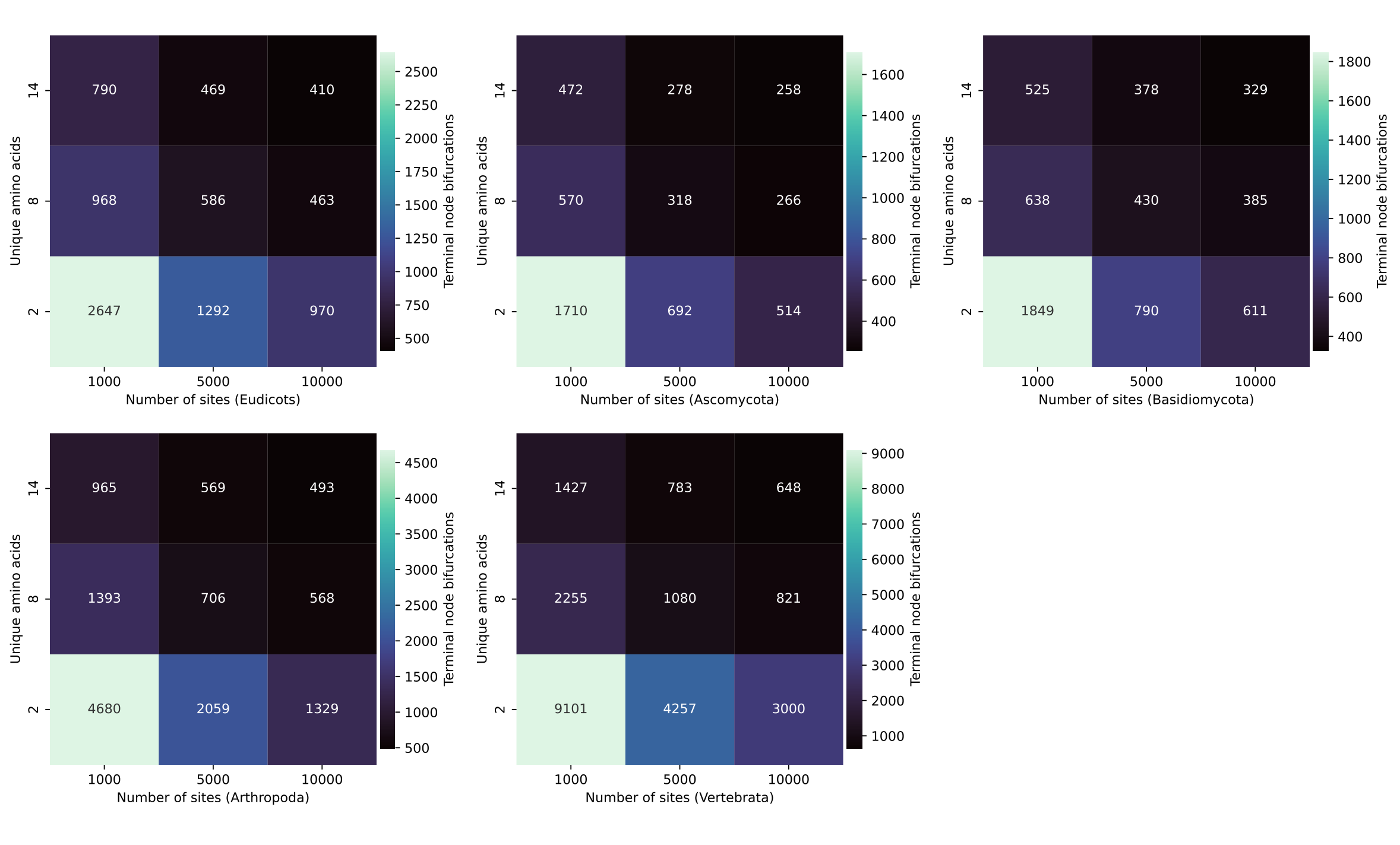

Fig. S7.

Counts of unique terminal leaf bifurcations in 9 conditions for 5 tested lineages. Higher rates and longer alignments produce less terminally variable tree sets.

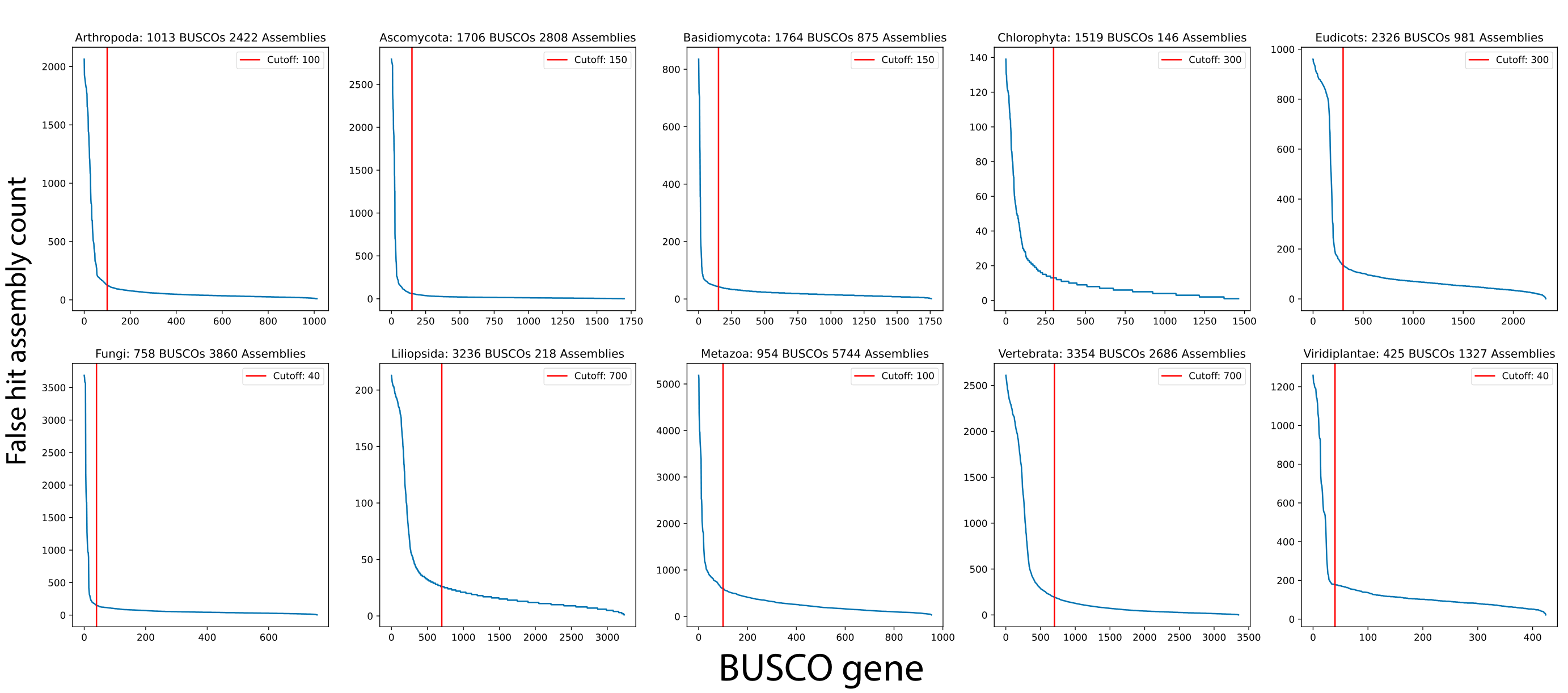

Fig. S8.

Exclusion of BUSCO genes that are misidentified at far greater propensity than others. In all tested lineages, a subset of BUSCO genes was found to be misannotated a higher number of times compared to others. These genes were removed from BUSCO search results to improve annotation accuracy.

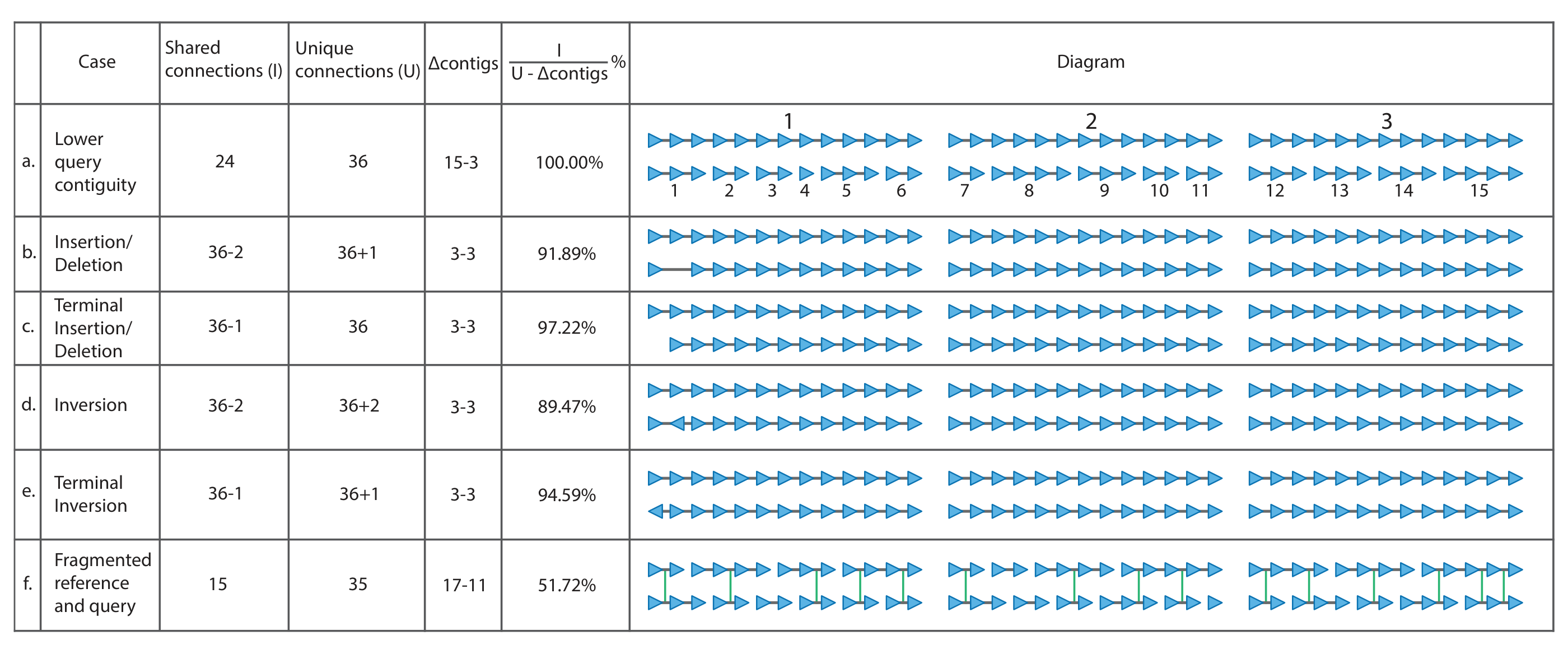

Fig. S9.
Demonstration of using the proposed BUSCO syntenic identity metric and how it is affected by variations in assembly contiguity, insertions, deletions, inversion and fragmentation in both assemblies.

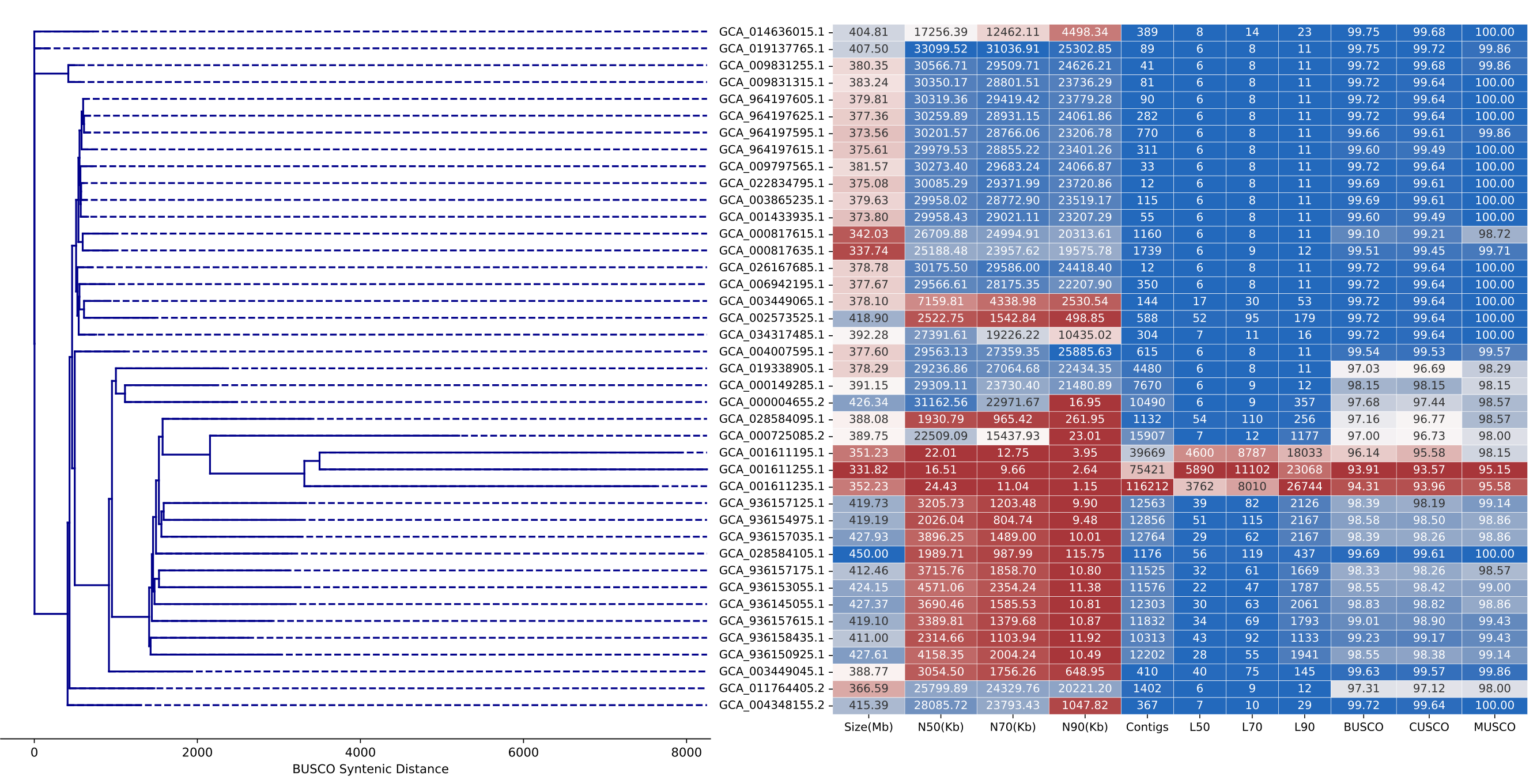

Fig. S10.

BUSCO synteny used as a distance metric to represent the magnitude of differences between 41 *Oryza sativa* assemblies.

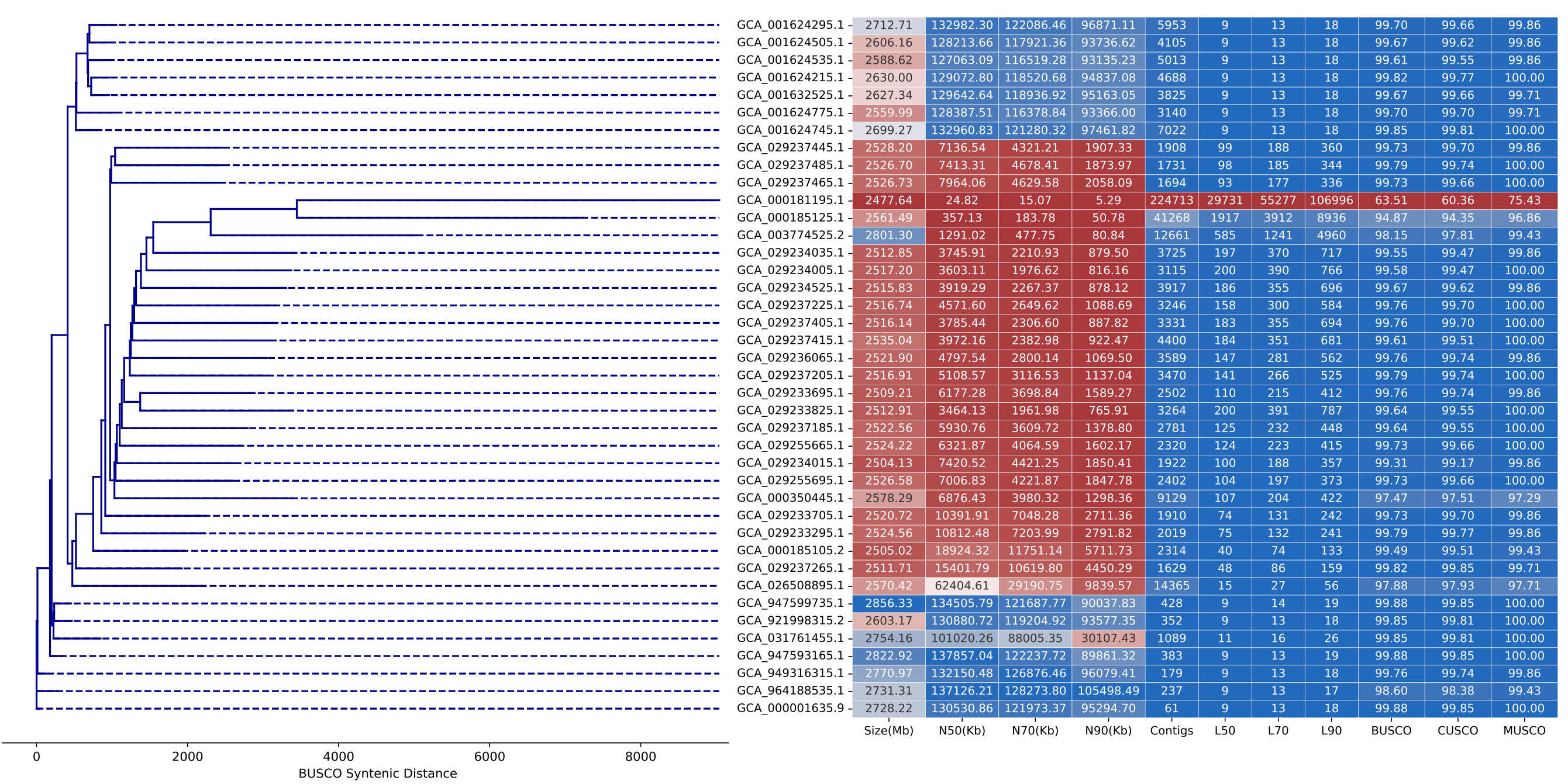

Fig. S11.

BUSCO synteny used as a distance metric to represent the magnitude of differences between 41 *Mus musculus* assemblies.

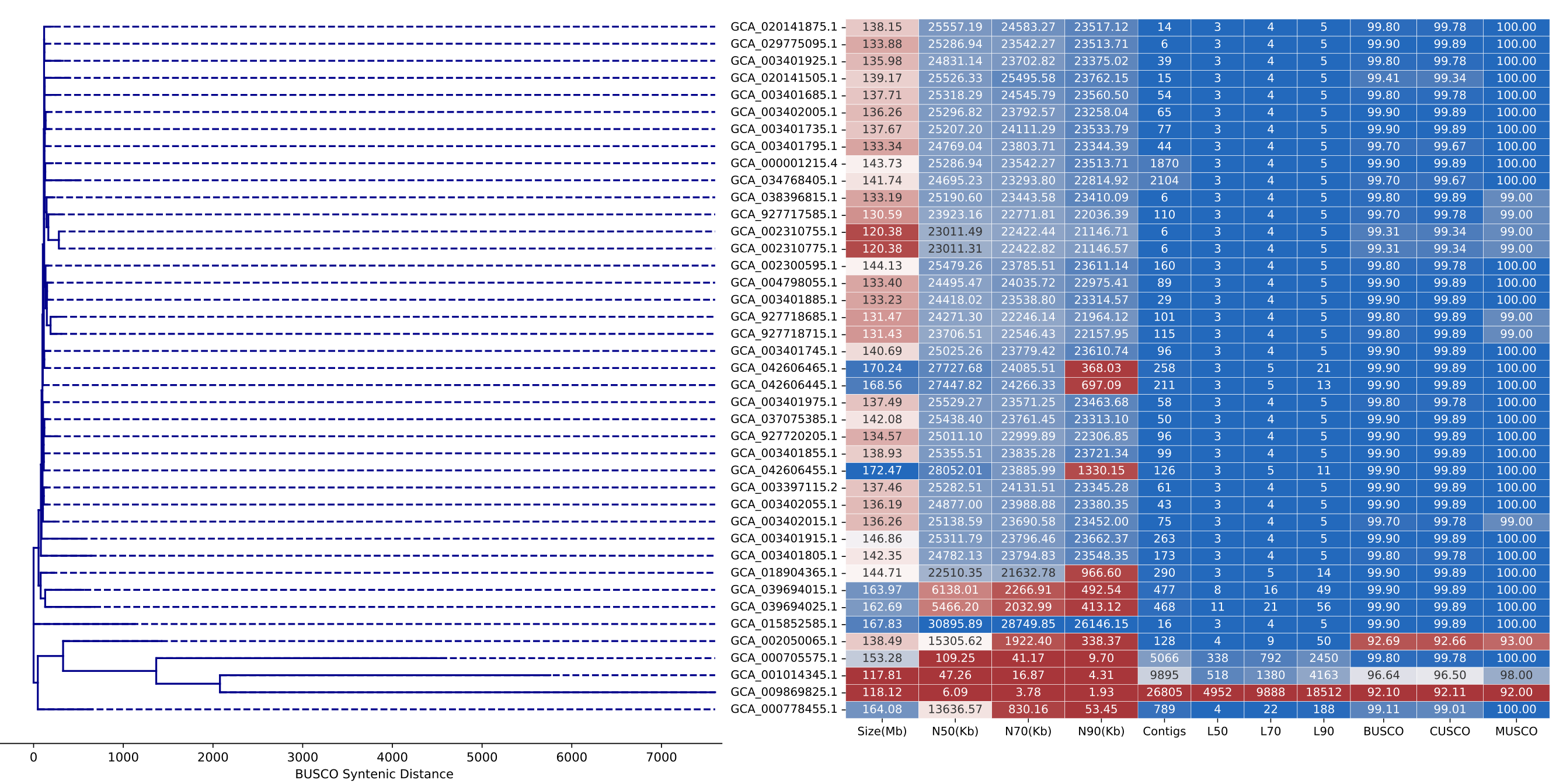

Fig. S12.

BUSCO synteny used as a distance metric to represent the magnitude of differences between 41 *Drosophila melanogaster* assemblies.

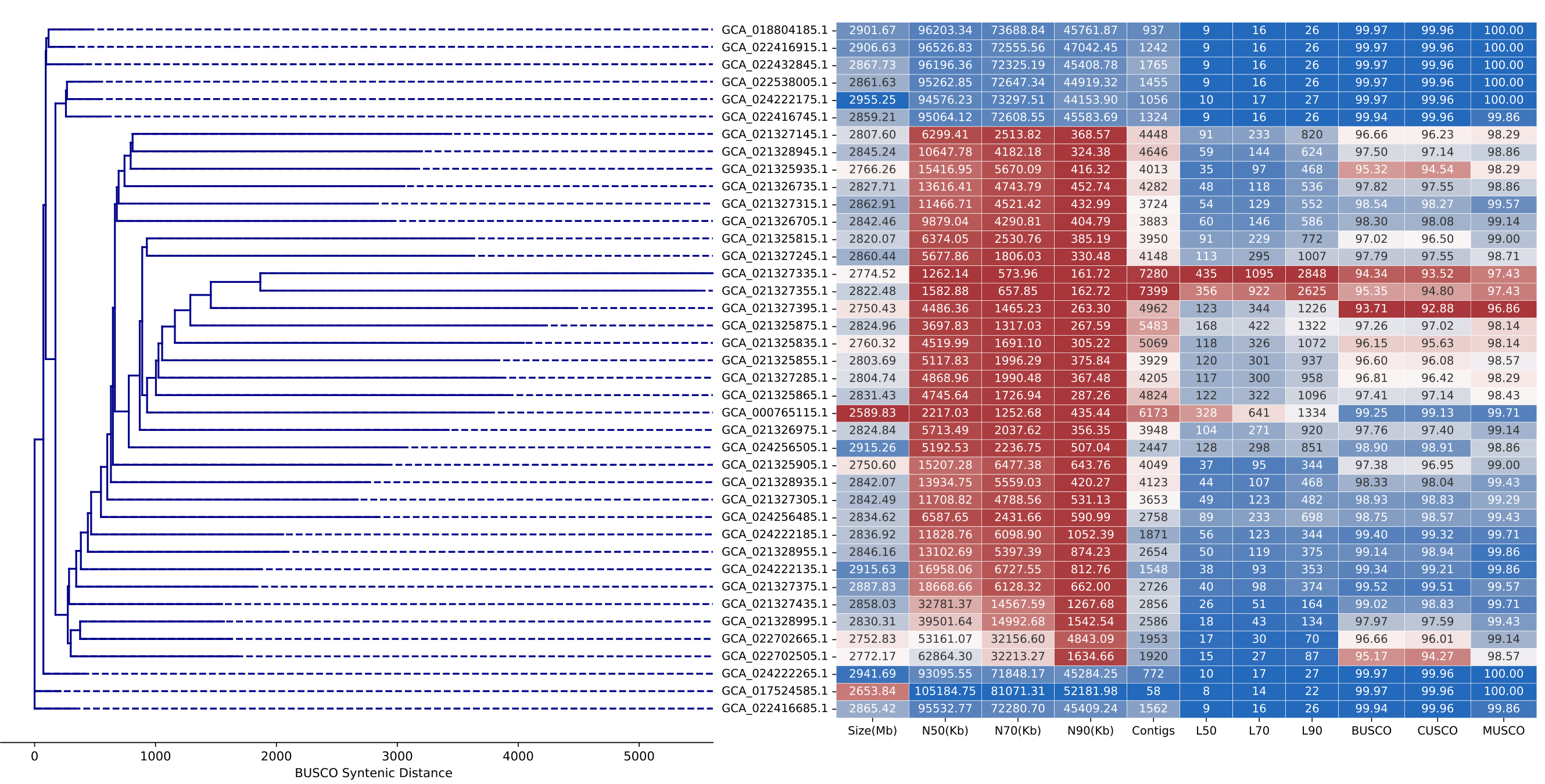

Fig. S13.

BUSCO synteny used as a distance metric to represent the magnitude of differences between 41 *Ovis aries* assemblies.

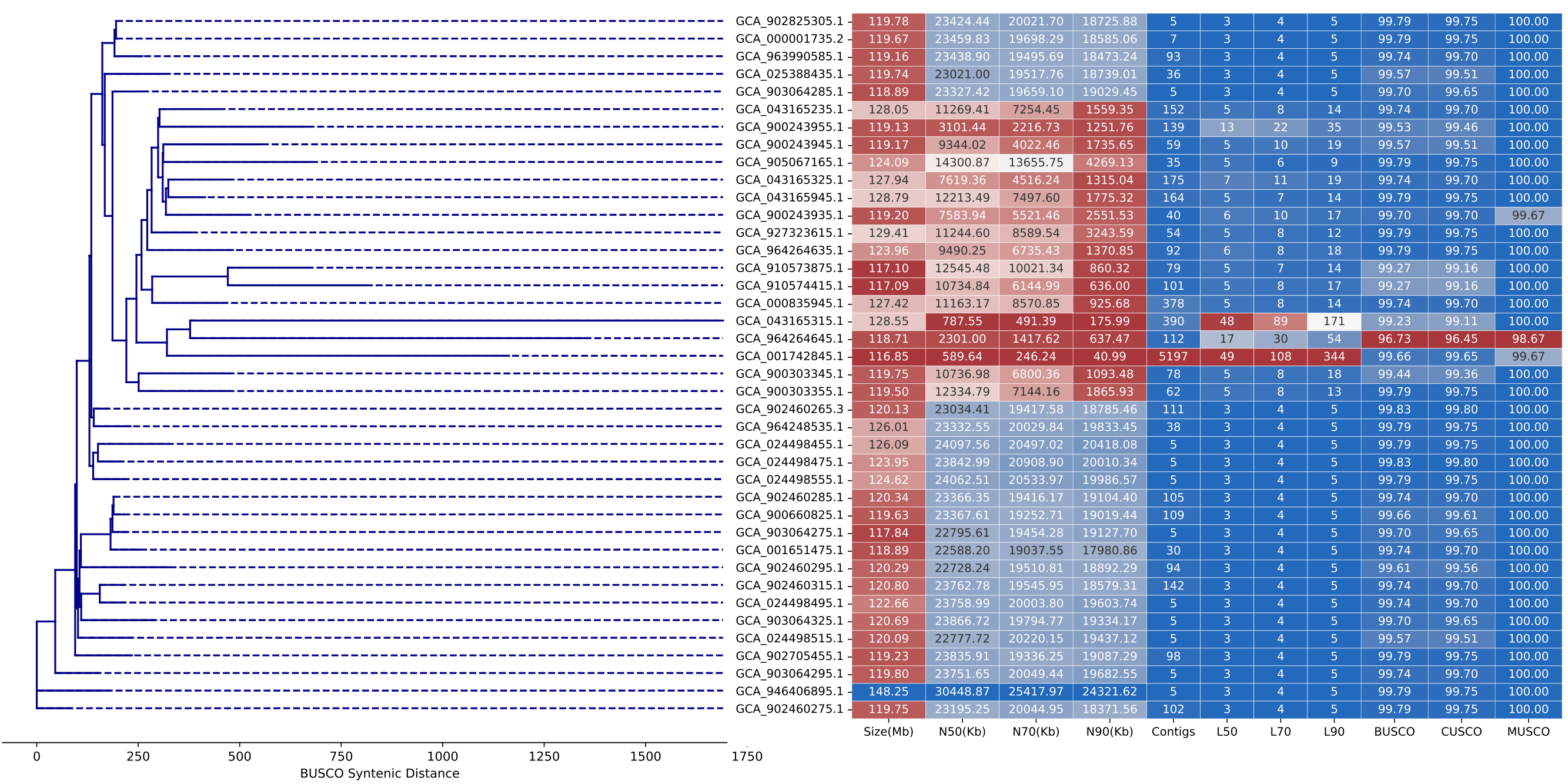

Fig. S14.

BUSCO synteny used as a distance metric to represent the magnitude of differences between 41 *Arabidopsis thaliana* assemblies.

Works cited:

Edgar, R. C. (2021). Muscle5: High-accuracy alignment ensembles enable unbiased assessments of sequence homology and phylogeny. *Nature Communications*, *13*(1), 6968. <https://doi.org/10.1038/s41467-022-34630-w>

Minh, B. Q., Schmidt, H. A., Chernomor, O., Schrempf, D., Woodhams, M. D., von Haeseler, A., & Lanfear, R. (2020). IQ-TREE 2: New Models and Efficient Methods for Phylogenetic Inference in the Genomic Era. *Mol. Biol. Evol.*, *37*(5), 1530-1534. <https://doi.org/10.1093/molbev/msaa015>

Zhang, C., & Mirarab, S. (2022). ASTRAL-Pro 2: ultrafast species tree reconstruction from multi-copy gene family trees. *Bioinformatics*, *38*(21), 4949-4950. <https://doi.org/10.1093/bioinformatics/btac620>
