## Supplementary material for "Universal orthologs infer deep phylogenies and improve genome quality assessments": All figures and tables: SFig09.pdf

| | Case | Shared connections (I) | Unique connections (U) | Δcontigs | $\frac{I}{U - \Delta contigs}$ % | Diagram |
|---|---|---|---|---|---|---|
| a. | Lower query contiguity | 24 | 36 | 15-3 | 100.00% | 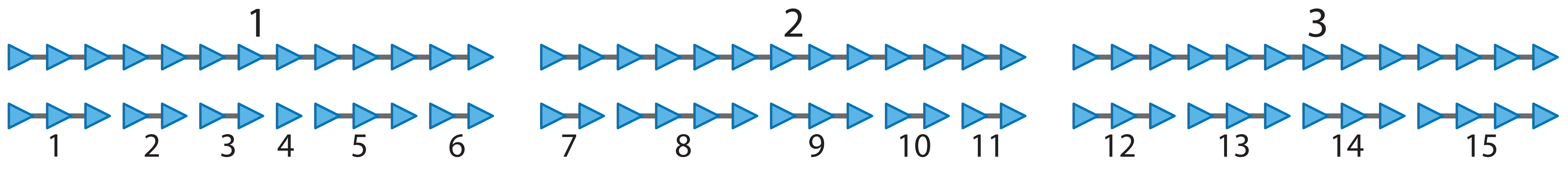 |
| b. | Insertion/ Deletion | 36-2 | 36+1 | 3-3 | 91.89% | 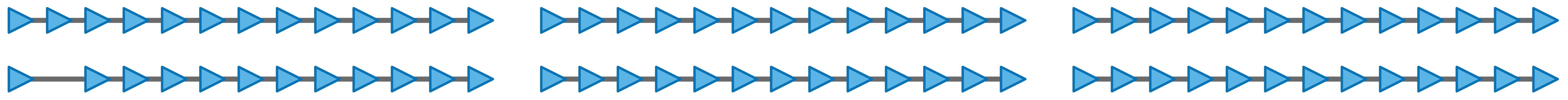 |
| c. | Terminal Insertion/ Deletion | 36-1 | 36 | 3-3 | 97.22% | 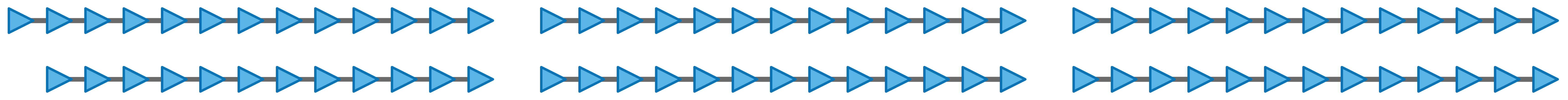 |
| d. | Inversion | 36-2 | 36+2 | 3-3 | 89.47% | 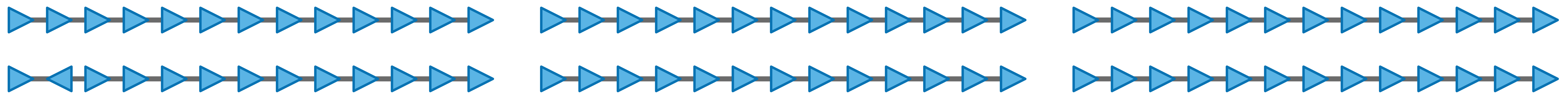 |
| e. | Terminal Inversion | 36-1 | 36+1 | 3-3 | 94.59% | 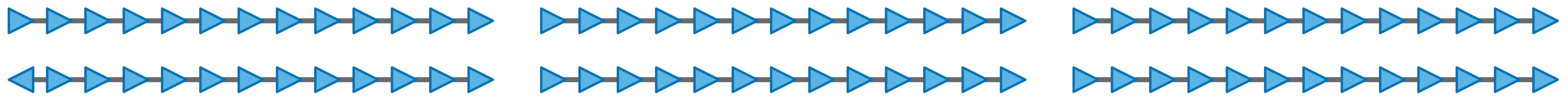 |
| f. | Fragmented reference and query | 15 | 35 | 17-11 | 51.72% | 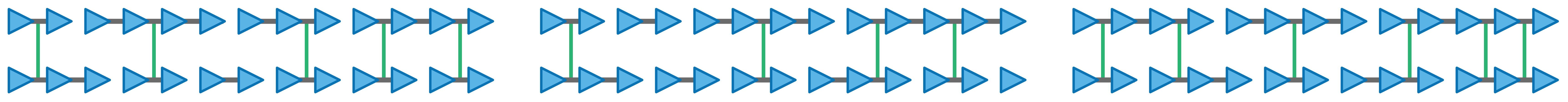 |
