## Supplementary material for "Universal orthologs infer deep phylogenies and improve genome quality assessments": All figures and tables: SFig10.pdf

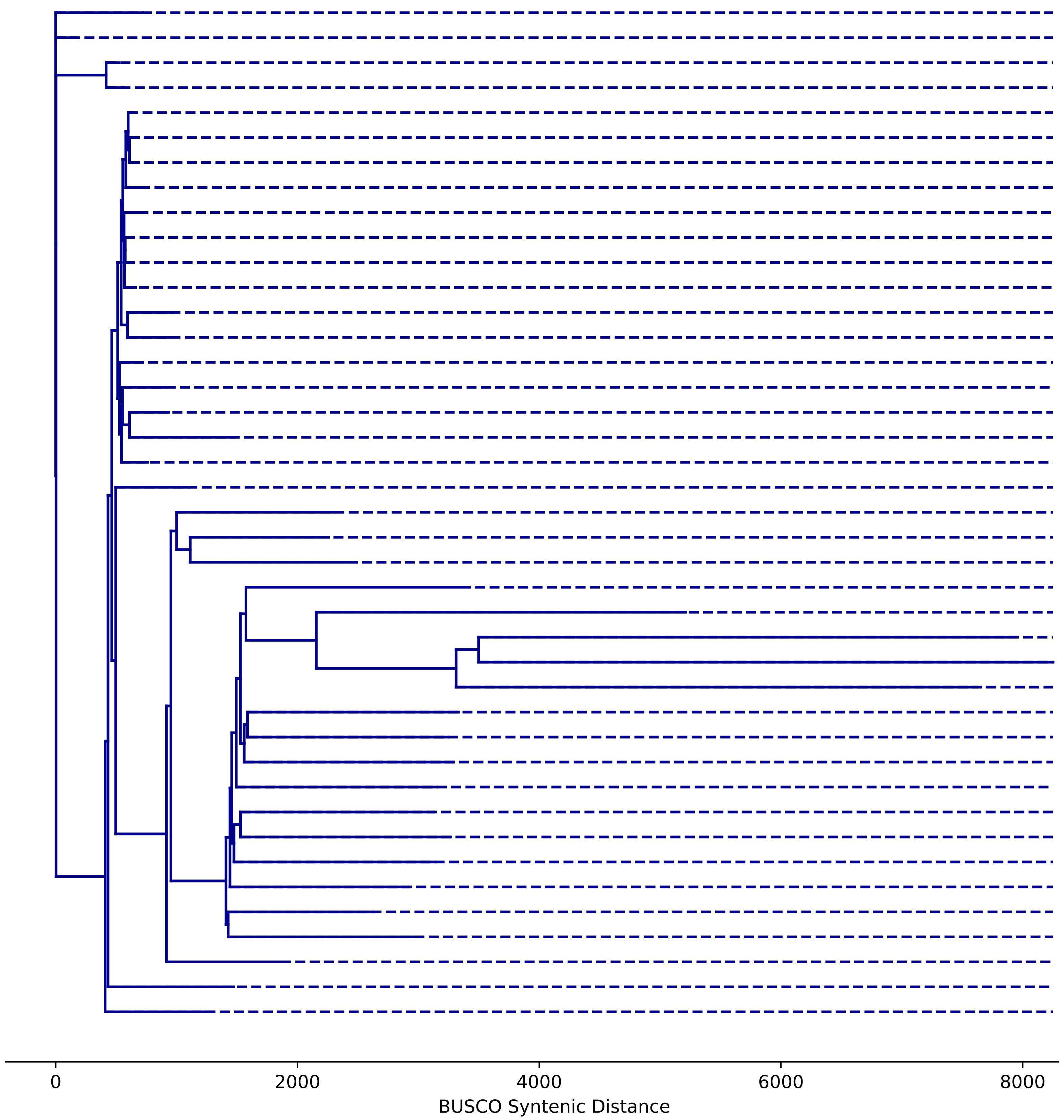


| | Size(Mb) | N50(Kb) | N70(Kb) | N90(Kb) | Contigs | L50 | L70 | L90 | BUSCO | CUSCO | MUSCO |
|---|---|---|---|---|---|---|---|---|---|---|---|
| GCA_014636015.1 | 404.81 | 17256.39 | 12462.11 | 4498.34 | 389 | 8 | 14 | 23 | 99.75 | 99.68 | 100.00 |
| GCA_019137765.1 | 407.50 | 33099.52 | 31036.91 | 25302.85 | 89 | 6 | 8 | 11 | 99.75 | 99.72 | 99.86 |
| GCA_009831255.1 | 380.35 | 30566.71 | 29509.71 | 24626.21 | 41 | 6 | 8 | 11 | 99.72 | 99.68 | 99.86 |
| GCA_009831315.1 | 383.24 | 30350.17 | 28801.51 | 23736.29 | 81 | 6 | 8 | 11 | 99.72 | 99.64 | 100.00 |
| GCA_964197605.1 | 379.81 | 30319.36 | 29419.42 | 23779.28 | 90 | 6 | 8 | 11 | 99.72 | 99.64 | 100.00 |
| GCA_964197625.1 | 377.36 | 30259.89 | 28931.15 | 24061.86 | 282 | 6 | 8 | 11 | 99.72 | 99.64 | 100.00 |
| GCA_964197595.1 | 373.56 | 30201.57 | 28766.06 | 23206.78 | 770 | 6 | 8 | 11 | 99.66 | 99.61 | 99.86 |
| GCA_964197615.1 | 375.61 | 29979.53 | 28855.22 | 23401.26 | 311 | 6 | 8 | 11 | 99.60 | 99.49 | 100.00 |
| GCA_009797565.1 | 381.57 | 30273.40 | 29683.24 | 24066.87 | 33 | 6 | 8 | 11 | 99.72 | 99.64 | 100.00 |
| GCA_022834795.1 | 375.08 | 30085.29 | 29371.99 | 23720.86 | 12 | 6 | 8 | 11 | 99.69 | 99.61 | 100.00 |
| GCA_003865235.1 | 379.63 | 29958.02 | 28772.90 | 23519.17 | 115 | 6 | 8 | 11 | 99.69 | 99.61 | 100.00 |
| GCA_001433935.1 | 373.80 | 29958.43 | 29021.11 | 23207.29 | 55 | 6 | 8 | 11 | 99.60 | 99.49 | 100.00 |
| GCA_000817615.1 | 342.03 | 26709.88 | 24994.91 | 20313.61 | 1160 | 6 | 8 | 11 | 99.10 | 99.21 | 98.72 |
| GCA_000817635.1 | 337.74 | 25188.48 | 23957.62 | 19575.78 | 1739 | 6 | 9 | 12 | 99.51 | 99.45 | 99.71 |
| GCA_026167685.1 | 378.78 | 30175.50 | 29586.00 | 24418.40 | 12 | 6 | 8 | 11 | 99.72 | 99.64 | 100.00 |
| GCA_006942195.1 | 377.67 | 29566.61 | 28175.35 | 22207.90 | 350 | 6 | 8 | 11 | 99.72 | 99.64 | 100.00 |
| GCA_003449065.1 | 378.10 | 7159.81 | 4338.98 | 2530.54 | 144 | 17 | 30 | 53 | 99.72 | 99.64 | 100.00 |
| GCA_002573525.1 | 418.90 | 2522.75 | 1542.84 | 498.85 | 588 | 52 | 95 | 179 | 99.72 | 99.64 | 100.00 |
| GCA_034317485.1 | 392.28 | 27391.61 | 19226.22 | 10435.02 | 304 | 7 | 11 | 16 | 99.72 | 99.64 | 100.00 |
| GCA_004007595.1 | 377.60 | 29563.13 | 27359.35 | 25885.63 | 615 | 6 | 8 | 11 | 99.54 | 99.53 | 99.57 |
| GCA_019338905.1 | 378.29 | 29236.86 | 27064.68 | 22434.35 | 4480 | 6 | 8 | 11 | 97.03 | 96.69 | 98.29 |
| GCA_000149285.1 | 391.15 | 29309.11 | 23730.40 | 21480.89 | 7670 | 6 | 9 | 12 | 98.15 | 98.15 | 98.15 |
| GCA_000004655.2 | 426.34 | 31162.56 | 22971.67 | 16.95 | 10490 | 6 | 9 | 357 | 97.68 | 97.44 | 98.57 |
| GCA_028584095.1 | 388.08 | 1930.79 | 965.42 | 261.95 | 1132 | 54 | 110 | 256 | 97.16 | 96.77 | 98.57 |
| GCA_000725085.2 | 389.75 | 22509.09 | 15437.93 | 23.01 | 15907 | 7 | 12 | 1177 | 97.00 | 96.73 | 98.00 |
| GCA_001611195.1 | 351.23 | 22.01 | 12.75 | 3.95 | 39669 | 4600 | 8787 | 18033 | 96.14 | 95.58 | 98.15 |
| GCA_001611255.1 | 331.82 | 16.51 | 9.66 | 2.64 | 75421 | 5890 | 11102 | 23068 | 93.91 | 93.57 | 95.15 |
| GCA_001611235.1 | 352.23 | 24.43 | 11.04 | 1.15 | 116212 | 3762 | 8010 | 26744 | 94.31 | 93.96 | 95.58 |
| GCA_936157125.1 | 419.73 | 3205.73 | 1203.48 | 9.90 | 12563 | 39 | 82 | 2126 | 98.39 | 98.19 | 99.14 |
| GCA_936154975.1 | 419.19 | 2026.04 | 804.74 | 9.48 | 12856 | 51 | 115 | 2167 | 98.58 | 98.50 | 98.86 |
| GCA_936157035.1 | 427.93 | 3896.25 | 1489.00 | 10.01 | 12764 | 29 | 62 | 2167 | 98.39 | 98.26 | 98.86 |
| GCA_028584105.1 | 450.00 | 1989.71 | 987.99 | 115.75 | 1176 | 56 | 119 | 437 | 99.69 | 99.61 | 100.00 |
| GCA_936157175.1 | 412.46 | 3715.76 | 1858.70 | 10.80 | 11525 | 32 | 61 | 1669 | 98.33 | 98.26 | 98.57 |
| GCA_936153055.1 | 424.15 | 4571.06 | 2354.24 | 11.38 | 11576 | 22 | 47 | 1787 | 98.55 | 98.42 | 99.00 |
| GCA_936145055.1 | 427.37 | 3690.46 | 1585.53 | 10.81 | 12303 | 30 | 63 | 2061 | 98.83 | 98.82 | 98.86 |
| GCA_936157615.1 | 419.10 | 3389.81 | 1379.68 | 10.87 | 11832 | 34 | 69 | 1793 | 99.01 | 98.90 | 99.43 |
| GCA_936158435.1 | 411.00 | 2314.66 | 1103.94 | 11.92 | 10313 | 43 | 92 | 1133 | 99.23 | 99.17 | 99.43 |
| GCA_936150925.1 | 427.61 | 4158.35 | 2004.24 | 10.49 | 12202 | 28 | 55 | 1941 | 98.55 | 98.38 | 99.14 |
| GCA_003449045.1 | 388.77 | 3054.50 | 1756.26 | 648.95 | 410 | 40 | 75 | 145 | 99.63 | 99.57 | 99.86 |
| GCA_011764405.2 | 366.59 | 25799.89 | 24329.76 | 20221.20 | 1402 | 6 | 9 | 12 | 97.31 | 97.12 | 98.00 |
| GCA_004348155.2 | 415.39 | 28085.72 | 23793.43 | 1047.82 | 367 | 7 | 10 | 29 | 99.72 | 99.64 | 100.00 |
