## Supplementary material for "Universal orthologs infer deep phylogenies and improve genome quality assessments": All figures and tables: SFig11.pdf

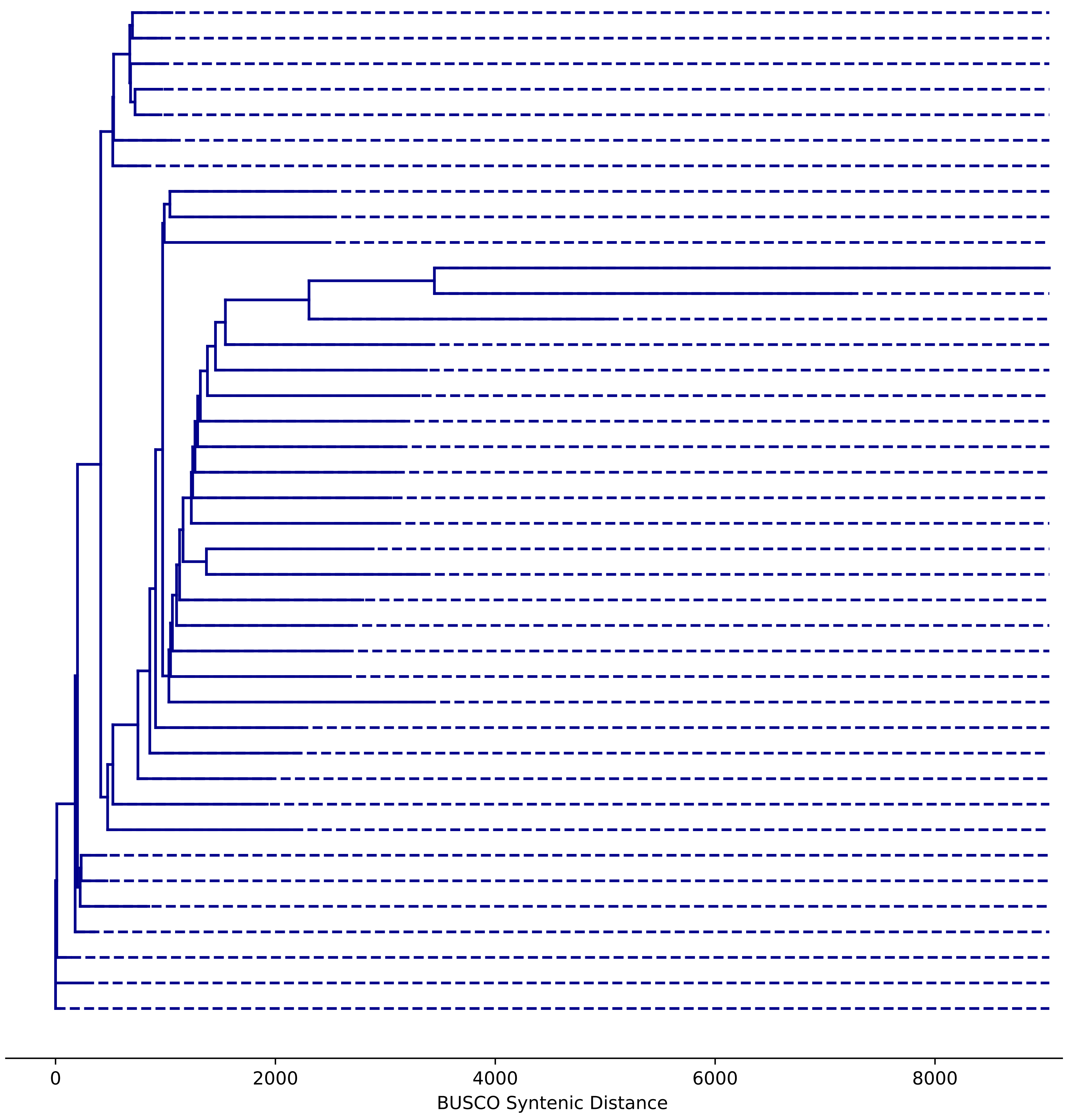


| | Size(Mb) | N50(Kb) | N70(Kb) | N90(Kb) | Contigs | L50 | L70 | L90 | BUSCO | CUSCO | MUSCO |
|---|---|---|---|---|---|---|---|---|---|---|---|
| GCA_001624295.1 | 2712.71 | 132982.30 | 122086.46 | 96871.11 | 5953 | 9 | 13 | 18 | 99.70 | 99.66 | 99.86 |
| GCA_001624505.1 | 2606.16 | 128213.66 | 117921.36 | 93736.62 | 4105 | 9 | 13 | 18 | 99.67 | 99.62 | 99.86 |
| GCA_001624535.1 | 2588.62 | 127063.09 | 116519.28 | 93135.23 | 5013 | 9 | 13 | 18 | 99.61 | 99.55 | 99.86 |
| GCA_001624215.1 | 2630.00 | 129072.80 | 118520.68 | 94837.08 | 4688 | 9 | 13 | 18 | 99.82 | 99.77 | 100.00 |
| GCA_001632525.1 | 2627.34 | 129642.64 | 118936.92 | 95163.05 | 3825 | 9 | 13 | 18 | 99.67 | 99.66 | 99.71 |
| GCA_001624775.1 | 2559.99 | 128387.51 | 116378.84 | 93366.00 | 3140 | 9 | 13 | 18 | 99.70 | 99.70 | 99.71 |
| GCA_001624745.1 | 2699.27 | 132960.83 | 121280.32 | 97461.82 | 7022 | 9 | 13 | 18 | 99.85 | 99.81 | 100.00 |
| GCA_029237445.1 | 2528.20 | 7136.54 | 4321.21 | 1907.33 | 1908 | 99 | 188 | 360 | 99.73 | 99.70 | 99.86 |
| GCA_029237485.1 | 2526.70 | 7413.31 | 4678.41 | 1873.97 | 1731 | 98 | 185 | 344 | 99.79 | 99.74 | 100.00 |
| GCA_029237465.1 | 2526.73 | 7964.06 | 4629.58 | 2058.09 | 1694 | 93 | 177 | 336 | 99.73 | 99.66 | 100.00 |
| GCA_000181195.1 | 2477.64 | 24.82 | 15.07 | 5.29 | 224713 | 29731 | 55277 | 106996 | 63.51 | 60.36 | 75.43 |
| GCA_000185125.1 | 2561.49 | 357.13 | 183.78 | 50.78 | 41268 | 1917 | 3912 | 8936 | 94.87 | 94.35 | 96.86 |
| GCA_003774525.2 | 2801.30 | 1291.02 | 477.75 | 80.84 | 12661 | 585 | 1241 | 4960 | 98.15 | 97.81 | 99.43 |
| GCA_029234035.1 | 2512.85 | 3745.91 | 2210.93 | 879.50 | 3725 | 197 | 370 | 717 | 99.55 | 99.47 | 99.86 |
| GCA_029234005.1 | 2517.20 | 3603.11 | 1976.62 | 816.16 | 3115 | 200 | 390 | 766 | 99.58 | 99.47 | 100.00 |
| GCA_029234525.1 | 2515.83 | 3919.29 | 2267.37 | 878.12 | 3917 | 186 | 355 | 696 | 99.67 | 99.62 | 99.86 |
| GCA_029237225.1 | 2516.74 | 4571.60 | 2649.62 | 1088.69 | 3246 | 158 | 300 | 584 | 99.76 | 99.70 | 100.00 |
| GCA_029237405.1 | 2516.14 | 3785.44 | 2306.60 | 887.82 | 3331 | 183 | 355 | 694 | 99.76 | 99.70 | 100.00 |
| GCA_029237415.1 | 2535.04 | 3972.16 | 2382.98 | 922.47 | 4400 | 184 | 351 | 681 | 99.61 | 99.51 | 100.00 |
| GCA_029236065.1 | 2521.90 | 4797.54 | 2800.14 | 1069.50 | 3589 | 147 | 281 | 562 | 99.76 | 99.74 | 99.86 |
| GCA_029237205.1 | 2516.91 | 5108.57 | 3116.53 | 1137.04 | 3470 | 141 | 266 | 525 | 99.79 | 99.74 | 100.00 |
| GCA_029233695.1 | 2509.21 | 6177.28 | 3698.84 | 1589.27 | 2502 | 110 | 215 | 412 | 99.76 | 99.74 | 99.86 |
| GCA_029233825.1 | 2512.91 | 3464.13 | 1961.98 | 765.91 | 3264 | 200 | 391 | 787 | 99.64 | 99.55 | 100.00 |
| GCA_029237185.1 | 2522.56 | 5930.76 | 3609.72 | 1378.80 | 2781 | 125 | 232 | 448 | 99.64 | 99.55 | 100.00 |
| GCA_029255665.1 | 2524.22 | 6321.87 | 4064.59 | 1602.17 | 2320 | 124 | 223 | 415 | 99.73 | 99.66 | 100.00 |
| GCA_029234015.1 | 2504.13 | 7420.52 | 4421.25 | 1850.41 | 1922 | 100 | 188 | 357 | 99.31 | 99.17 | 99.86 |
| GCA_029255695.1 | 2526.58 | 7006.83 | 4221.87 | 1847.78 | 2402 | 104 | 197 | 373 | 99.73 | 99.66 | 100.00 |
| GCA_000350445.1 | 2578.29 | 6876.43 | 3980.32 | 1298.36 | 9129 | 107 | 204 | 422 | 97.47 | 97.51 | 97.29 |
| GCA_029233705.1 | 2520.72 | 10391.91 | 7048.28 | 2711.36 | 1910 | 74 | 131 | 242 | 99.73 | 99.70 | 99.86 |
| GCA_029233295.1 | 2524.56 | 10812.48 | 7203.99 | 2791.82 | 2019 | 75 | 132 | 241 | 99.79 | 99.77 | 99.86 |
| GCA_000185105.2 | 2505.02 | 18924.32 | 11751.14 | 5711.73 | 2314 | 40 | 74 | 133 | 99.49 | 99.51 | 99.43 |
| GCA_029237265.1 | 2511.71 | 15401.79 | 10619.80 | 4450.29 | 1629 | 48 | 86 | 159 | 99.82 | 99.85 | 99.71 |
| GCA_026508895.1 | 2570.42 | 62404.61 | 29190.75 | 9839.57 | 14365 | 15 | 27 | 56 | 97.88 | 97.93 | 97.71 |
| GCA_947599735.1 | 2856.33 | 134505.79 | 121687.77 | 90037.83 | 428 | 9 | 14 | 19 | 99.88 | 99.85 | 100.00 |
| GCA_921998315.2 | 2603.17 | 130880.72 | 119204.92 | 93577.35 | 352 | 9 | 13 | 18 | 99.85 | 99.81 | 100.00 |
| GCA_031761455.1 | 2754.16 | 101020.26 | 88005.35 | 30107.43 | 1089 | 11 | 16 | 26 | 99.85 | 99.81 | 100.00 |
| GCA_947593165.1 | 2822.92 | 137857.04 | 122237.72 | 89861.32 | 383 | 9 | 13 | 19 | 99.88 | 99.85 | 100.00 |
| GCA_949316315.1 | 2770.97 | 132150.48 | 126876.46 | 96079.41 | 179 | 9 | 13 | 18 | 99.76 | 99.74 | 99.86 |
| GCA_964188535.1 | 2731.31 | 137126.21 | 128273.80 | 105498.49 | 237 | 9 | 13 | 17 | 98.60 | 98.38 | 99.43 |
| GCA_000001635.9 | 2728.22 | 130530.86 | 121973.37 | 95294.70 | 61 | 9 | 13 | 18 | 99.88 | 99.85 | 100.00 |
