## Supplementary material for "Universal orthologs infer deep phylogenies and improve genome quality assessments": All figures and tables: SFig12.pdf

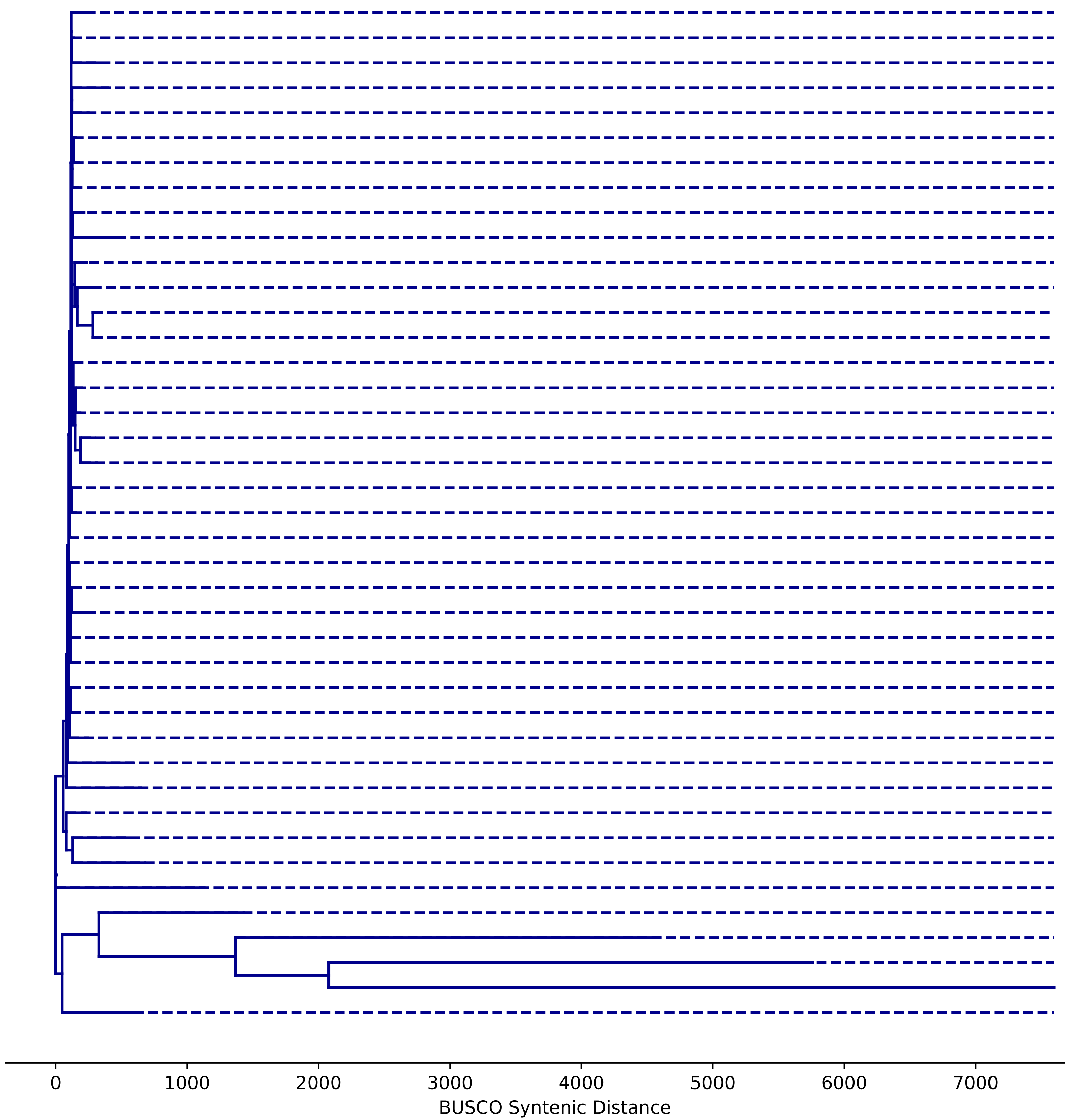


| | Size(Mb) | N50(Kb) | N70(Kb) | N90(Kb) | Contigs | L50 | L70 | L90 | BUSCO | CUSCO | MUSCO |
|---|---|---|---|---|---|---|---|---|---|---|---|
| GCA_020141875.1 | 138.15 | 25557.19 | 24583.27 | 23517.12 | 14 | 3 | 4 | 5 | 99.80 | 99.78 | 100.00 |
| GCA_029775095.1 | 133.88 | 25286.94 | 23542.27 | 23513.71 | 6 | 3 | 4 | 5 | 99.90 | 99.89 | 100.00 |
| GCA_003401925.1 | 135.98 | 24831.14 | 23702.82 | 23375.02 | 39 | 3 | 4 | 5 | 99.80 | 99.78 | 100.00 |
| GCA_020141505.1 | 139.17 | 25526.33 | 25495.58 | 23762.15 | 15 | 3 | 4 | 5 | 99.41 | 99.34 | 100.00 |
| GCA_003401685.1 | 137.71 | 25318.29 | 24545.79 | 23560.50 | 54 | 3 | 4 | 5 | 99.80 | 99.78 | 100.00 |
| GCA_003402005.1 | 136.26 | 25296.82 | 23792.57 | 23258.04 | 65 | 3 | 4 | 5 | 99.90 | 99.89 | 100.00 |
| GCA_003401735.1 | 137.67 | 25207.20 | 24111.29 | 23533.79 | 77 | 3 | 4 | 5 | 99.90 | 99.89 | 100.00 |
| GCA_003401795.1 | 133.34 | 24769.04 | 23803.71 | 23344.39 | 44 | 3 | 4 | 5 | 99.70 | 99.67 | 100.00 |
| GCA_000001215.4 | 143.73 | 25286.94 | 23542.27 | 23513.71 | 1870 | 3 | 4 | 5 | 99.90 | 99.89 | 100.00 |
| GCA_034768405.1 | 141.74 | 24695.23 | 23293.80 | 22814.92 | 2104 | 3 | 4 | 5 | 99.70 | 99.67 | 100.00 |
| GCA_038396815.1 | 133.19 | 25190.60 | 23443.58 | 23410.09 | 6 | 3 | 4 | 5 | 99.80 | 99.89 | 99.00 |
| GCA_927717585.1 | 130.59 | 23923.16 | 22771.81 | 22036.39 | 110 | 3 | 4 | 5 | 99.70 | 99.78 | 99.00 |
| GCA_002310755.1 | 120.38 | 23011.49 | 22422.44 | 21146.71 | 6 | 3 | 4 | 5 | 99.31 | 99.34 | 99.00 |
| GCA_002310775.1 | 120.38 | 23011.31 | 22422.82 | 21146.57 | 6 | 3 | 4 | 5 | 99.31 | 99.34 | 99.00 |
| GCA_002300595.1 | 144.13 | 25479.26 | 23785.51 | 23611.14 | 160 | 3 | 4 | 5 | 99.80 | 99.78 | 100.00 |
| GCA_004798055.1 | 133.40 | 24495.47 | 24035.72 | 22975.41 | 89 | 3 | 4 | 5 | 99.90 | 99.89 | 100.00 |
| GCA_003401885.1 | 133.23 | 24418.02 | 23538.80 | 23314.57 | 29 | 3 | 4 | 5 | 99.90 | 99.89 | 100.00 |
| GCA_927718685.1 | 131.47 | 24271.30 | 22246.14 | 21964.12 | 101 | 3 | 4 | 5 | 99.80 | 99.89 | 99.00 |
| GCA_927718715.1 | 131.43 | 23706.51 | 22546.43 | 22157.95 | 115 | 3 | 4 | 5 | 99.80 | 99.89 | 99.00 |
| GCA_003401745.1 | 140.69 | 25025.26 | 23779.42 | 23610.74 | 96 | 3 | 4 | 5 | 99.90 | 99.89 | 100.00 |
| GCA_042606465.1 | 170.24 | 27727.68 | 24085.51 | 368.03 | 258 | 3 | 5 | 21 | 99.90 | 99.89 | 100.00 |
| GCA_042606445.1 | 168.56 | 27447.82 | 24266.33 | 697.09 | 211 | 3 | 5 | 13 | 99.90 | 99.89 | 100.00 |
| GCA_003401975.1 | 137.49 | 25529.27 | 23571.25 | 23463.68 | 58 | 3 | 4 | 5 | 99.80 | 99.78 | 100.00 |
| GCA_037075385.1 | 142.08 | 25438.40 | 23761.45 | 23313.10 | 50 | 3 | 4 | 5 | 99.90 | 99.89 | 100.00 |
| GCA_927720205.1 | 134.57 | 25011.10 | 22999.89 | 22306.85 | 96 | 3 | 4 | 5 | 99.90 | 99.89 | 100.00 |
| GCA_003401855.1 | 138.93 | 25355.51 | 23835.28 | 23721.34 | 99 | 3 | 4 | 5 | 99.90 | 99.89 | 100.00 |
| GCA_042606455.1 | 172.47 | 28052.01 | 23885.99 | 1330.15 | 126 | 3 | 5 | 11 | 99.90 | 99.89 | 100.00 |
| GCA_003397115.2 | 137.46 | 25282.51 | 24131.51 | 23345.28 | 61 | 3 | 4 | 5 | 99.90 | 99.89 | 100.00 |
| GCA_003402055.1 | 136.19 | 24877.00 | 23988.88 | 23380.35 | 43 | 3 | 4 | 5 | 99.90 | 99.89 | 100.00 |
| GCA_003402015.1 | 136.26 | 25138.59 | 23690.58 | 23452.00 | 75 | 3 | 4 | 5 | 99.70 | 99.78 | 99.00 |
| GCA_003401915.1 | 146.86 | 25311.79 | 23796.46 | 23662.37 | 263 | 3 | 4 | 5 | 99.90 | 99.89 | 100.00 |
| GCA_003401805.1 | 142.35 | 24782.13 | 23794.83 | 23548.35 | 173 | 3 | 4 | 5 | 99.80 | 99.78 | 100.00 |
| GCA_018904365.1 | 144.71 | 22510.35 | 21632.78 | 966.60 | 290 | 3 | 5 | 14 | 99.90 | 99.89 | 100.00 |
| GCA_039694015.1 | 163.97 | 6138.01 | 2266.91 | 492.54 | 477 | 8 | 16 | 49 | 99.90 | 99.89 | 100.00 |
| GCA_039694025.1 | 162.69 | 5466.20 | 2032.99 | 413.12 | 468 | 11 | 21 | 56 | 99.90 | 99.89 | 100.00 |
| GCA_015852585.1 | 167.83 | 30895.89 | 28749.85 | 26146.15 | 16 | 3 | 4 | 5 | 99.90 | 99.89 | 100.00 |
| GCA_002050065.1 | 138.49 | 15305.62 | 1922.40 | 338.37 | 128 | 4 | 9 | 50 | 92.69 | 92.66 | 93.00 |
| GCA_000705575.1 | 153.28 | 109.25 | 41.17 | 9.70 | 5066 | 338 | 792 | 2450 | 99.80 | 99.78 | 100.00 |
| GCA_001014345.1 | 117.81 | 47.26 | 16.87 | 4.31 | 9895 | 518 | 1380 | 4163 | 96.64 | 96.50 | 98.00 |
| GCA_009869825.1 | 118.12 | 6.09 | 3.78 | 1.93 | 26805 | 4952 | 9888 | 18512 | 92.10 | 92.11 | 92.00 |
| GCA_000778455.1 | 164.08 | 13636.57 | 830.16 | 53.45 | 789 | 4 | 22 | 188 | 99.11 | 99.01 | 100.00 |
