## Supplementary material for "Universal orthologs infer deep phylogenies and improve genome quality assessments": All figures and tables: SFig13.pdf

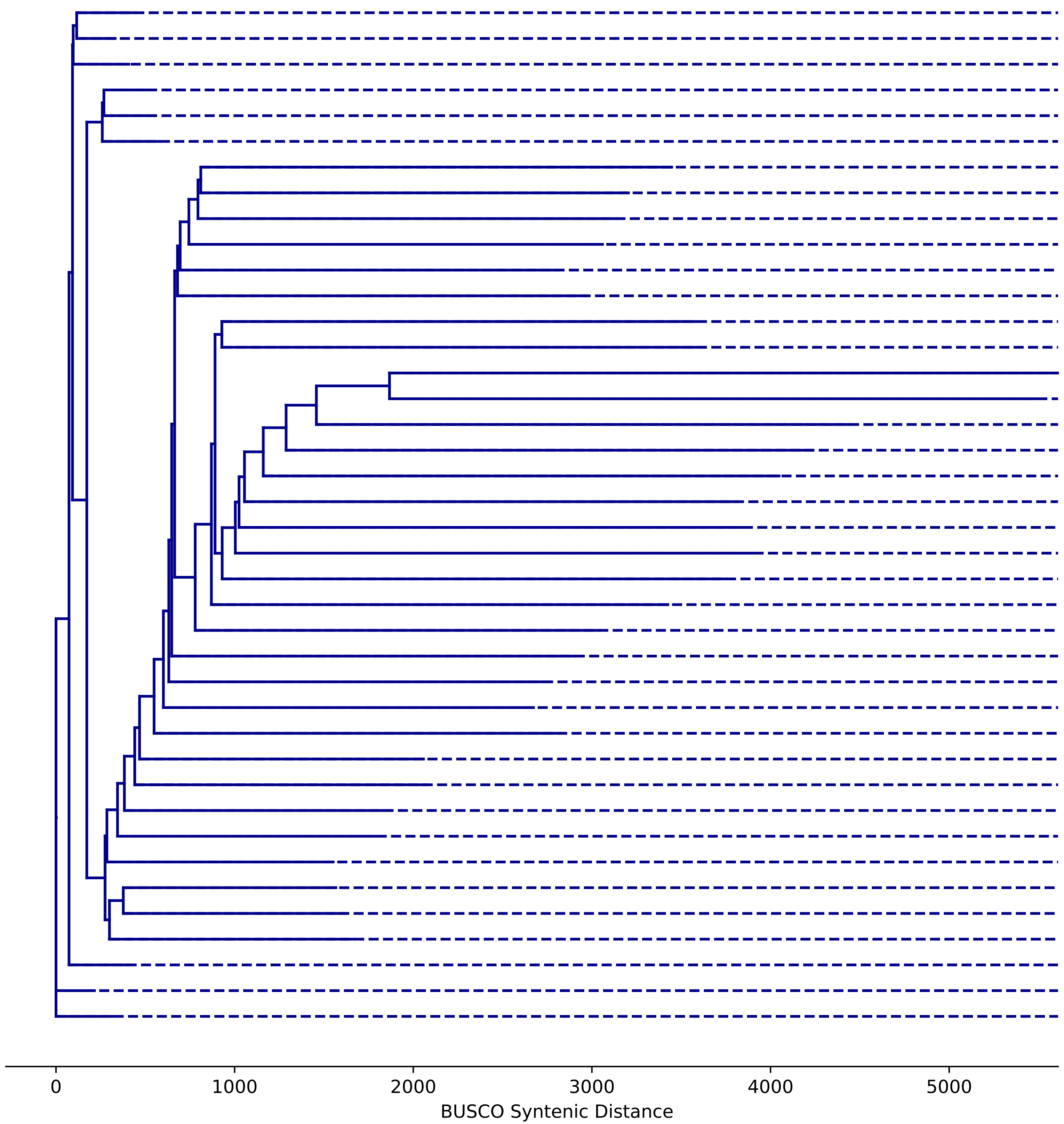


| | Size(Mb) | N50(Kb) | N70(Kb) | N90(Kb) | Contigs | L50 | L70 | L90 | BUSCO | CUSCO | MUSCO |
|---|---|---|---|---|---|---|---|---|---|---|---|
| GCA_018804185.1 | 2901.67 | 96203.34 | 73688.84 | 45761.87 | 937 | 9 | 16 | 26 | 99.97 | 99.96 | 100.00 |
| GCA_022416915.1 | 2906.63 | 96526.83 | 72555.56 | 47042.45 | 1242 | 9 | 16 | 26 | 99.97 | 99.96 | 100.00 |
| GCA_022432845.1 | 2867.73 | 96196.36 | 72325.19 | 45408.78 | 1765 | 9 | 16 | 26 | 99.97 | 99.96 | 100.00 |
| GCA_022538005.1 | 2861.63 | 95262.85 | 72647.34 | 44919.32 | 1455 | 9 | 16 | 26 | 99.97 | 99.96 | 100.00 |
| GCA_024222175.1 | 2955.25 | 94576.23 | 73297.51 | 44153.90 | 1056 | 10 | 17 | 27 | 99.97 | 99.96 | 100.00 |
| GCA_022416745.1 | 2859.21 | 95064.12 | 72608.55 | 45583.69 | 1324 | 9 | 16 | 26 | 99.94 | 99.96 | 99.86 |
| GCA_021327145.1 | 2807.60 | 6299.41 | 2513.82 | 368.57 | 4448 | 91 | 233 | 820 | 96.66 | 96.23 | 98.29 |
| GCA_021328945.1 | 2845.24 | 10647.78 | 4182.18 | 324.38 | 4646 | 59 | 144 | 624 | 97.50 | 97.14 | 98.86 |
| GCA_021325935.1 | 2766.26 | 15416.95 | 5670.09 | 416.32 | 4013 | 35 | 97 | 468 | 95.32 | 94.54 | 98.29 |
| GCA_021326735.1 | 2827.71 | 13616.41 | 4743.79 | 452.74 | 4282 | 48 | 118 | 536 | 97.82 | 97.55 | 98.86 |
| GCA_021327315.1 | 2862.91 | 11466.71 | 4521.42 | 432.99 | 3724 | 54 | 129 | 552 | 98.54 | 98.27 | 99.57 |
| GCA_021326705.1 | 2842.46 | 9879.04 | 4290.81 | 404.79 | 3883 | 60 | 146 | 586 | 98.30 | 98.08 | 99.14 |
| GCA_021325815.1 | 2820.07 | 6374.05 | 2530.76 | 385.19 | 3950 | 91 | 229 | 772 | 97.02 | 96.50 | 99.00 |
| GCA_021327245.1 | 2860.44 | 5677.86 | 1806.03 | 330.48 | 4148 | 113 | 295 | 1007 | 97.79 | 97.55 | 98.71 |
| GCA_021327335.1 | 2774.52 | 1262.14 | 573.96 | 161.72 | 7280 | 435 | 1095 | 2848 | 94.34 | 93.52 | 97.43 |
| GCA_021327355.1 | 2822.48 | 1582.88 | 657.85 | 162.72 | 7399 | 356 | 922 | 2625 | 95.35 | 94.80 | 97.43 |
| GCA_021327395.1 | 2750.43 | 4486.36 | 1465.23 | 263.30 | 4962 | 123 | 344 | 1226 | 93.71 | 92.88 | 96.86 |
| GCA_021325875.1 | 2824.96 | 3697.83 | 1317.03 | 267.59 | 5483 | 168 | 422 | 1322 | 97.26 | 97.02 | 98.14 |
| GCA_021325835.1 | 2760.32 | 4519.99 | 1691.10 | 305.22 | 5069 | 118 | 326 | 1072 | 96.15 | 95.63 | 98.14 |
| GCA_021325855.1 | 2803.69 | 5117.83 | 1996.29 | 375.84 | 3929 | 120 | 301 | 937 | 96.60 | 96.08 | 98.57 |
| GCA_021327285.1 | 2804.74 | 4868.96 | 1990.48 | 367.48 | 4205 | 117 | 300 | 958 | 96.81 | 96.42 | 98.29 |
| GCA_021325865.1 | 2831.43 | 4745.64 | 1726.94 | 287.26 | 4824 | 122 | 322 | 1096 | 97.41 | 97.14 | 98.43 |
| GCA_000765115.1 | 2589.83 | 2217.03 | 1252.68 | 435.44 | 6173 | 328 | 641 | 1334 | 99.25 | 99.13 | 99.71 |
| GCA_021326975.1 | 2824.84 | 5713.49 | 2037.62 | 356.35 | 3948 | 104 | 271 | 920 | 97.76 | 97.40 | 99.14 |
| GCA_024256505.1 | 2915.26 | 5192.53 | 2236.75 | 507.04 | 2447 | 128 | 298 | 851 | 98.90 | 98.91 | 98.86 |
| GCA_021325905.1 | 2750.60 | 15207.28 | 6477.38 | 643.76 | 4049 | 37 | 95 | 344 | 97.38 | 96.95 | 99.00 |
| GCA_021328935.1 | 2842.07 | 13934.75 | 5559.03 | 420.27 | 4123 | 44 | 107 | 468 | 98.33 | 98.04 | 99.43 |
| GCA_021327305.1 | 2842.49 | 11708.82 | 4788.56 | 531.13 | 3653 | 49 | 123 | 482 | 98.93 | 98.83 | 99.29 |
| GCA_024256485.1 | 2834.62 | 6587.65 | 2431.66 | 590.99 | 2758 | 89 | 233 | 698 | 98.75 | 98.57 | 99.43 |
| GCA_024222185.1 | 2836.92 | 11828.76 | 6098.90 | 1052.39 | 1871 | 56 | 123 | 344 | 99.40 | 99.32 | 99.71 |
| GCA_021328955.1 | 2846.16 | 13102.69 | 5397.39 | 874.23 | 2654 | 50 | 119 | 375 | 99.14 | 98.94 | 99.86 |
| GCA_024222135.1 | 2915.63 | 16958.06 | 6727.55 | 812.76 | 1548 | 38 | 93 | 353 | 99.34 | 99.21 | 99.86 |
| GCA_021327375.1 | 2887.83 | 18668.66 | 6128.32 | 662.00 | 2726 | 40 | 98 | 374 | 99.52 | 99.51 | 99.57 |
| GCA_021327435.1 | 2858.03 | 32781.37 | 14567.59 | 1267.68 | 2856 | 26 | 51 | 164 | 99.02 | 98.83 | 99.71 |
| GCA_021328995.1 | 2830.31 | 39501.64 | 14992.68 | 1542.54 | 2586 | 18 | 43 | 134 | 97.97 | 97.59 | 99.43 |
| GCA_022702665.1 | 2752.83 | 53161.07 | 32156.60 | 4843.09 | 1953 | 17 | 30 | 70 | 96.66 | 96.01 | 99.14 |
| GCA_022702505.1 | 2772.17 | 62864.30 | 32213.27 | 1634.66 | 1920 | 15 | 27 | 87 | 95.17 | 94.27 | 98.57 |
| GCA_024222265.1 | 2941.69 | 93095.55 | 71848.17 | 45284.25 | 772 | 10 | 17 | 27 | 99.97 | 99.96 | 100.00 |
| GCA_017524585.1 | 2653.84 | 105184.75 | 81071.31 | 52181.98 | 58 | 8 | 14 | 22 | 99.97 | 99.96 | 100.00 |
| GCA_022416685.1 | 2865.42 | 95532.77 | 72280.70 | 45409.24 | 1562 | 9 | 16 | 26 | 99.94 | 99.96 | 99.86 |
