## Supplementary material for "Universal orthologs infer deep phylogenies and improve genome quality assessments": All figures and tables: SFig14.pdf

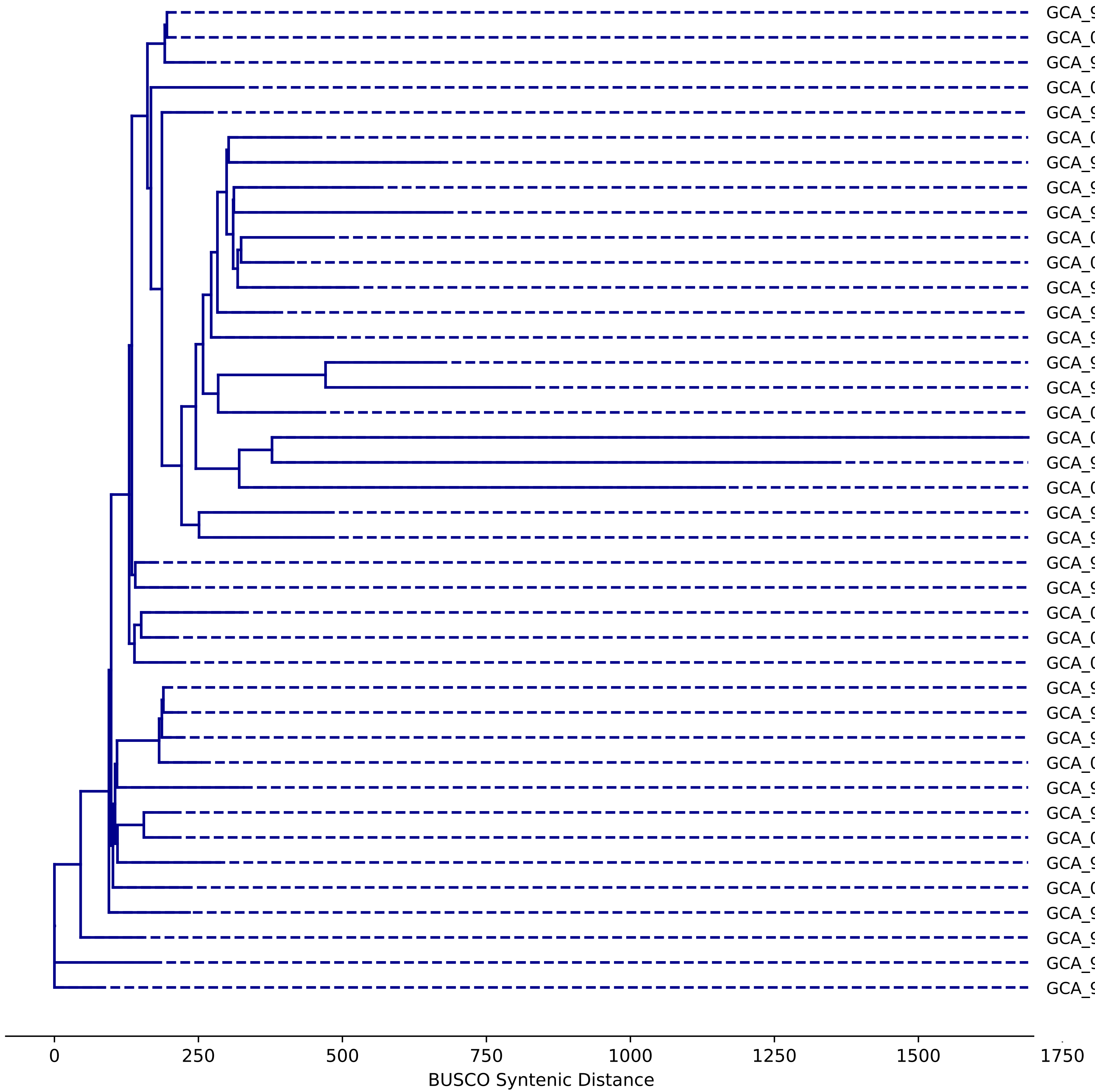


| | Size(Mb) | N50(Kb) | N70(Kb) | N90(Kb) | Contigs | L50 | L70 | L90 | BUSCO | CUSCO | MUSCO |
|---|---|---|---|---|---|---|---|---|---|---|---|
| GCA_902825305.1 | 119.78 | 23424.44 | 20021.70 | 18725.88 | 5 | 3 | 4 | 5 | 99.79 | 99.75 | 100.00 |
| GCA_000001735.2 | 119.67 | 23459.83 | 19698.29 | 18585.06 | 7 | 3 | 4 | 5 | 99.79 | 99.75 | 100.00 |
| GCA_963990585.1 | 119.16 | 23438.90 | 19495.69 | 18473.24 | 93 | 3 | 4 | 5 | 99.74 | 99.70 | 100.00 |
| GCA_025388435.1 | 119.74 | 23021.00 | 19517.76 | 18739.01 | 36 | 3 | 4 | 5 | 99.57 | 99.51 | 100.00 |
| GCA_903064285.1 | 118.89 | 23327.42 | 19659.10 | 19029.45 | 5 | 3 | 4 | 5 | 99.70 | 99.65 | 100.00 |
| GCA_043165235.1 | 128.05 | 11269.41 | 7254.45 | 1559.35 | 152 | 5 | 8 | 14 | 99.74 | 99.70 | 100.00 |
| GCA_900243955.1 | 119.13 | 3101.44 | 2216.73 | 1251.76 | 139 | 13 | 22 | 35 | 99.53 | 99.46 | 100.00 |
| GCA_900243945.1 | 119.17 | 9344.02 | 4022.46 | 1735.65 | 59 | 5 | 10 | 19 | 99.57 | 99.51 | 100.00 |
| GCA_905067165.1 | 124.09 | 14300.87 | 13655.75 | 4269.13 | 35 | 5 | 6 | 9 | 99.79 | 99.75 | 100.00 |
| GCA_043165325.1 | 127.94 | 7619.36 | 4516.24 | 1315.04 | 175 | 7 | 11 | 19 | 99.74 | 99.70 | 100.00 |
| GCA_043165945.1 | 128.79 | 12213.49 | 7497.60 | 1775.32 | 164 | 5 | 7 | 14 | 99.79 | 99.75 | 100.00 |
| GCA_900243935.1 | 119.20 | 7583.94 | 5521.46 | 2551.53 | 40 | 6 | 10 | 17 | 99.70 | 99.70 | 99.67 |
| GCA_927323615.1 | 129.41 | 11244.60 | 8589.54 | 3243.59 | 54 | 5 | 8 | 12 | 99.79 | 99.75 | 100.00 |
| GCA_964264635.1 | 123.96 | 9490.25 | 6735.43 | 1370.85 | 92 | 6 | 8 | 18 | 99.79 | 99.75 | 100.00 |
| GCA_910573875.1 | 117.10 | 12545.48 | 10021.34 | 860.32 | 79 | 5 | 7 | 14 | 99.27 | 99.16 | 100.00 |
| GCA_910574415.1 | 117.09 | 10734.84 | 6144.99 | 636.00 | 101 | 5 | 8 | 17 | 99.27 | 99.16 | 100.00 |
| GCA_000835945.1 | 127.42 | 11163.17 | 8570.85 | 925.68 | 378 | 5 | 8 | 14 | 99.74 | 99.70 | 100.00 |
| GCA_043165315.1 | 128.55 | 787.55 | 491.39 | 175.99 | 390 | 48 | 89 | 171 | 99.23 | 99.11 | 100.00 |
| GCA_964264645.1 | 118.71 | 2301.00 | 1417.62 | 637.47 | 112 | 17 | 30 | 54 | 96.73 | 96.45 | 98.67 |
| GCA_001742845.1 | 116.85 | 589.64 | 246.24 | 40.99 | 5197 | 49 | 108 | 344 | 99.66 | 99.65 | 99.67 |
| GCA_900303345.1 | 119.75 | 10736.98 | 6800.36 | 1093.48 | 78 | 5 | 8 | 18 | 99.44 | 99.36 | 100.00 |
| GCA_900303355.1 | 119.50 | 12334.79 | 7144.16 | 1865.93 | 62 | 5 | 8 | 13 | 99.79 | 99.75 | 100.00 |
| GCA_902460265.3 | 120.13 | 23034.41 | 19417.58 | 18785.46 | 111 | 3 | 4 | 5 | 99.83 | 99.80 | 100.00 |
| GCA_964248535.1 | 126.01 | 23332.55 | 20029.84 | 19833.45 | 38 | 3 | 4 | 5 | 99.79 | 99.75 | 100.00 |
| GCA_024498455.1 | 126.09 | 24097.56 | 20497.02 | 20418.08 | 5 | 3 | 4 | 5 | 99.79 | 99.75 | 100.00 |
| GCA_024498475.1 | 123.95 | 23842.99 | 20908.90 | 20010.34 | 5 | 3 | 4 | 5 | 99.83 | 99.80 | 100.00 |
| GCA_024498555.1 | 124.62 | 24062.51 | 20533.97 | 19986.57 | 5 | 3 | 4 | 5 | 99.79 | 99.75 | 100.00 |
| GCA_902460285.1 | 120.34 | 23366.35 | 19416.17 | 19104.40 | 105 | 3 | 4 | 5 | 99.74 | 99.70 | 100.00 |
| GCA_900660825.1 | 119.63 | 23367.61 | 19252.71 | 19019.44 | 109 | 3 | 4 | 5 | 99.66 | 99.61 | 100.00 |
| GCA_903064275.1 | 117.84 | 22795.61 | 19454.28 | 19127.70 | 5 | 3 | 4 | 5 | 99.70 | 99.65 | 100.00 |
| GCA_001651475.1 | 118.89 | 22588.20 | 19037.55 | 17980.86 | 30 | 3 | 4 | 5 | 99.74 | 99.70 | 100.00 |
| GCA_902460295.1 | 120.29 | 22728.24 | 19510.81 | 18892.29 | 94 | 3 | 4 | 5 | 99.61 | 99.56 | 100.00 |
| GCA_902460315.1 | 120.80 | 23762.78 | 19545.95 | 18579.31 | 142 | 3 | 4 | 5 | 99.74 | 99.70 | 100.00 |
| GCA_024498495.1 | 122.66 | 23758.99 | 20003.80 | 19603.74 | 5 | 3 | 4 | 5 | 99.74 | 99.70 | 100.00 |
| GCA_903064325.1 | 120.69 | 23866.72 | 19794.77 | 19334.17 | 5 | 3 | 4 | 5 | 99.70 | 99.65 | 100.00 |
| GCA_024498515.1 | 120.09 | 22777.72 | 20220.15 | 19437.12 | 5 | 3 | 4 | 5 | 99.57 | 99.51 | 100.00 |
| GCA_902705455.1 | 119.23 | 23835.91 | 19336.25 | 19087.29 | 98 | 3 | 4 | 5 | 99.79 | 99.75 | 100.00 |
| GCA_903064295.1 | 119.80 | 23751.65 | 20049.44 | 19682.55 | 5 | 3 | 4 | 5 | 99.74 | 99.70 | 100.00 |
| GCA_946406895.1 | 148.25 | 30448.87 | 25417.97 | 24321.62 | 5 | 3 | 4 | 5 | 99.79 | 99.75 | 100.00 |
| GCA_902460275.1 | 119.75 | 23195.25 | 20044.95 | 18371.56 | 102 | 3 | 4 | 5 | 99.79 | 99.75 | 100.00 |
