## Supplementary figures and images for "Universal orthologs infer deep phylogenies and improve genome quality assessments"

### Fig01.pdf

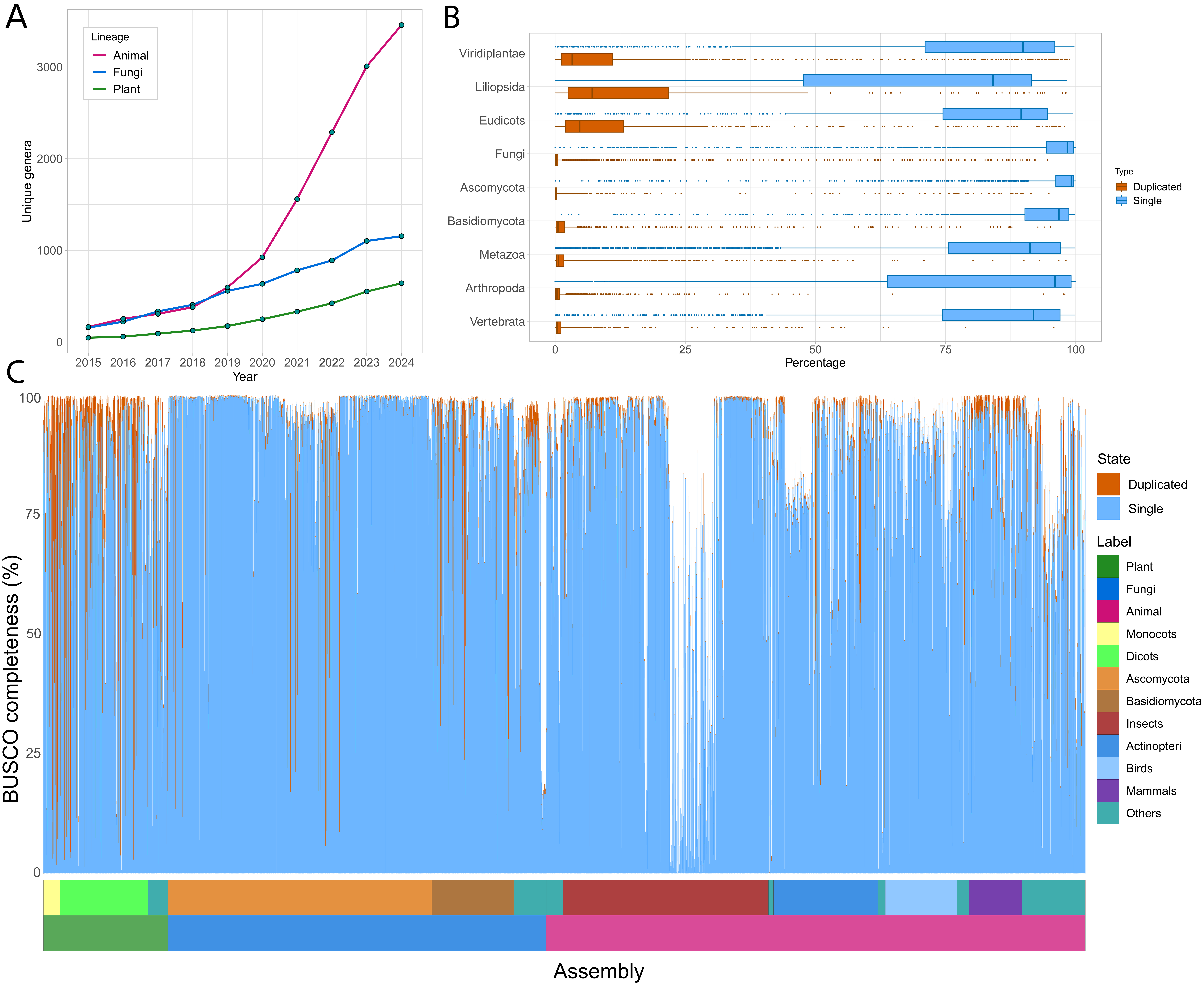

### Fig02.pdf

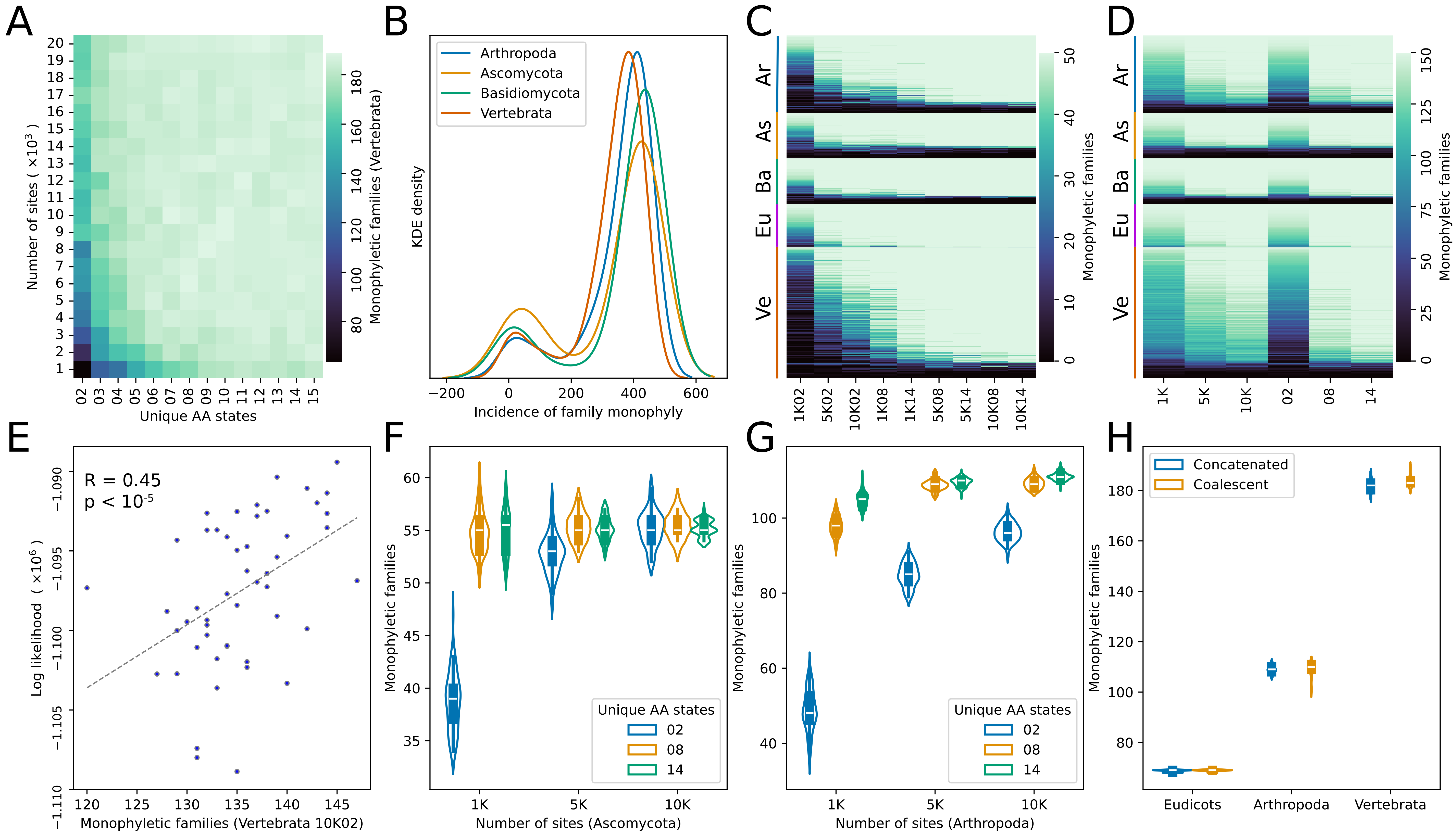

### Fig03.pdf

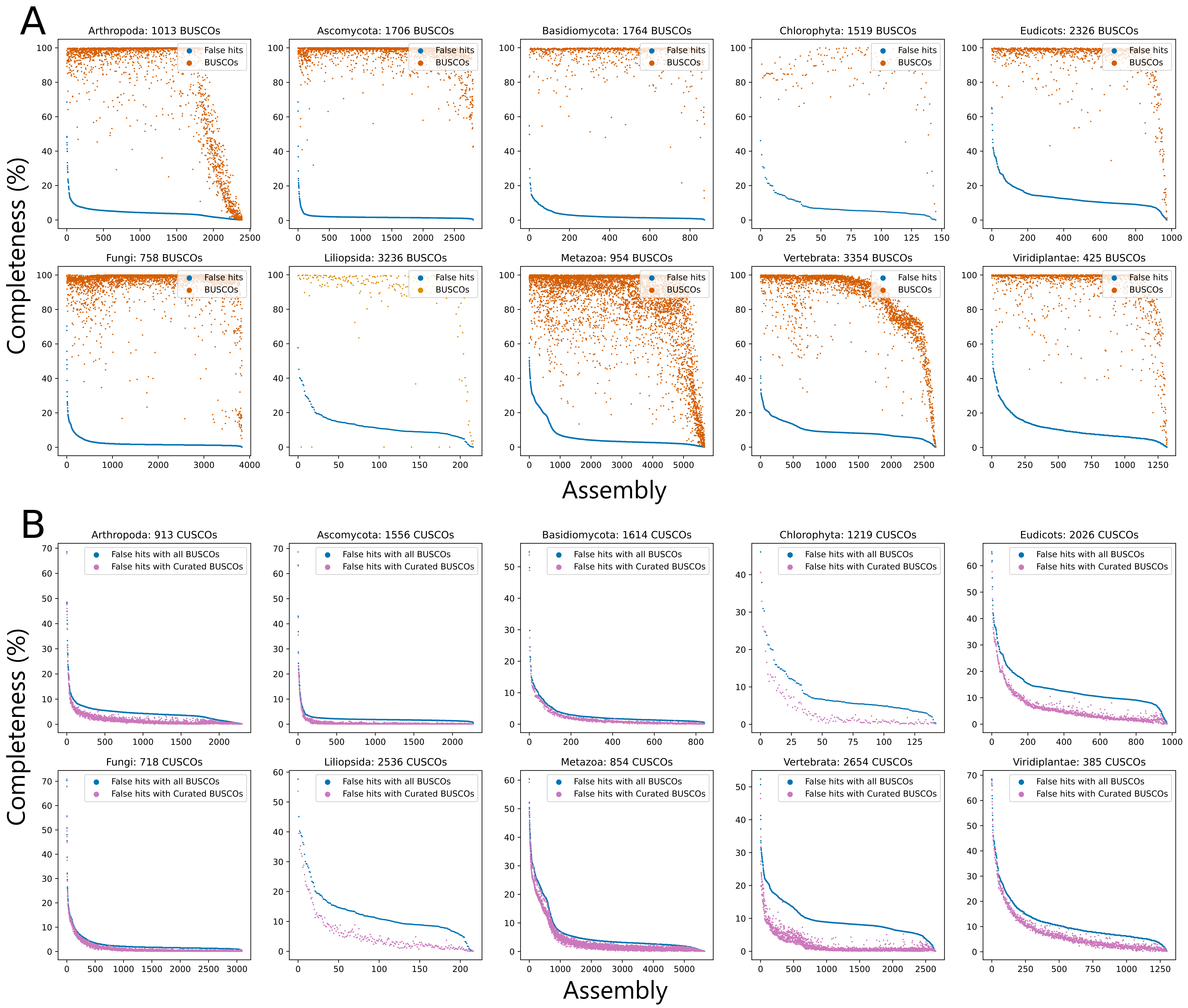

### Fig04.pdf

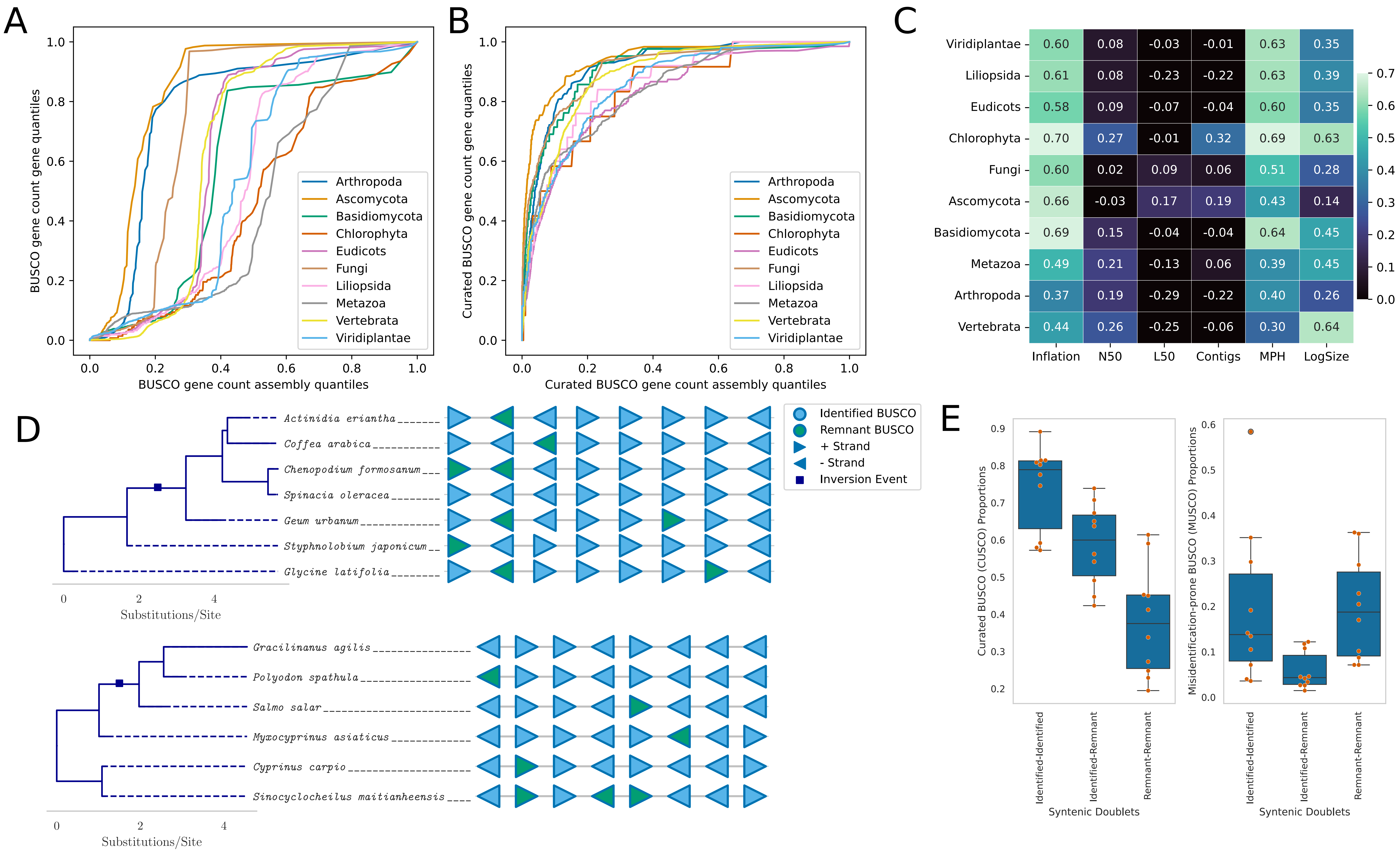

### Fig05.pdf

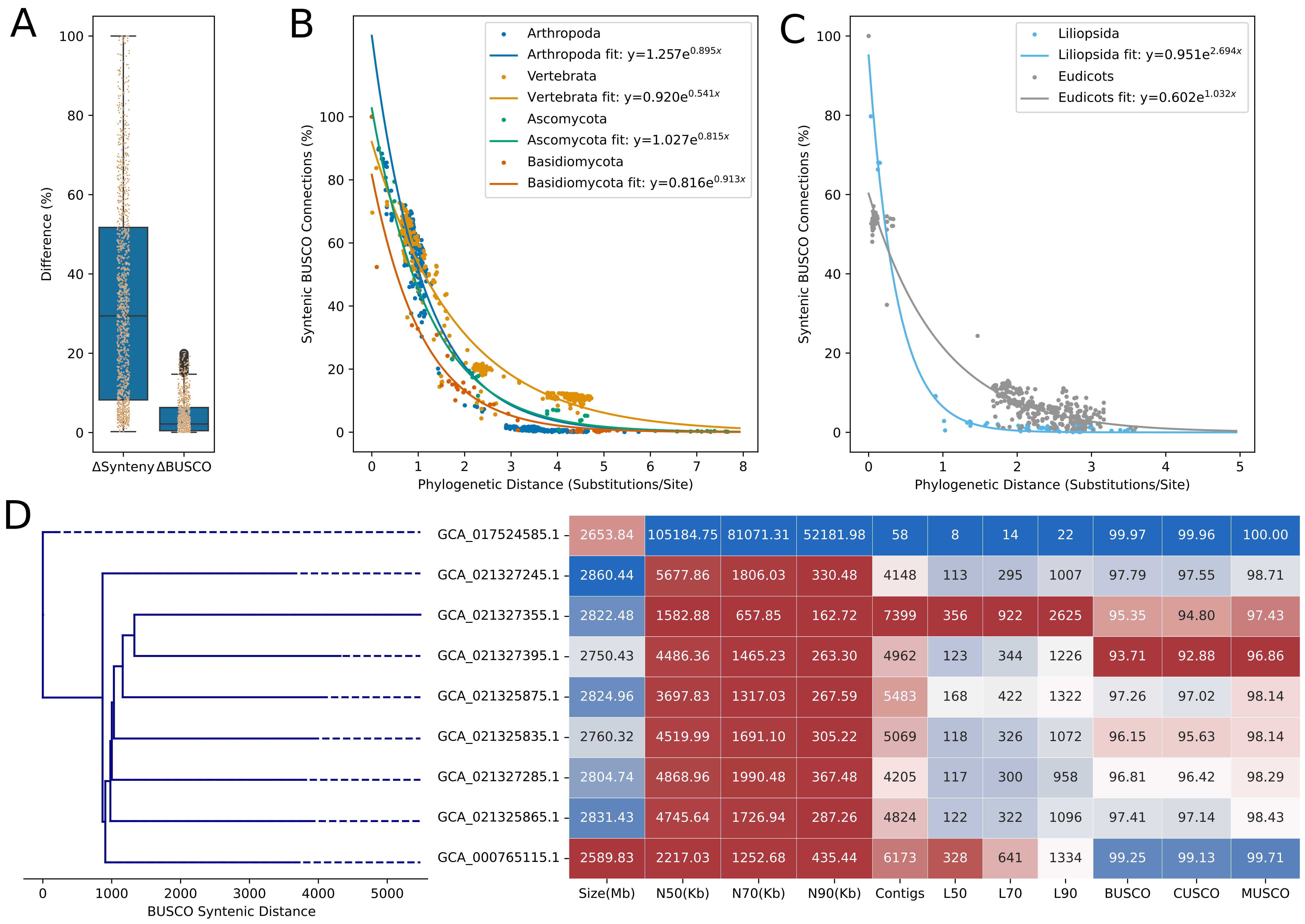

### Fig06.pdf

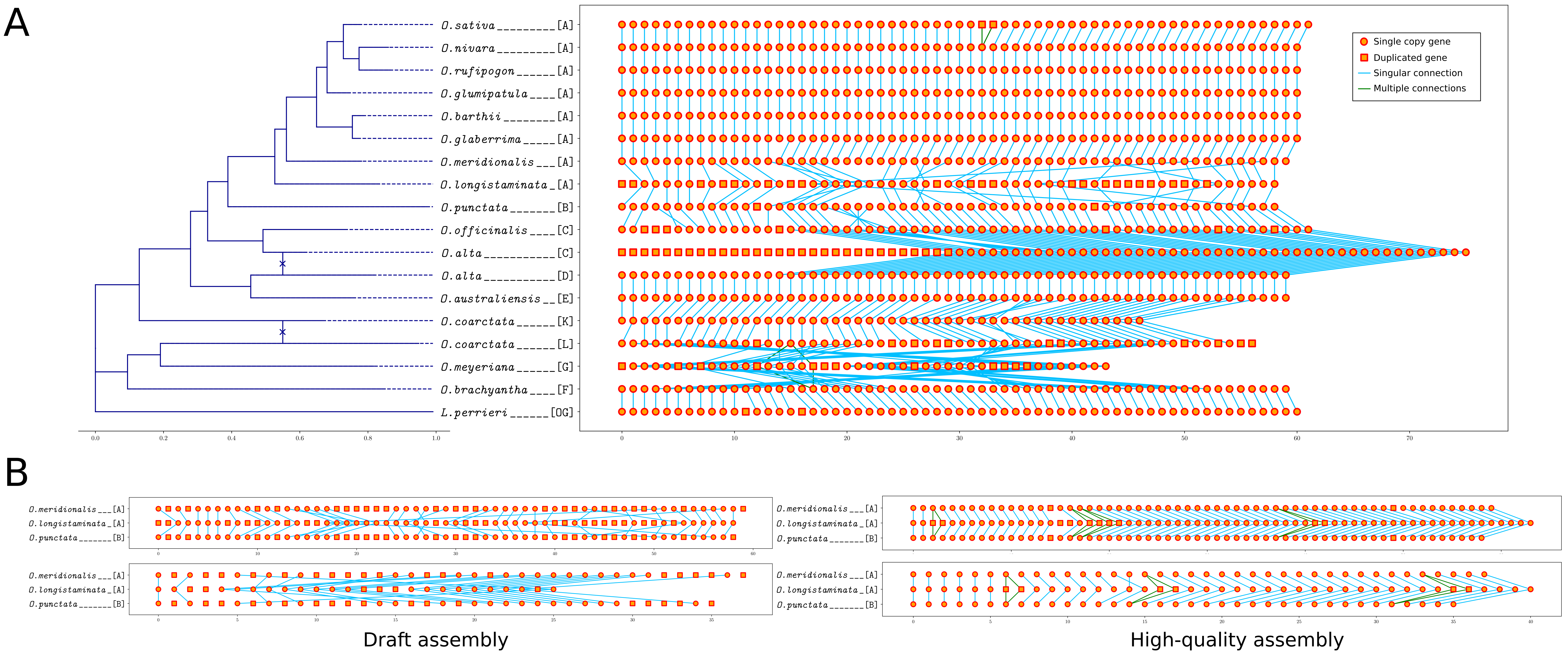

### SFig01.pdf

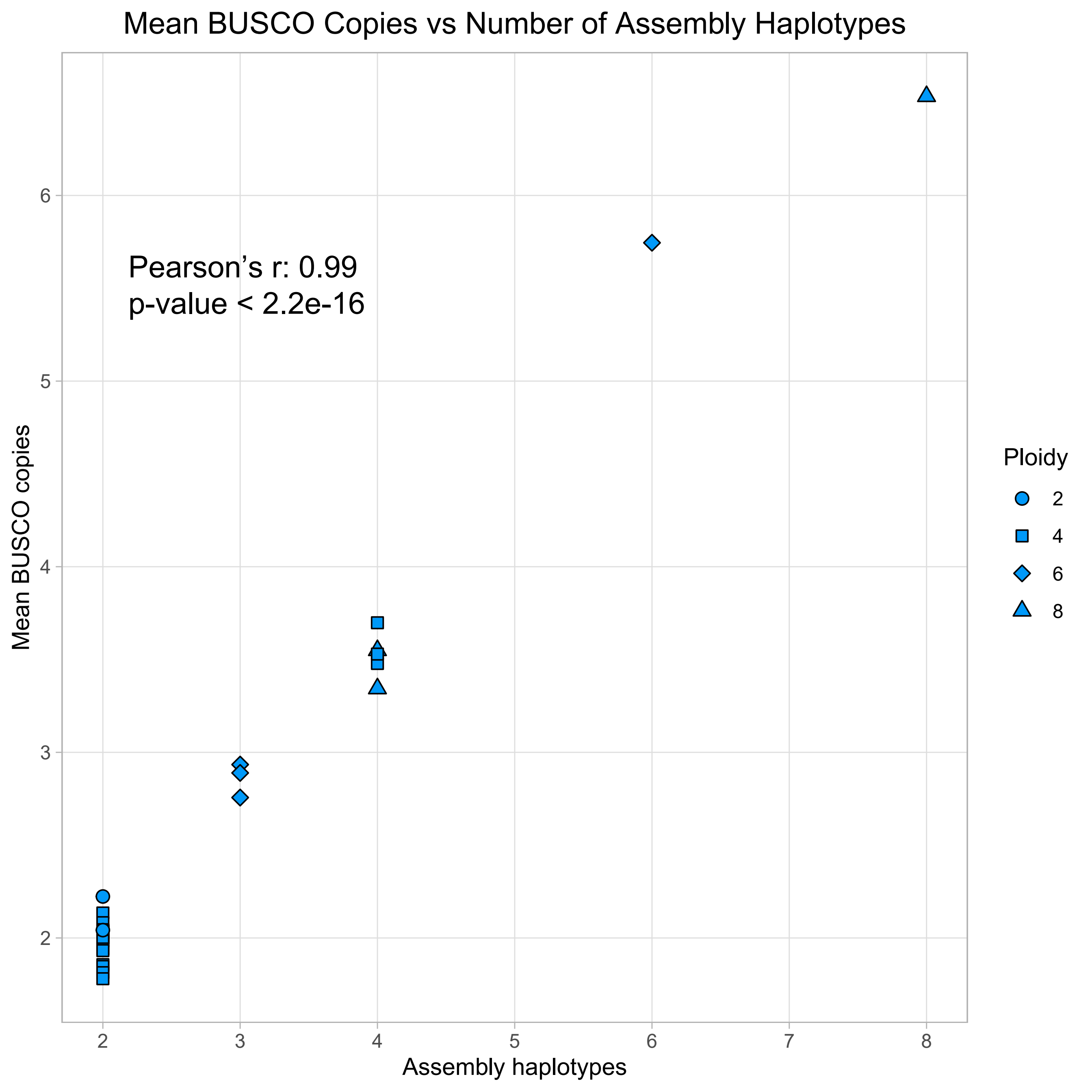

### SFig02.pdf

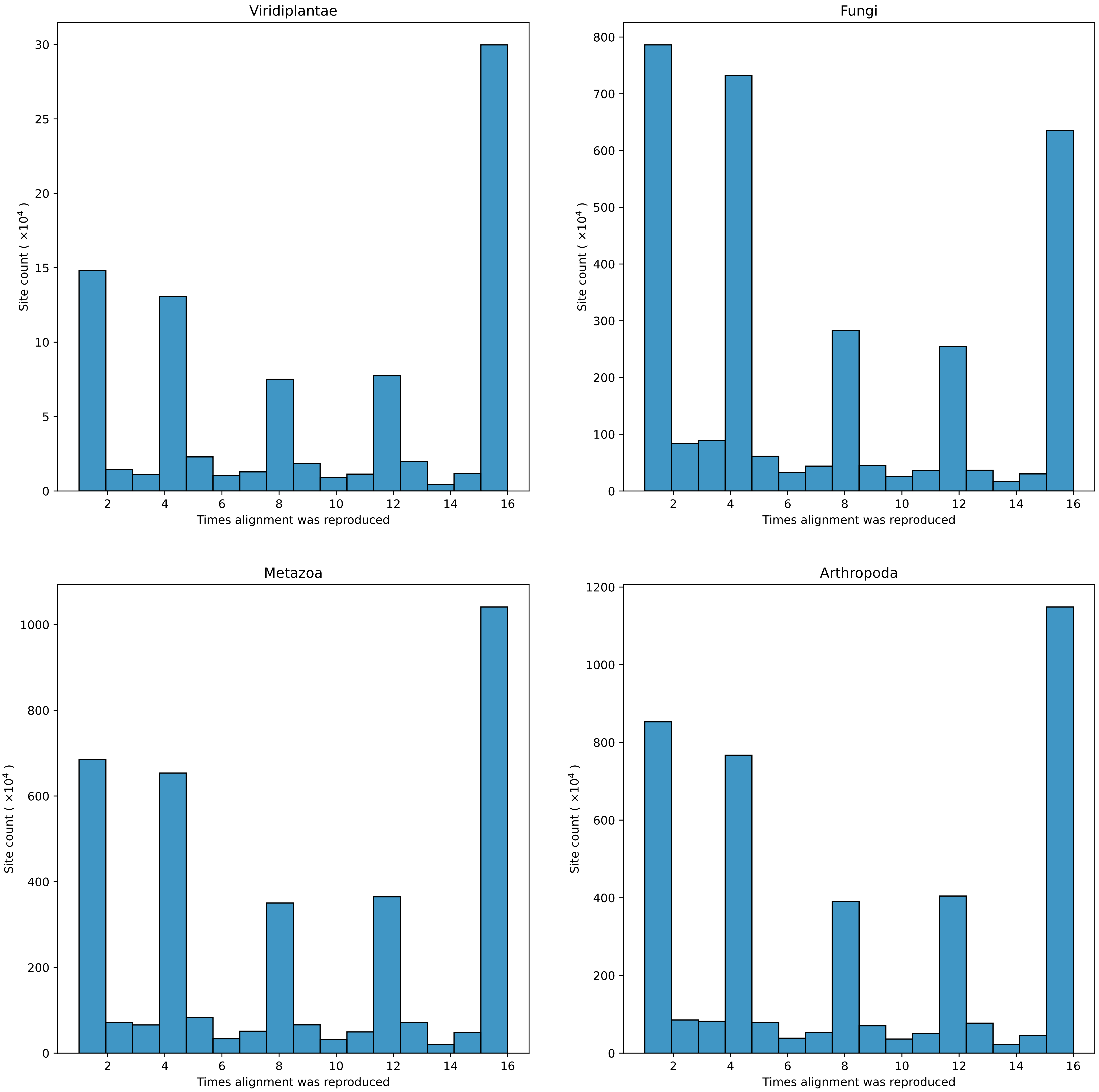

### SFig03.pdf

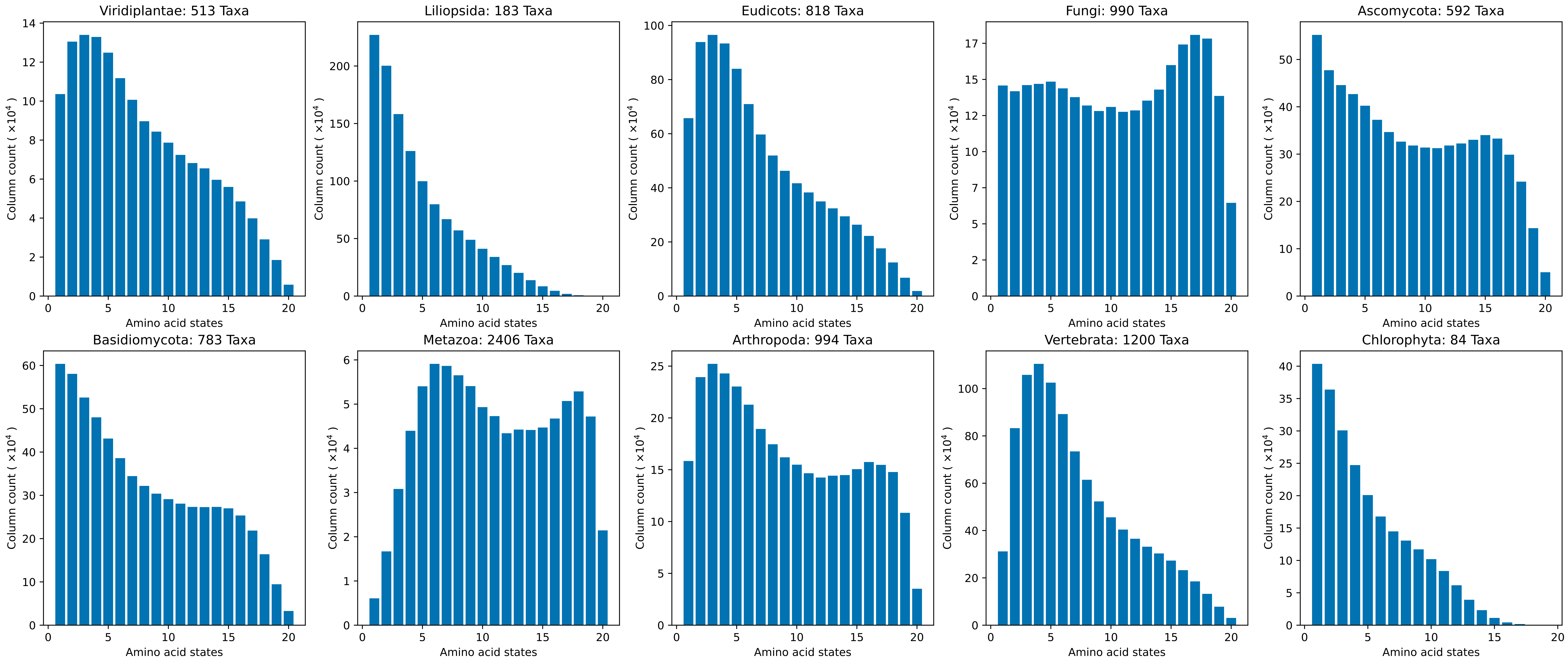

### SFig04.pdf

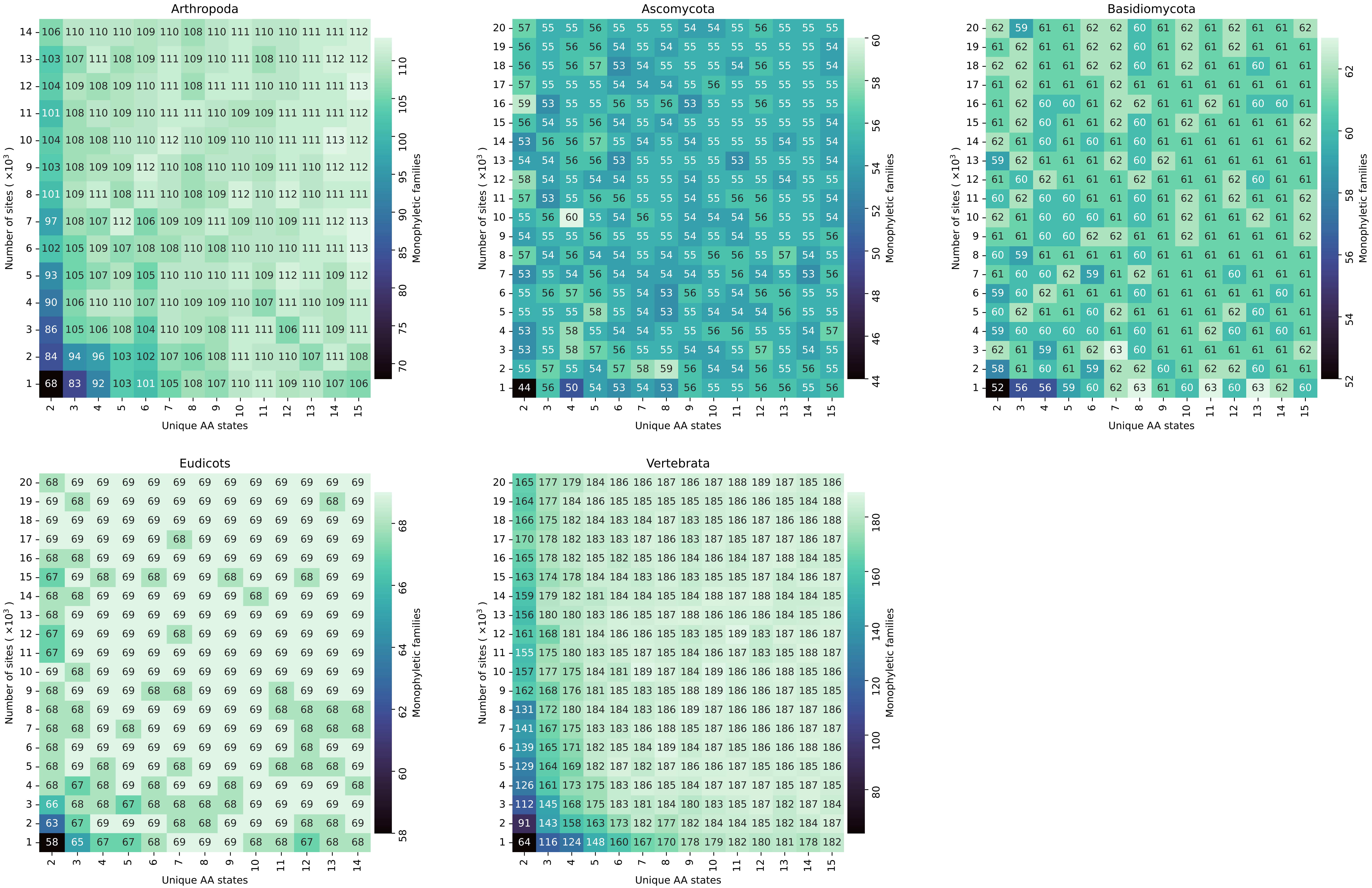

### SFig05.pdf

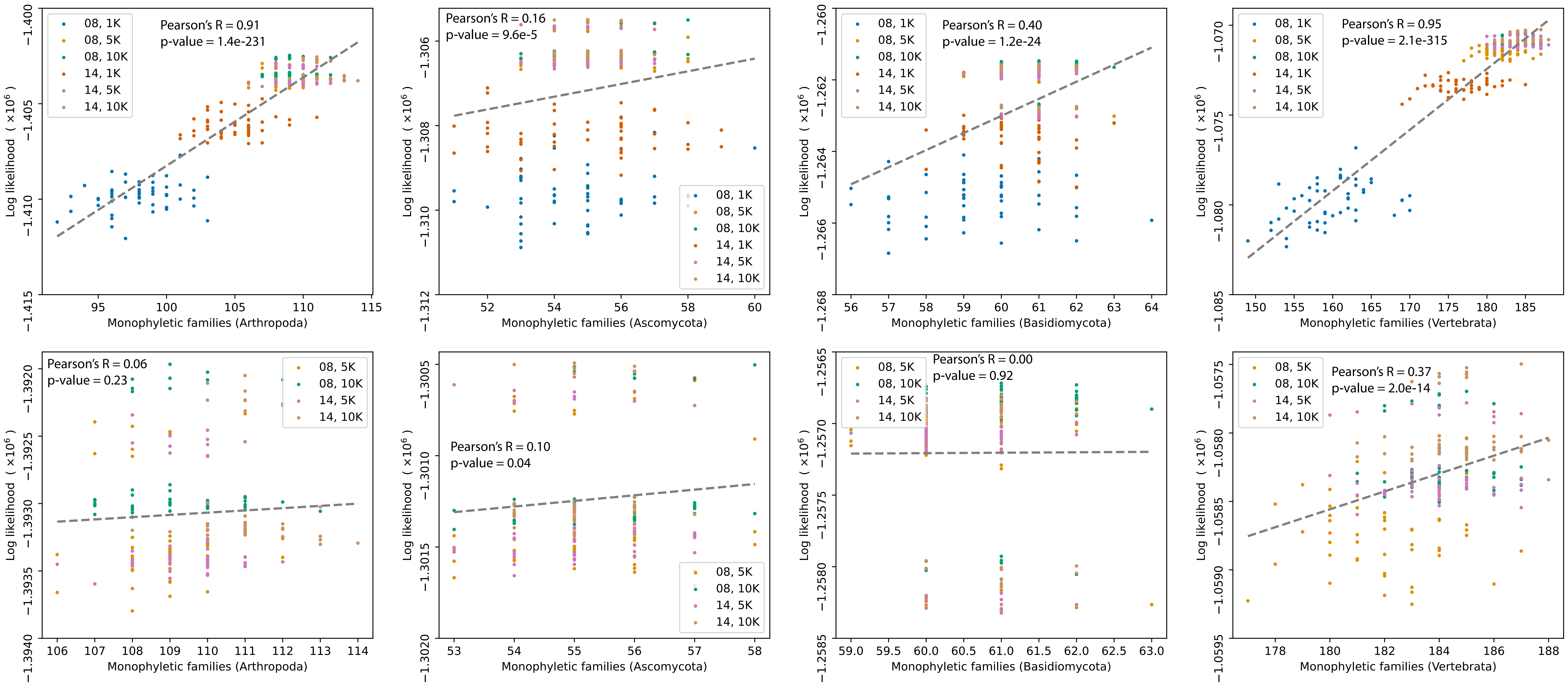

### SFig06.pdf

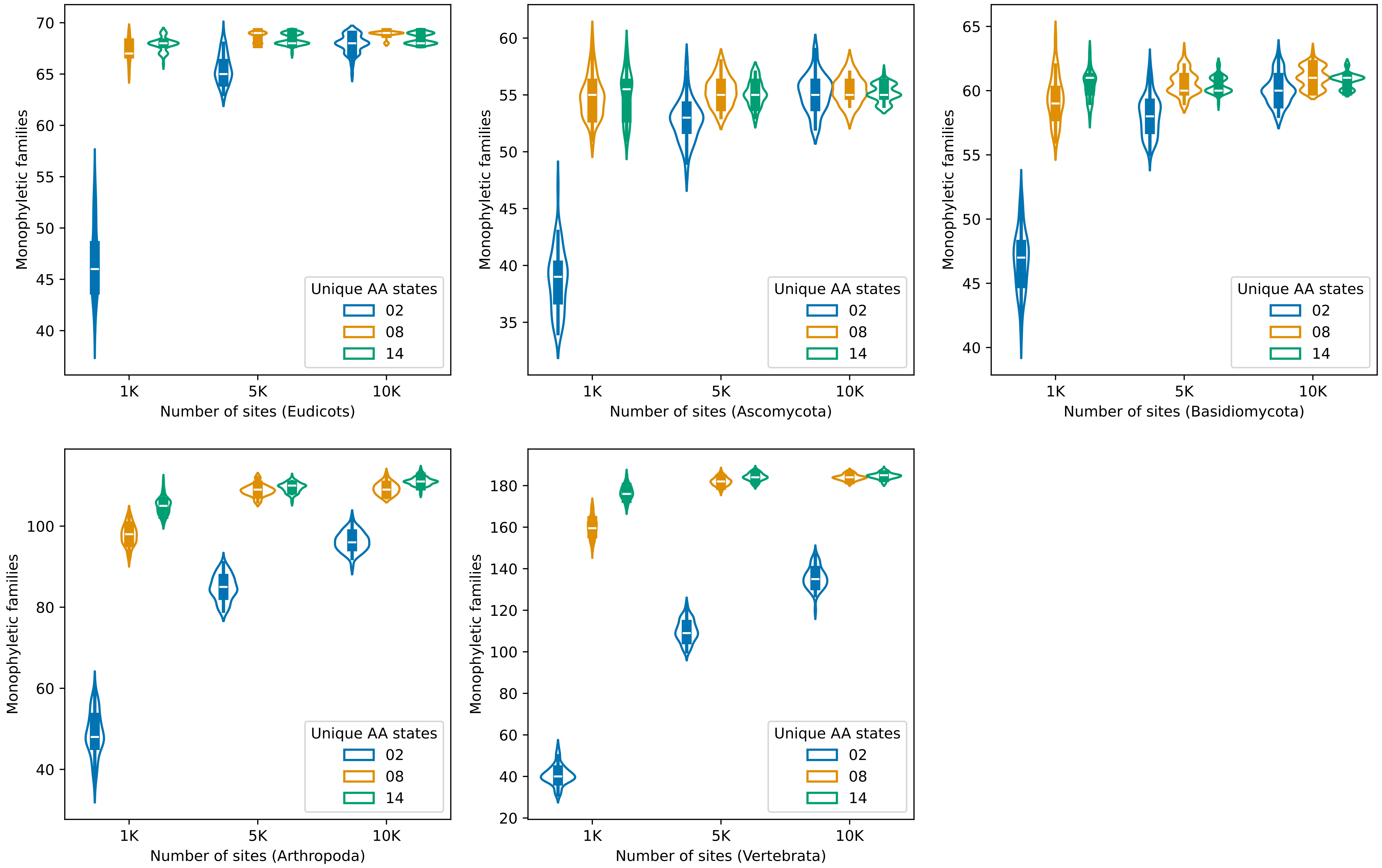

### SFig07.pdf

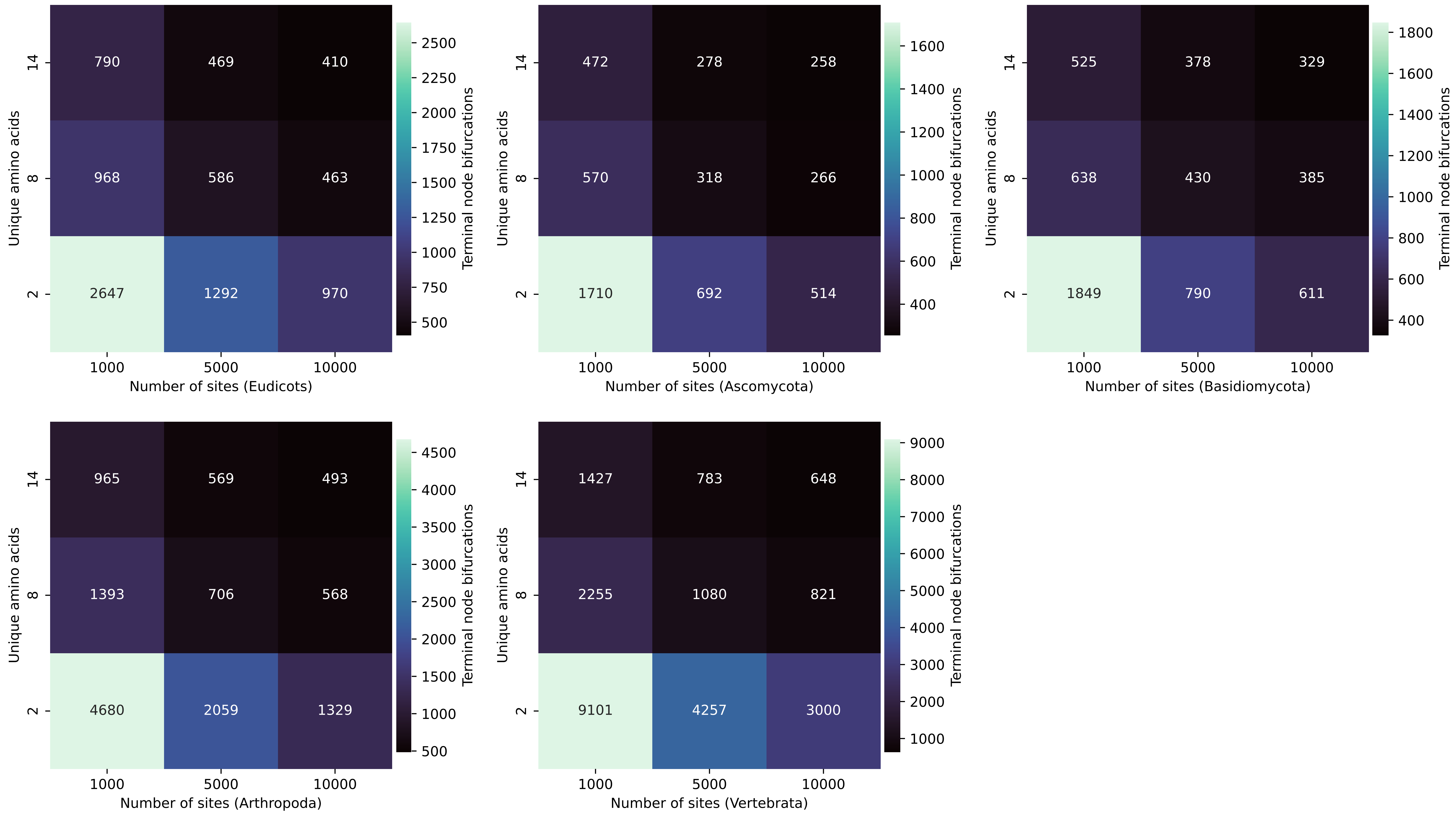

### SFig08.pdf

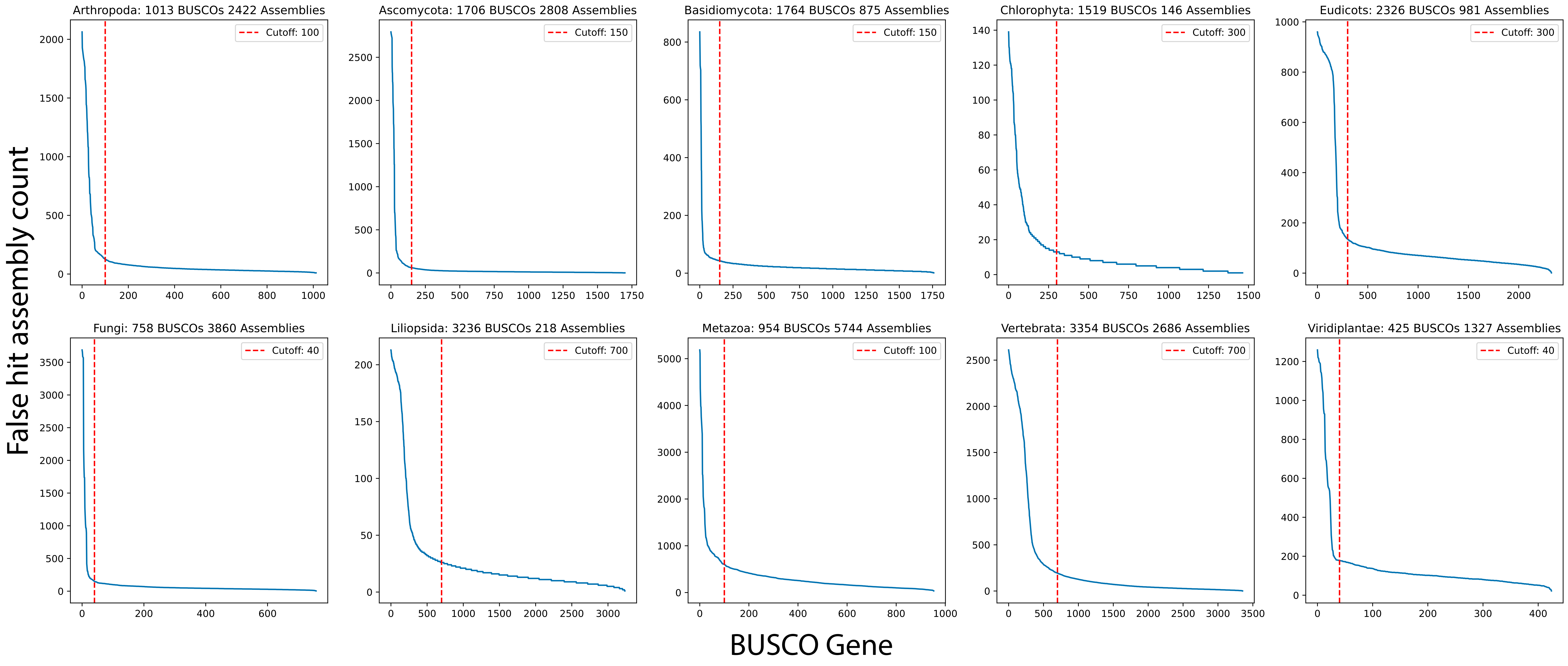
